# Accurate spatial localization of Allen Human Brain Atlas gene expression data for human neuroimaging

**DOI:** 10.1101/2025.06.02.656812

**Authors:** Yohan Yee, Yuhan Liu, Leon French, Violet M. Liu, Alaa Taha, Bradley Karat, Ali R. Khan, Jonathan C. Lau, Yashar Zeighami, Gabriel A. Devenyi, M. Mallar Chakravarty

## Abstract

The Allen Human Brain Atlas has been a tremendously impactful resource in neuroimaging. The usefulness of this resource in neuroimaging arises from spatial coordinates of dissected tissue samples being provided in relation to a Montreal Neurological Institute (MNI)-space standard brain template, thereby allowing for the integration of gene expression and spatially standardized neuroimaging data. In fact, two previous sets of spatial coordinates exist, and surprisingly, the accuracy of these coordinates in placing dissected tissue samples in correct anatomical locations within MNI-space have not been examined. Here, we show that there are significant inaccuracies in the two previous sets of coordinates, and provide a refined set of coordinates as a resource to the neuroscience community. We show (through analyses of meta-analytic data and a re-analysis of real study data) that using previous inaccurate coordinates can result in dramatically different genes being identified, which could compromise further downstream analyses.

## Introduction

The Allen Human Brain Atlas (AHBA) consists of spatially-localized gene expression data obtained from tissue samples acquired from six post-mortem donor brains (*1*). The AHBA has accelerated the development of the field of *imaging-transcriptomics*, a neuroscientific subfield that examines the relationship between brain structure and function and patterns of spatial gene expression (*2–4*). The ability of neuroimaging researchers to use the AHBA stems from the mapping of each individual tissue sample into Montreal Neurological Institute (MNI)-space (*5*), thereby allowing for statistical inference of gene expression properties in relationship to a functional or structural phenotype (represented as a statistical map in MNI-space). Results from these analyses can then be subject to enrichment analyses to determine putative cellular and molecular underpinnings of the observed phenotype (*4*). The popularity and complexity of these analyses have led to the development of various toolboxes and protocols that allow for the manipulation of gene expression data in MNI-space (*4, 6–10*). These toolboxes make use of samples mapped to MNI-space using coordinates originally reported by the Allen Institute or updated MNI coordinates from the ALLENINF software package developed by Gorgolewski et al. (*11*). Importantly, these toolboxes and analyses assume that the reported MNI-space locations are an accurate representation of the sampling site within donor brains. Surprisingly, the accuracy of both these MNI coordinate representations and the sample locations within this context have not been formally examined.

Based on mapping inaccuracies that we observed and are reported below, we developed and report a third improved mapping of AHBA samples to MNI-space, using the MNI ICBM152 nonlinear symmetric 2009c template (“sym09c”, the most recent and recommended multimodal population template defined within MNI-space) as the target template in a multispectral image registration pipeline. Developed at the Cerebral Imaging Centre (CIC; Douglas Research Centre), we refer to this third set of AHBA sample locations as CIC coordinates. We show—both visually and quantitatively—that our CIC coordinate mapping dramatically improves accuracy over previously reported (original and alleninf) coordinate representations. Furthermore, we show that when inputting both real data and a set of statistical maps representative of brain imaging findings (obtained via NeuroQuery (*12*)) into a typical imaging-transcriptomics study, that previously reported coordinates can yield substantially different gene associations compared to CIC coordinates. We provide these improved CIC coordinates to the community for use in future studies in data S1 and at https://github.com/CoBrALab/AllenHumanGeneMNI.

## Results

### Overview of neuroimaging coordinates for tissue samples from the Allen Human Brain Atlas

The Allen Human Brain Atlas (AHBA) consists of genome-wide gene expression data acquired within 3702 tissue samples from six donor brains. Originally, the Allen Institute provided MNI coordinates (referred to hereafter as “original” coordinates) derived from image alignment of donor brains to an MNI-space reference brain, with four out of six brains only being aligned using affine transformations. Soon after, an updated set of coordinates—derived through nonlinear image alignment of all donor brains to an MNI-space template—were made available by Gorgolewski et al. (*11*) as part of the ALLENINF software package (hereafter referred to as “alleninf” coordinates). Though not explicitly stated, the reference brain that is the target of both these image registrations appears to be the MNI152 6th generation nonlinear asymmetric (“nlin6asym”) template (*13*). More recently, we derived an improved set of coordinates for AHBA tissue samples using the newer MNI ICBM152 nonlinear symmetric 2009c template (“CIC” coordinates), as described below (Figure 1A).

**Figure 1:**
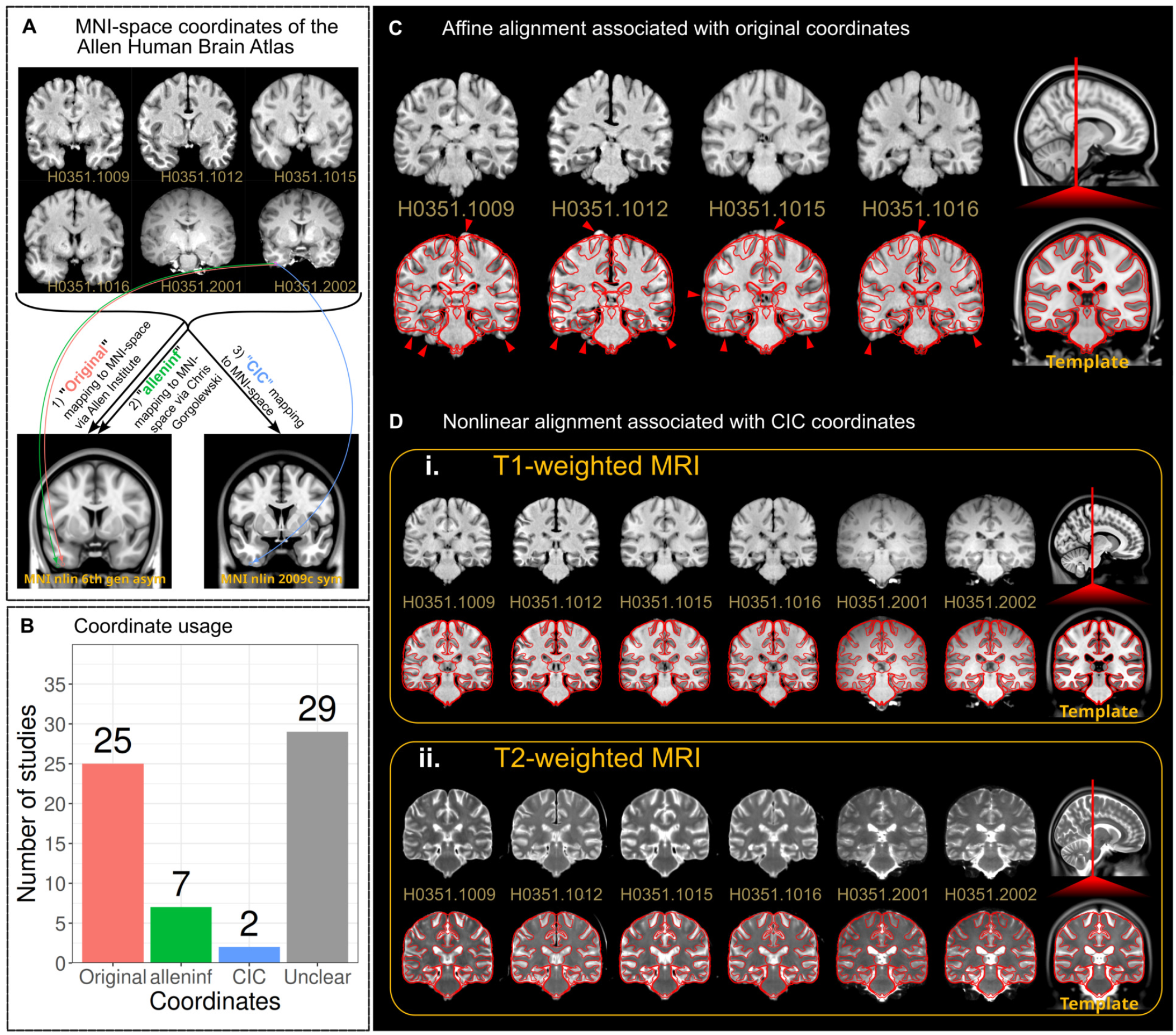
Mapping AHBA data to neuroimaging templates. (**A**) A schematic demonstrating (a coronal slice of) each donor brain MR image, along with the alignment target templates for the three separate coordinate sets. (**B and C**) Coordinates used by a representative set of studies that compute associations between gene expression maps and spatial phenotypes (**B**). Most studies (that explicitly report this information) use original coordinates, which are derived from imprecise alignments of donor MR images to an MNI-space template as shown in (**C**) for a single donor. Specifically, panel (**C**) shows the alignment of subject’s H0351.1009 brain to the MNI152 6th generation nonlinear asymmetric template (determined to only contain an affine component in the transformation) from which original coordinates are derived; contours derived from template intensities are identically projected onto both template and aligned subject images to show alignment failure. (**D**) To overcome coordinate errors, we used an improved image registration framework, which resulted in enhanced alignment between donor and MNI-space template, as shown for all subjects using both (**i**) *T*_1_- and (**ii**) *T*_2_-weighted contrasts.

At present, there is no consensus on which of the two previous sets of coordinates (original or alleninf) best represents true anatomical locations in which gene expression data are obtained. A representative sample of the imaging-transcriptomics literature suggests that approximately three quarters (25/34) of studies (for which the source of coordinates could be identified) rely on original coordinates (Figure 1B and data S2). Importantly, coordinate choice remains an afterthought; almost half of all studies examined (29/63) do not report which set of AHBA coordinates were used. Inaccuracies in either set of coordinates can result in systemic biases due to inaccurate image registration of donor brains to the template (Figure 1C). This is particularly an issue in the original set of coordinates that are derived from affine-only alignment for four out of the six donor brains (Supplementary Text), and for which no quality control on anatomical mapping to MNI-space is specified (*1*). These observations motivated our derivation of the new CIC coordinates for AHBA tissue samples. We formally compare these new coordinates to the previous two sets and validate the anatomical accuracy of each set of coordinates through a variety of independent tests, and subsequently ascertain the impact of coordinate choice on neuroimaging-transcriptomics studies.

### Registration to the MNI ICBM152 nonlinear symmetric 2009c template

Based on the observations above, we sought to re-align all magnetic resonance images (MRI) of the donor brains to the MNI ICBM152 nonlinear symmetric 2009c (“sym09c”) template, a nonlinear average of 152 brains derived from a normative young adult population used to define MNI-space at a 1 mm isotropic resolution (*5*). Briefly, each set of donor brain images (*T*_1_-weighted and *T*_2_-weighted) were bias field-corrected (*14, 15*). Masked donor brains were then individually aligned to the template using ANTS (*16*) using both affine and nonlinear stages (*17*). Specifically, each donor’s *T*_1_- and *T*_2_-weighted brain images were aligned to their corresponding *T*_1_- and *T*_2_-weighted sym09c target templates using a joint optimization with equal weighting of metric loss function to produce a single donor-template transformation. This strategy leverages complimentary image contrast information across both images as a means of obtaining a more accurate nonlinear mapping between donors and the sym09c template. In the case of a single donor (H0351.1016) in which a sufficiently large deviation of cerebellar position was observed (figure S1), the cerebellum was masked out and aligned separately.

Donor MR images aligned well to the MNI-space template (Figure 1D), and passed visual quality assessment checks. Tissue coordinates in MRI space—provided by the Allen Institute—were subsequently transformed to MNI-space using transformations obtained from the aforementioned image registrations, resulting in the CIC coordinates which we provide and examine here. Transformations used to generate CIC coordinates achieve donor-template alignment through roughly equal amounts of nonlinear deformations of donor brains as compared to transformations generating alleninf coordinates, but use greater deformations than transformations generating original coordinates (figure S2). CIC coordinates, along with code used to generate them through image registration of donor brains, are freely available on GitHub: https://github.com/CoBrALab/AllenHumanGeneMNI. CIC coordinates are also included as data S1.

To enable quantitative comparisons between all three coordinate representations of sample locations and determine their accuracies, sample coordinates were further transformed to sym09c and nlin6asym templates via transformations obtained from an alignment between the *T*_1_-weighted nlin6asym and sym09c templates. Specifically, original and alleninf coordinate representations were further mapped to the sym09c template using this aforementioned template-to-template transformation, and CIC coordinates were transformed to the nlin6asym template using the inverse of the aforementioned transformation. Unless otherwise stated, all results are presented using coordinate representations in relation to the sym09c template. To ensure that downstream analyses are not impacted by this additional transformation, we repeated all major analyses using coordinate representations relative to the nlin6asym template, where only CIC coordinate-based locations are impacted by this additional transformation.

### Comparison of CIC coordinates to previous sources

Visual examination of the samples mapped to MNI-space using CIC coordinates suggested strong correspondence between true neuroanatomical locations of samples within subject brains and locations relative to neuroanatomy defined by the sym09c template. Comparing CIC coordinates to original and alleninf coordinates (see Figure 2A for an illustrative example from one donor), we found that clear misalignment to MNI-space was visually apparent for many samples in the original and alleninf coordinate sets, but not with CIC coordinates. In general, differences between coordinate sets were not random, but tended to be pronounced within certain structures or regions of space, likely reflecting regularization within image registration (Figure 2B). In the case of donor H0351.1009, differences between CIC coordinates and original coordinates were particularly evident in the hippocampus.

**Figure 2:**
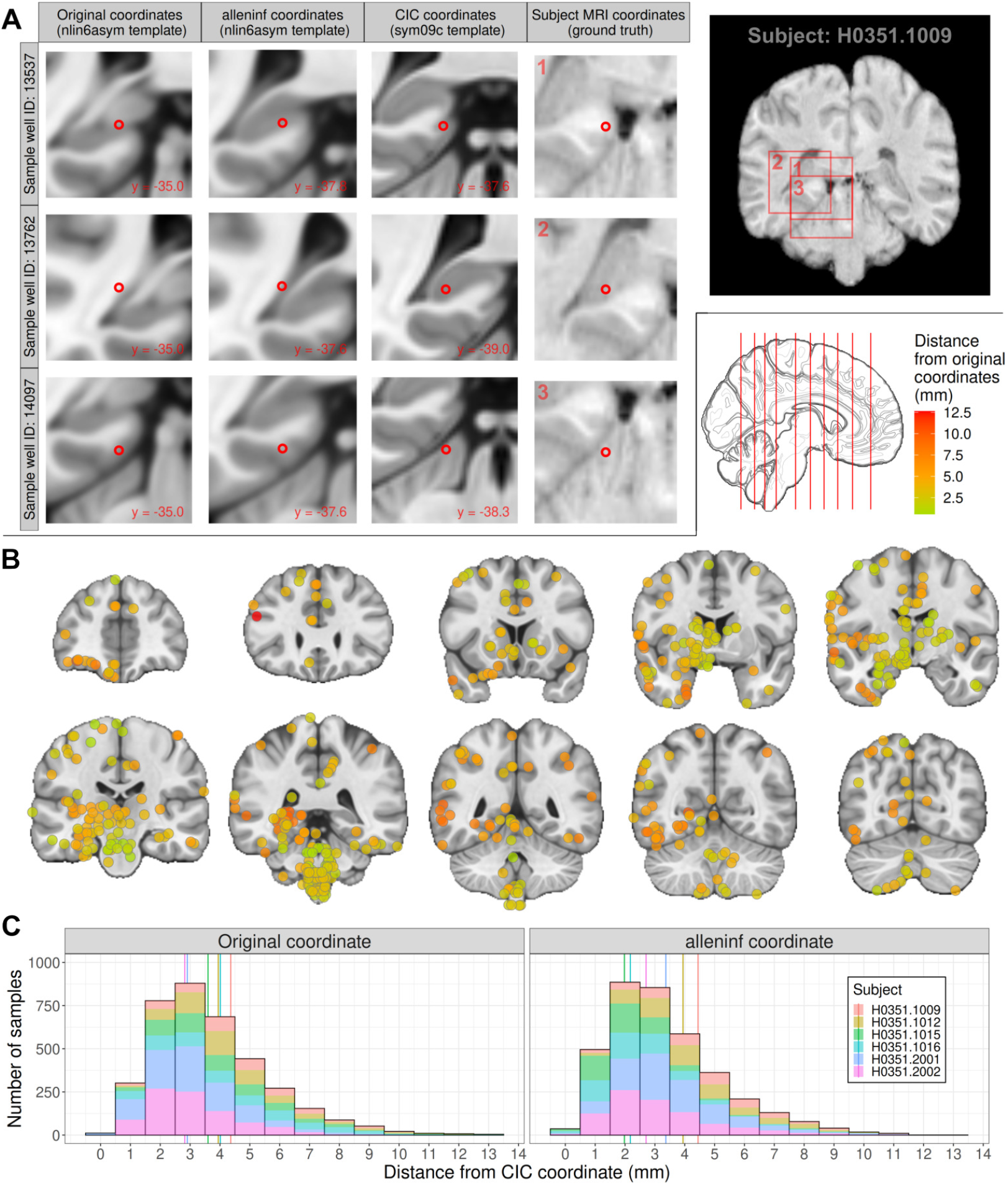
Comparison of sample coordinate sets. (**A**) Comparison of sample locations in subject and MNI-space. Shown are three example samples (rows) obtained from donor H0351.1009 from within a single slice (anterior-posterior MRI world coordinate: -18.0). Samples are derived from the left parahippocampal gyrus, lateral bank of gyrus (top row), left dentate gyrus (middle row), and left cerebellar lobule IV, lateral hemisphere (bottom row), as annotated by an expert neuroanatomist at dissection time. Samples are plotted in MNI-space using coordinates provided by each of the three coordinate representations (original, alleninf, CIC; first three columns), and are displayed over their respective *T*_1_-weighted template (target of registration; either the MNI ICBM152 sixth generation nonlinear asymmetric template for original and alleninf coordinates or the symmetric 2009c template for CIC coordinates). The final column displays samples in subject space, using the subject MRI voxel coordinates reported by the Allen Institute (which we take as ground truth) and plotted over the subject *T*_1_-weighted scan. (**B**) Samples from all subjects are plotted on coronal slices using CIC coordinates. Additionally, samples are coloured to show the total Euclidean distance that points are shifted under CIC coordinates, relative to the original coordinates. Only 579 samples that lie within the slice planes are shown (15.6% of all AHBA samples). (**C**) The distribution of these coordinate shifts are quantified across all samples (including those out-of-plane) and all subjects. Distances shifted under CIC coordinates relative to both original (left) and alleninf (right) coordinates are visualized as histograms, and vertical lines represent median distance between CIC and original and CIC and alleninf for each subject.

Compared to original and alleninf coordinates, the median distances to CIC coordinates were at least 2 mm in all except one donor brain (Figure 2C, table S1), and the distribution of these distances between coordinates formed a long tail, indicating that many samples are placed in very different locations depending on the coordinate representation used. For 59 samples, differences between MNI-space coordinate locations are greater than 1 cm when comparing at least one pair of coordinate representations.

### CIC coordinates yield template intensity profiles that are more similar to donor brain scans

While pulse sequence and image processing will result in different intensity scales between the sym09c template and donor MRI, image registration should, by definition, align voxels of similar relative intensity together (*18*). We propose that the accurate placement of coordinates in MNI-space should result in monotonic relationships between the intensity values of aligned subject and template images, as measured across sets of samples. To test this, we quantitatively assessed the accuracy of all three coordinate representations by correlating the template intensity values at the sample locations transformed to MNI-space to intensity values of donor brain MRI scans under ground truth MRI coordinates. We observed that CIC coordinates generally produced dramatically higher Spearman correlations across *T*_1_- and *T*_2_-weighted images as compared to original and alleninf coordinates (table S2). Figure 3A shows these intensity correlations for one subject (H0351.1009) and Figure 3B compares correlations obtained from original, alleninf, and CIC coordinates for all subjects. Our results were robust to the choice of template on which analyses were carried out (Supplementary Text) and were also observed within coarse subdivisions of the brain (cortex, subcortex, brainstem, cerebellum; figure S3) from which the samples were dissected.

**Figure 3:**
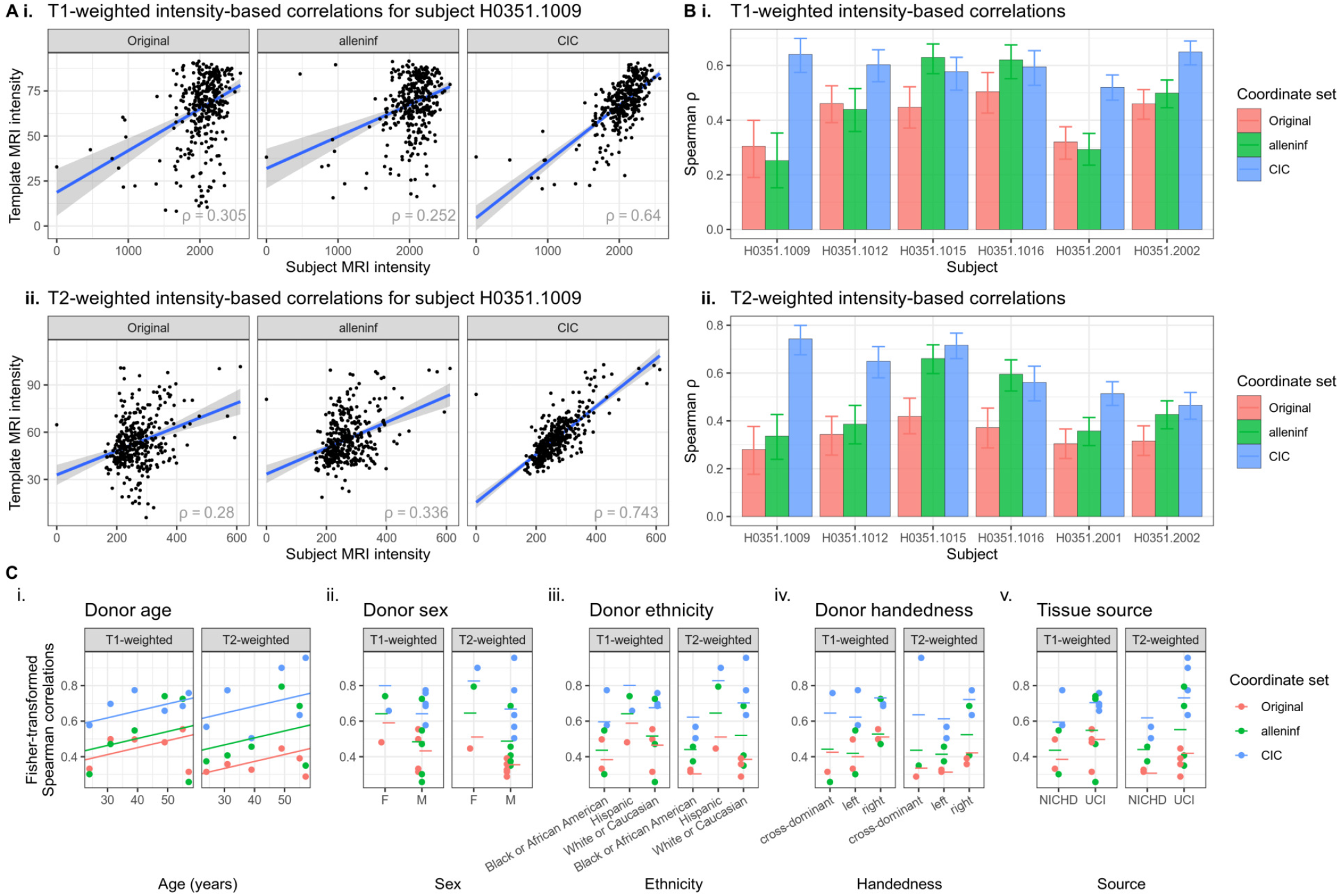
Intensity-based validation of registration quality. (**A**) Spearman correlations between the image intensities of subject scans and template images, for (**i**) *T*_1_-weighted and (**ii**) *T*_2_-weighted contrasts at reported sample locations for donor H0351.1009. Each point is a sample, and correlations are computed separately for each coordinate set. Higher (Spearman) correlations imply stronger monotonic relationships between the intensities of subject and template scans (i.e. lighter voxels mapped to lighter voxels, and vice versa), and therefore better registration quality for the given subject. (**B**) Spearman correlations are further shown for all other donors, using both (**i**) *T*_1_-weighted and (**ii**) *T*_2_-weighted contrasts, and error bars denote the 95% range of Spearman correlations obtained through bootstrapping. (**C**) The relationship between registration quality (as quantified by the Spearman correlation coefficient) and donor characteristics are further explored for (**i**) age (years), (**ii**) sex, (**iii**) ethnicity, (**iv**) handedness, and (**v**) tissue source (NICHD: University of Maryland, Baltimore; UCI: University of California, Irvine, imaged *ex cranio)*. Lines show linear mixed effects model predictions of (variance-stabilized) Spearman correlations when using each characteristic as a fixed effect shared across all coordinate sets and contrasts, but random intercepts (Supplementary Text).

### CIC coordinates result in fewer misclassified samples

Many studies that use the AHBA analyze data at the regional level and compute average gene expression under regions of interest (ROIs) based on groups of samples that lie within those structures (*6, 19*). Given that we observed an improvement in sample coordinate mapping accuracy, we sought to determine how much of a difference CIC coordinates can make in such ROI-based studies. We used the Allen Human Reference Atlas (*20*) (which consists of volumetric segmentations of the MNI ICBM152 nonlinear symmetric 2009b template; referred to here as the “AHRA”) and further transformed this atlas onto the sym09c and nlin6asym templates to define ROIs for the purpose of this study). We compared atlas-based annotations from the AHRA to ground truth annotations provided within the AHBA. For each sample, we determined the AHRA regions that they were located in according to the original, alleninf, and CIC coordinate representations.

First, we examined neuroanatomical concordance by determining the structure that sample coordinates fall within according to the AHRA definitions of regions within the sym09c template (data S3), and based on these regions, further classifying samples as falling within in one of five coarse structures (cortex, subcortex, brainstem, cerebellum, and white matter and ventricles) or out-of-brain (data S4 and data S5). Here, we use terminology that is consistent with the AHBA ontology; coarse structures are derived by traversing up the tree of regions. Specifically, samples dissected from the choroid plexus of the lateral ventricle are described as being in the “white matter and ventricles” coarse structure, since one arrives at the “ventricles” node when traversing up two levels of the AHBA tree from the “choroid plexus of the lateral ventricle” node. Of the 3702 samples in the AHBA, only 5 are from the ventricles (all annotated as being dissected from the choroid plexus of the lateral ventricle), and 15 from the white matter (13 corpus callosum, 2 from the cingulum bundle).

Across all three coordinate representations, a major mode of location inaccuracy arose from many samples being placed in white matter/ventricles or outside the brain, despite only 20 being truly sampled from the white matter/ventricles. In contrast, CIC coordinates provided a more accurate description from which samples were truly derived (table S3), with 30-40% fewer grey matter samples being placed outside grey matter (769 samples) as compared to alleninf (1095 samples) and original (1283 samples) coordinates respectively (Figure 4A-B and table S4). A particularly striking example of coordinate inaccuracy is that 25 samples dissected from the cerebellum are classified as being in the cortex when using original coordinates, as compared to 3 when using alleninf coordinates, and 3 when using CIC coordinates. For each of the five coarse structures from where samples were obtained, we computed the proportion of misclassified samples (i.e. the false negative rate; FNR), and found that for all except white matter and ventricles (which corresponded to by far the fewest samples), the FNR was lowest when using CIC coordinates (Figure 4C and table S5). The ontologies used by the AHBA and ARHA both divide the cortex into the same lobes (data S6), allowing for samples’ coordinate-based regions to also be compared to expert annotations at a finer level (data S7). For cortical samples placed within lobes based on MNI coordinates, we found that CIC coordinates placed samples in the lobe from which they were dissected most often (Figure 4D and tables S6-8).

**Figure 4:**
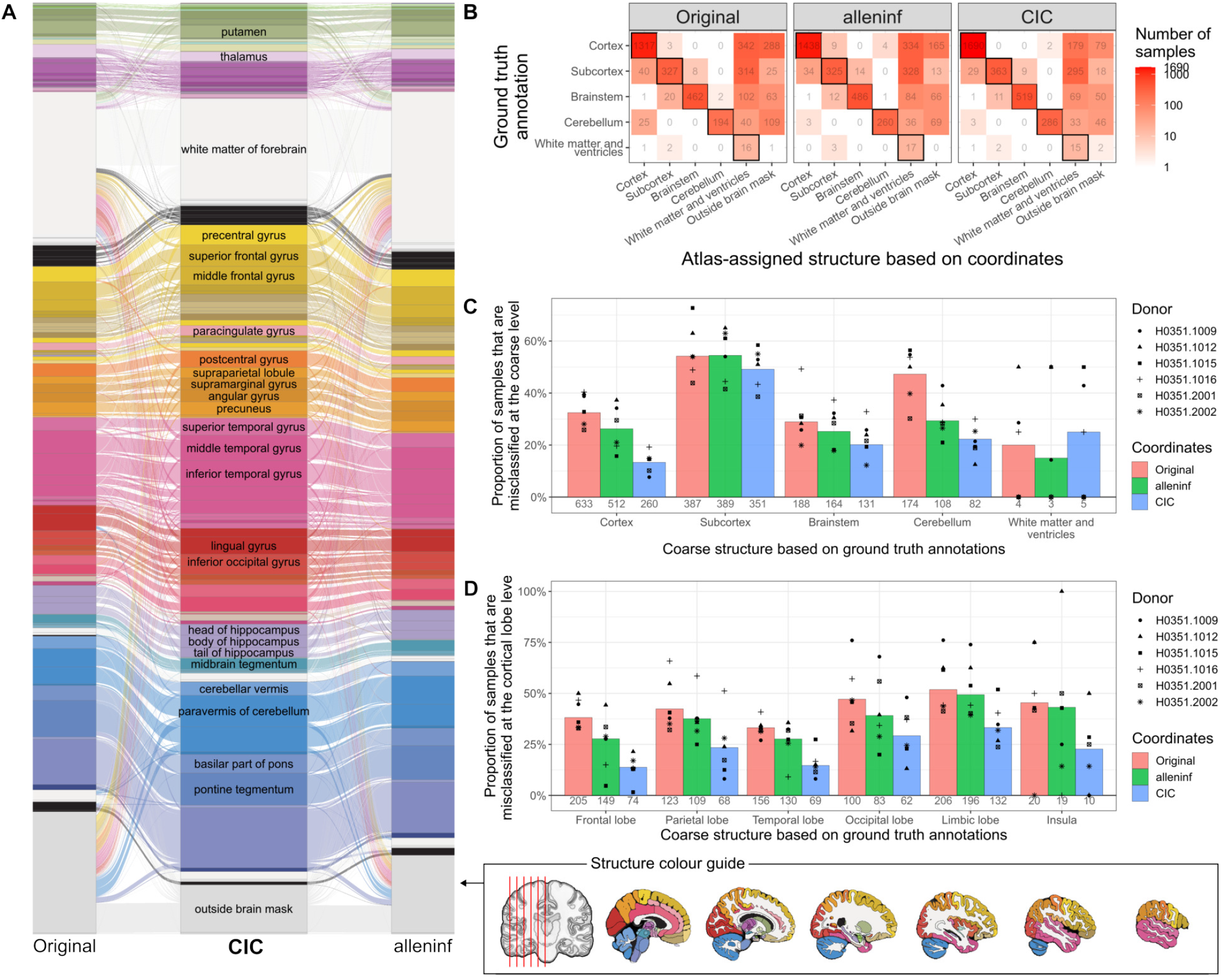
Assignment of samples to structures based on coordinate locations within the Allen Institute’s Human Reference Atlas (AHRA) relative to the sym09c template. (**A**) Comparison of sample assignment within the Allen Institute’s Human Reference Atlas (AHRA). Shifts in sample assignment based on original, CIC, and alleninf coordinates relative to the AHRA are displayed as a Sankey diagram. Many of the samples reported to be in the white matter or outside the brain based on the original (left) or alleninf (right) coordinates fall within cortical structures when using CIC coordinates. (**B**) Assignment of samples to five large-scale (coarse) structures and comparison to ground truth. Confusion matrices for each coordinate set show the number of samples assigned to one of 5 coarse structures (y-axis) based on expert annotations, along with coordinate-based assignment based on sample location within the AHRA. In all coordinate sets, significant mislabelling in which grey matter samples are either shifted to white matter or outside the brain occurs, but true positive rates (TPR) remain highest for CIC coordinates. (**C**) Mislabeling above is quantified as the false negative rate (i.e. proportion of samples misclassified, 1-TPR). The false negative rate (FNR) is lowest for CIC coordinates in all coarse structures except “white matter and ventricles”, which consist of vastly fewer samples compared to the other four coarse structures. Each point is a subject, and bars show the false negative rate after pooling samples across all subjects. (**D**) Similar to (**C**), the FNR is computed based on the coordinate-based placement of cortical samples within cortical lobe structures.

The FNR quantifies the proportion of samples that are mislabeled given a true structure label. In practice, it is often helpful to know the probability that one is wrong given the observation of a sample coordinate falling within a structure, i.e. the false discovery rate. Here too, in examining coordinate placement within coarse structures and cortical coordinate placement within lobes, we found CIC coordinates to be most optimal (Supplementary Text).

The above analyses examining FNR and FDR were repeated using coordinate representations relative to the nlin6asym template (see Supplementary Text), and nearly identical results were obtained.

### CIC coordinates remain most accurate after remapping mislocated samples to grey matter

The issue of many AHBA samples being located outside grey matter has been recognized in the literature (*4, 8, 21*), and a common approach to rectify this is to simply assign them to their nearest structure. For instance, Arnatkevičiūtė et al. (*4*) recommend applying this procedure to samples within 2 mm of a region, to avoid inaccurate mappings due to large shifts. Given the large number of samples found outside grey matter (including when using CIC coordinates), along with the aforementioned preprocessing recommendations, we repeated previous analyses (Figure 5A) after re-mapping each sample (except the 20 non-grey matter samples) to the nearest grey matter voxel based on Euclidean distance (Figure 5B(i)) independently for each of the three coordinate sets (data S8-9). Non-cortical structures gained the most samples due to remapping across all three coordinate sets, namely the thalamus, medulla oblongata, cerebellum, putamen, and caudate (table S9). The spatial distribution of increase in sample numbers was generally consistent across all three coordinate sets, but due to fewer samples being placed outside grey matter by CIC coordinates, fewer samples had to be remapped when using CIC coordinates (Figure 5B(ii)).

**Figure 5:**
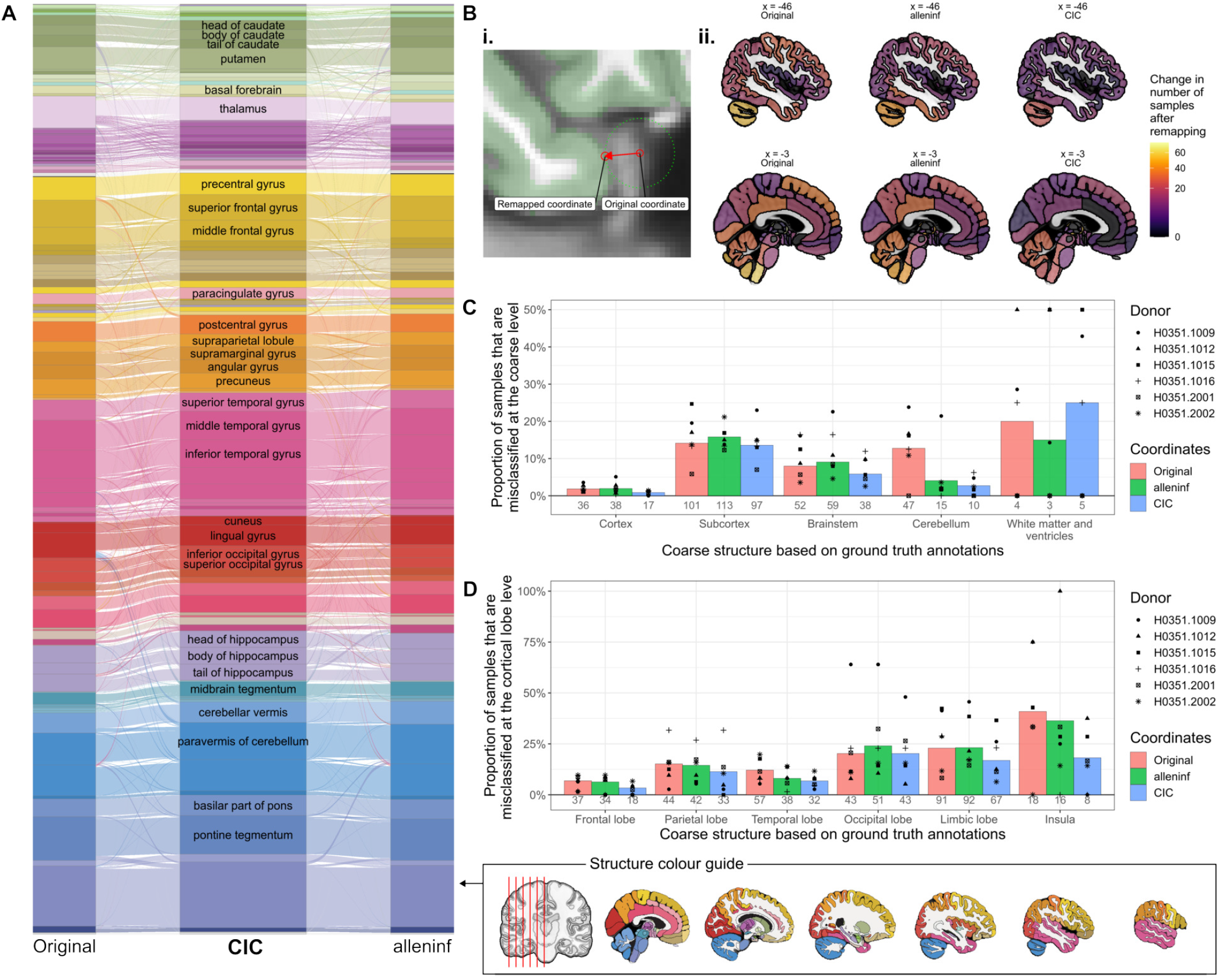
Assignment of samples to structures based on grey matter-remapped coordinate locations within the Allen Institute’s Human Reference Atlas (AHRA). (**A**) Comparison of sample assignment within the Allen Institute’s Human Reference Atlas (AHRA) after remapping coordinate-based locations to their respective samples’ closest grey matter voxel. Shifts in sample assignment based on original, CIC, and alleninf coordinates relative to the AHRA are displayed as a Sankey diagram. After remapping sample locations to grey matter, structure disagreements persist between CIC coordinate locations and other coordinate representations. (**B**) (**i**) Demonstration of a clearly mislocated sample (initially placed out of the brain) being remapped to the closest grey matter voxel. (**ii**) The numbers of samples that neuroanatomical structures gain after remapping are visualized as colours. Brainstem and cerebellar structures tend to gain the most samples after remapping for all coordinate representations, although this is most pronounced for original and alleninf coordinates. (**C**) Mislabeling above is quantified as the false negative rate (i.e. proportion of samples misclassified, 1-TPR). As expected, the FNR is generally lower after remapping MNI coordinates to grey matter. Notably, the false negative rate (FNR) remains lowest for CIC coordinates in all coarse structures except “white matter and ventricles”, which consist of vastly fewer samples compared to the other four coarse structures. Each point is a subject, and bars show the false negative rate after pooling samples across all subjects. (**D**) Similar to (**C**), the FNR is computed based on the coordinate-based placement of cortical samples within cortical lobe structures. FNR is generally lower after remapping MNI coordinates to grey matter, and CIC coordinates continue to have the lowest FNR.

We found that for samples which were remapped, the distribution of distances describing their shifts formed a long tail, with some samples being moved as much as 8 mm (figure S4 and table S10). If the guideline by Arnatkevičiūtė et al. (*4*) of excluding samples requiring a shift of over 2 mm were to be followed, then the lowest bound on the number of unusable samples is most optimal when using CIC coordinates (221 discarded samples; ∼6% of total) as compared to original (412 discarded samples; ∼11% of total) and alleninf (293 discarded samples; ∼8% of total) coordinates. In cortical samples, the lowest bound on the number of samples to exclude when using CIC coordinates continues to be most optimal (52 discarded samples, ∼2.7% of cortical samples) as compared to original (181 discarded samples, ∼9.3% of cortical samples) and alleninf (102 discarded samples, ∼5.2% of cortical samples) coordinates, a result that also holds in all other coarse substructures (subcortex, brainstem, cerebellum) and in five out of six subjects.

We confirmed that previously-described results did not change when using grey matter-remapped coordinates: intensity-based Spearman correlations remained highest when using remapped CIC coordinates (figure S5), and coarse structure misclassification based on spatial coordinates remained lowest for the remapped CIC coordinates, as compared to previous coordinate sets (tables S11-12). For cortical samples, CIC coordinates resulted in more than a 50% reduction in misclassified samples (17 samples), as compared to original (36 misclassified samples) and alleninf (38 misclassified samples) coordinates (Figure 5C). Using remapped coordinates that placed more samples in the cortex, we further examined concordance between cortical samples’ coordinate-based locations and expert annotations at a more granular level as before. Here too, we found that CIC coordinates resulted in fewer misclassified samples (Figure 5D).

### CIC coordinates provide expression patterns most consistent with their spatial location

While we have shown that CIC coordinates are likely a better reflection of the true locations of tissue samples, does the choice of coordinates make tangible differences in spatial gene expression patterns summarized over regions? To determine this, we first focused only on samples that were placed in different AHRA-defined regions when comparing (i) original and CIC coordinates, and (ii) alleninf and CIC coordinates. Given that regions are partly defined based on cytoarchitecture derived from histology (*20*), we expect that a sample’s gene expression profile (expression across all genes) will more likely be similar to all other samples truly located within the same region, as compared to samples in other regions. Thus, through gene expression, we can infer the region from which a sample is most likely derived from, and test which set of coordinates places the sample in this region. As demonstrated by a single sample which was placed in different regions when using original, alleninf, or CIC coordinates (Figure 6A), we examined the similarity (Pearson correlation) between each sample’s reported gene expression and the mean gene expression profiles of all other samples that share the same region label for each of the different coordinate sets (data S10). Comparing both original and CIC coordinates and alleninf and CIC coordinates when assigned region labels differ (data S11), we found a general increase in gene expression similarity when using CIC coordinates (Figure 6B), with differences in expression similarity being more pronounced when compared to original coordinates. The most significant and robust effects were seen for cerebellum, cortex, and subcortex, as opposed to the brainstem, and appeared to be driven by specific subjects (H0351.1009, H0351.1012, H0351.1015, H0351.1016 in the comparison with original coordinates; H0351.1009, H0351.1012, H0351.2001 in the comparison with alleninf coordinates; figure S6). CIC coordinates remain optimal when using grey matter-remapped coordinates (figure S7 and data S12-13).

**Figure 6:**
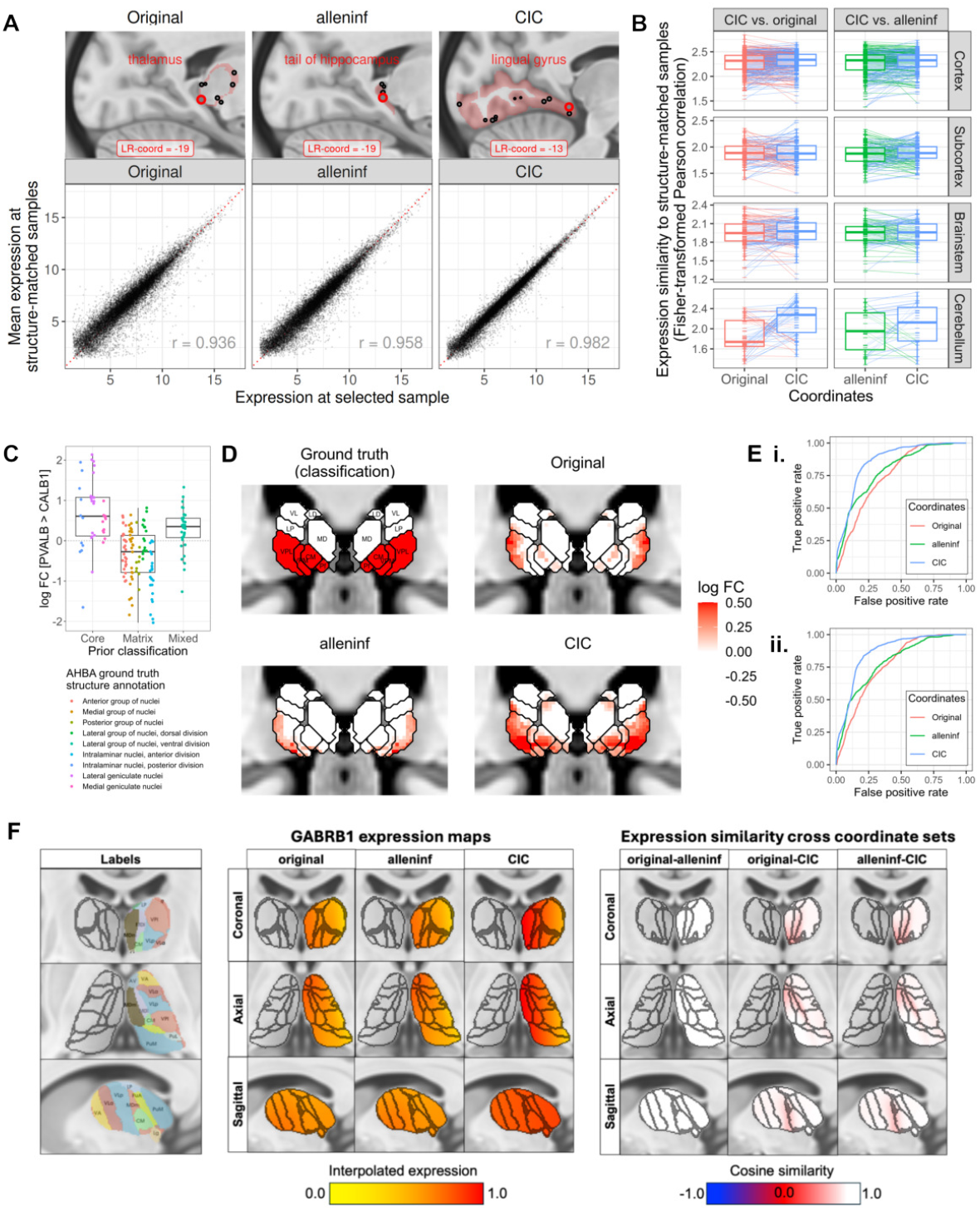
Effects of coordinate set choice on regional and voxelwise measures of gene expression. (**A**) an example describing a test of which region a tissue sample belongs to, based on the similarity between its gene expression profile and the expression profiles of other nearby samples. Top row: for a given sample (red circle) and a coordinate set, it is placed into a region defined by the AHRA based on its coordinates, and other samples within that region are identified (note: other samples are only shown if they are on or near the displayed sagittal plane). Bottom row: the gene expression profile of the given sample is then compared to the average gene expression profile of all other samples within the region it is placed in (each point in the plot represents a gene), and is quantified using Pearson’s correlation coefficient. A higher correlation implies a greater likelihood that the sample is part of the region in which it is placed. (**B**) Fisher-transformed correlation coefficients are compared between original and CIC (left column) and alleninf and CIC (right column) coordinate sets, with differences highlighted by lines joining the same samples. (**C**) Effects of coordinate choice on spatial gene expression within the subcortex are further assessed through an interpolation task. Expert annotations are used to verify that the ratio of parvalbumin to calbindin expression *(PVALB*/*CALB1)* are higher in samples derived from the posterior group of intralaminar nuclei, dorsal lateral geniculate nucleus, and medial geniculate nuclei, as compared to other thalamic samples, as suggested by the core-matrix framework. Each point is a thalamic tissue sample; points are coloured by the thalamic nuclear complex from which it is dissected (within the AHBA ontology). (**D**) Parvalbumin *(PVALB)* and calbindin *(CALB1)* expression are then interpolated voxelwise throughout the thalamus using all 175 AHBA samples derived from the thalamus, and the ratio of *PVALB*/*CALB1* interpolated expression—a transcriptomic marker of core nuclei—are compared to the spatial layout of core nuclei (as outlined by the AHRA). (**E**) Overlaps between interpolated *PVALB*/*CALB1* maps and an AHRA-based mask of core nuclei are quantified using ROC curves for all three coordinate representations using their (**i**) MNI coordinates and (**ii**) with grey matter-remapped MNI coordinates. **(F)** To further explore the effects of coordinate choice on spatial gene expression in independent atlases, we employed universal kriging to interpolate voxelwise GABRB1 expression, a known gene expressing strongly in the medial nuclei and decreases in the lateral nuclei of the human thalamus (*25*). *Left panel*: parcellations obtained from Iglesias et al. (*36*). *Right panels*: Universal kriging-derived *GABRB1* expression maps in three coordinate sets (RMSE original: 0.22, RMSE alleninf: 0.22, and RMSE CIC: 0.20), ranging between 0.0 (yellow) and 1.0 (red) of interpolated voxelwise expression. To quantify agreement in interpolated expression between coordinate sets, voxelwise direction of maximal expression change was computed with a 3D structure tensor between three coordinate sets. Angular differences between the primary direction of maximal change were summarized by cosine similarity between coordinate sets: 1=white (aligned direction), 0=red (orthogonal), −1=(opposite direction).

### CIC coordinates provide interpolated gene expression maps that are more reflective of thalamic organization

Switching focus to the subcortex, we examined how well each set of coordinates would perform on an interpolation task. Under the well-characterized core-matrix framework proposed by Jones (*22*) and examined in a neuroimaging context by Müller et al. (*23*) and Sydnor et al. (*24*), thalamic nuclei can be grouped based on the transcriptomic identities of neuronal projections to the cortex: the ventral posterior lateral (VPL), ventral posterior medial (VPM), dorsal lateral geniculate (DLG), medial geniculate (MG), centromedian (CM), and parafascicular (Pf) nuclei are strongly identified by parvalbumin-expressing “core” neurons that project to middle layers of the cortex; these are interspersed over a matrix of calbindin-expression neurons that project to superficial layers of the cortex (*22*). Within the thalamus, core neurons are spatially constrained by traditional nuclear boundaries, while matrix neurons are unconstrained and found throughout the thalamus but are generally higher in density in areas where core cells are not found. Under this principle, nuclei dominated by core neurons (“core nuclei”) should be marked by a high ratio of parvalbumin *(PVALB)* to calbindin *(CALB1)* expression. Using the 175 AHBA tissue samples that sparsely sample the thalamus (data S14-15), we interpolated the expression values of *PVALB* and *CALB1* voxelwise, and asked: which set of coordinates produces interpolated gene expression maps that best identify the spatial layout of core nuclei as outlined by the AHRA?

First, using expert annotations of the thalamic nuclei from which samples were derived, we confirmed that *PVALB*/*CALB1* expression was much higher in core nuclei (Figure 6C). Then, using *k*-nearest neighbour interpolation (with *k* ranging from 1—i.e. nearest neighbour interpolation—to 25), we generated voxelwise maps of *PVALB, CALB1*, and their ratio, and quantified the overlap with an AHRA-based mask of core nuclei by varying thresholds and generating ROC curves. We found that CIC coordinates resulted in dramatically better spatial overlaps as quantified by the areas under the ROC curves (AUC) across all *k* (figure S8 and table S13), as evident in an example where *k* = 5 (Figure 6D-E(i) and table S14), and also when using grey matter-remapped coordinates (Figure 6E(ii), table S15, figure S9; figure S10 and table S16).

Together, the analyses above suggest that when estimating spatial gene expression (both regional averages and voxelwise), the choice of coordinate set matters, and CIC coordinates provide measures of gene expression that are more consistent with their locations under the AHRA.

### CIC coordinates perform better under alternate processing choices, and in comparison to previous coordinates used by commonly-used pipelines

To further investigate the influence of coordinate choice in downstream gene expression analysis beyond the specific decisions we made for the analyses above, we tested coordinate performance on a similar interpolation task by using a commonly-used imaging-transcriptomics software package (`abagen` (*8*)) with default pipeline parameters. We examined the expression of *GABRB1*, a medially-concentrated gene encoding the GABA-A *β*-1 subunit, whose expression is known to decrease along the medial-lateral axis in the human thalamus (*25, 26*). For regional patterns observed across thalamic subnuclei, all three coordinate sets reproduced the expected expression pattern of *GABRB1*, with high expression in the mediodorsal nuclei (MD, composed of the medial MDm and lateral MDl) and decreasing towards the ventral lateral nuclei (VL, composed of the anterior VLa and posterior VLp; Figure 6F). We report descriptive summary statistics (mean, median, interquartile range, minimum, and maximum) across all coordinate sets in table S17. Notably, both the original and alleninf maps exhibited the highest expression in the rostromedial regions, including the central medial (CeM; mean=0.60 for both) and ventral anterior nuclei (VAmc; mean=0.59 for both), whereas CIC yielded higher values in these same regions (CeM=0.93, VAmc=0.84). In contrast, the CIC expression map shows the highest expression in the MDm (0.95 vs. 0.53 for original and alleninf), followed by CeM and the anteroventral nucleus (AV; 0.91 vs. 0.57), a pattern that is more consistent with the reported expression by the Human Protein Atlas and the immunohistochemistry studies conducted in non-human primate thalamus (*26*).

To quantitatively examine the agreement of interpolated expression maps, we used 3D structure tensor analysis to compute the directions of maximal expression change, and compared the principal eigenvectors through cosine similarity across coordinate sets. Original and alleninf coordinates were strongly aligned (max voxelwise cosine similarity = 1.00) along a rostromedial-caudolateral axis. Comparison with CIC showed lower alignment, which reflects a medial-lateral direction of gene expression change. The largest divergences between coordinates were localized to the MDm, MDl, CM, and VLp nuclei (max voxelwise cosine similarity = 0.79 for both original-CIC and alleninf-CIC comparisons; Figure 6F). Previous imaging-transcriptomics study reported GABRB1 as one of the medially-expressing genes that contribute to the primary principal component spanning the medial-lateral axis in the human thalamus (*25*), similar to the results obtained when using the CIC coordinates. Taken together, these analyses show that coordinate choices affect downstream spatial gene-expression interpolations under a broader range of analytical choices that include AHBA probe selection, independent atlases, and commonly used interpolation methods.

### Choice of coordinates can impact neuroimaging associations

CIC coordinates provide better measures of spatial gene expression as compared to the other coordinate sets, but would coordinate choice make a difference in an actual imaging-transcriptomics study? To address this, we used data from a recent paper on Alzheimer’s disease-associated hypometabolism (*27*) that derived ranked gene lists by correlating AHBA spatial gene expression to a brain map obtained from a meta-analysis of FDG-PET studies. Using each set of coordinates in an identical manner, we obtained gene ranks and computed rank differences between (i) CIC and original and (ii) CIC and alleninf coordinates. We observed large differences in rank for both comparisons; more than 10% of genes have a rank difference greater than 4100, a finding that also holds for negative rank differences (Figure 7). Thus, coordinate choice can dramatically impact the identification of genes through spatial correlation analyses.

**Figure 7:**
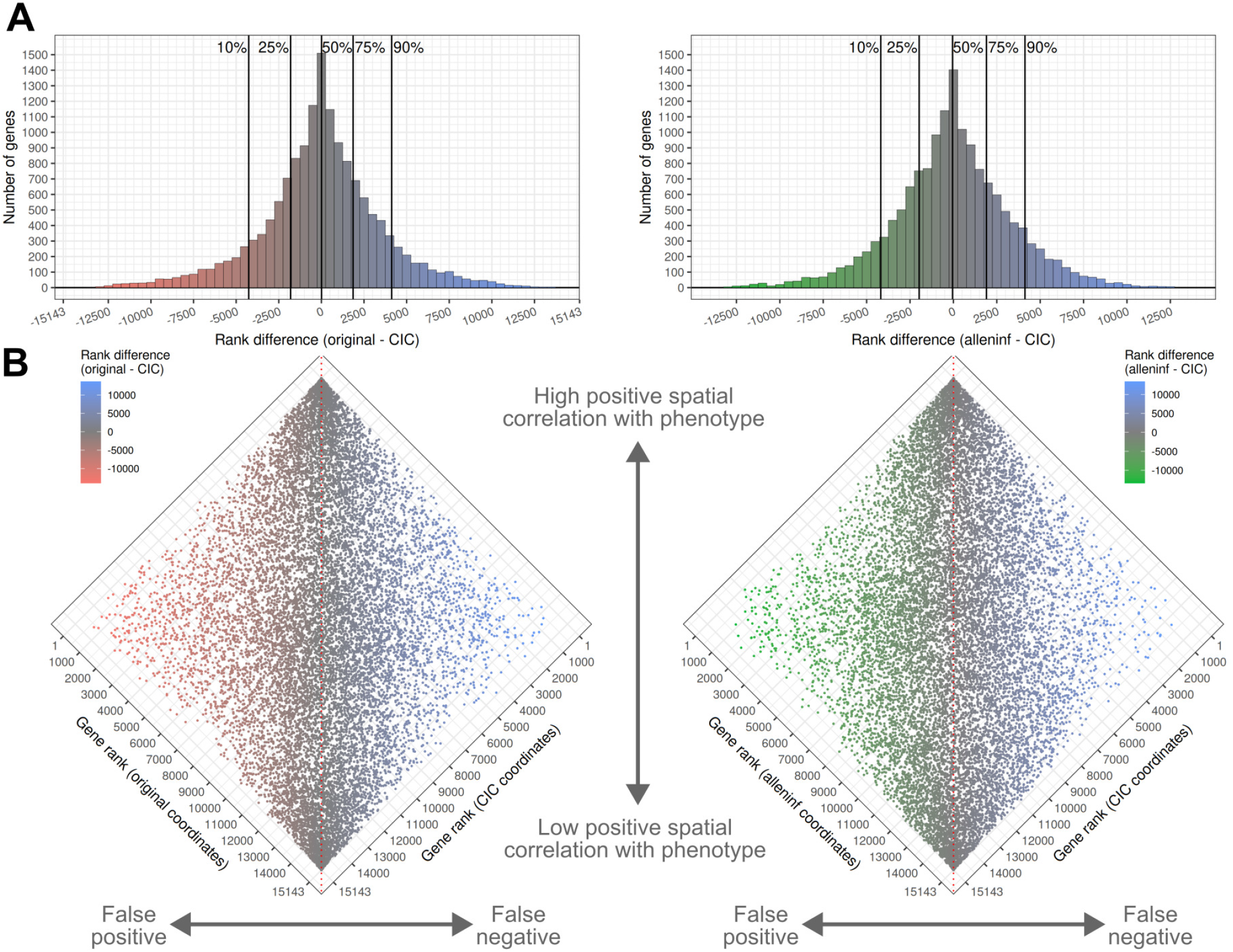
Measured bias arising from coordinate choice. (**A and B**) Differences in gene ranks between previous and CIC coordinates, based on ranking of the extent of gene expression association with the spatial phenotype described in Patel et al. (*27*). Distributions of rank differences over all genes are shown in (**A**), and are computed by comparing ranks of genes obtained using original coordinates (left subpanels) and alleninf coordinates (right subpanels) with CIC coordinates (**B**). Genes falling on the left side of (**A**) and (**B**) are more strongly associated with the phenotype under previous coordinates (i.e. false positives when viewing CIC coordinates as ground truth); genes on the right are more strongly associated with the phenotype under CIC coordinates (false negatives).

### Predicted biases due to coordinate choice in spatial gene expression analyses

Lastly, we sought to predict coordinate-associated biases in a general manner by simulating the impact of coordinate choice across a range of different spatial brain phenotypes. We generated a set of 66 brain phenotype maps using NeuroQuery (*12*) (see data S16 for a list of these terms) representing a variety of brain imaging findings across four broad classes: (i) cell types, (ii) diseases, (iii) healthy states, and (iv) signs and symptoms.

By comparing previous coordinates to CIC coordinates as before, we computed rank differences for each NeuroQuery phenotype (see figure S11 for an example) and observed large gene-wise rank differences similar in magnitude to those seen in the empirical analysis above (Figure 8A). Averaging rank differences across classes, higher mean rank differences were found for terms related to (i) cell types and neurotransmitters and (ii) signs and symptoms (Figure 8B) for both comparisons, indicating that coordinate-related biases in gene ranks might be more prominent for phenotypes associated with these classes. We further examined the variation in the magnitude of rank differences after separating genes based on positive and negative rank differences. Taking our CIC coordinates as ground truth, genes having positive rank differences are less strongly associated with the phenotype when using previous coordinates, are therefore less likely to be selected as genes of interest, and are therefore interpreted as false negatives. Similarly, genes having negative rank differences are interpreted as false positives. We quantified the extent of rank difference magnitude across all 66 maps by calculating the 80% quantile, and found that these values spanned an order of magnitude for both positive and negative rank differences (∼150-3700 for rank differences computed between original and CIC coordinates, ∼150-2200 for rank differences between alleninf and CIC coordinates, Figure 8C).

**Figure 8:**
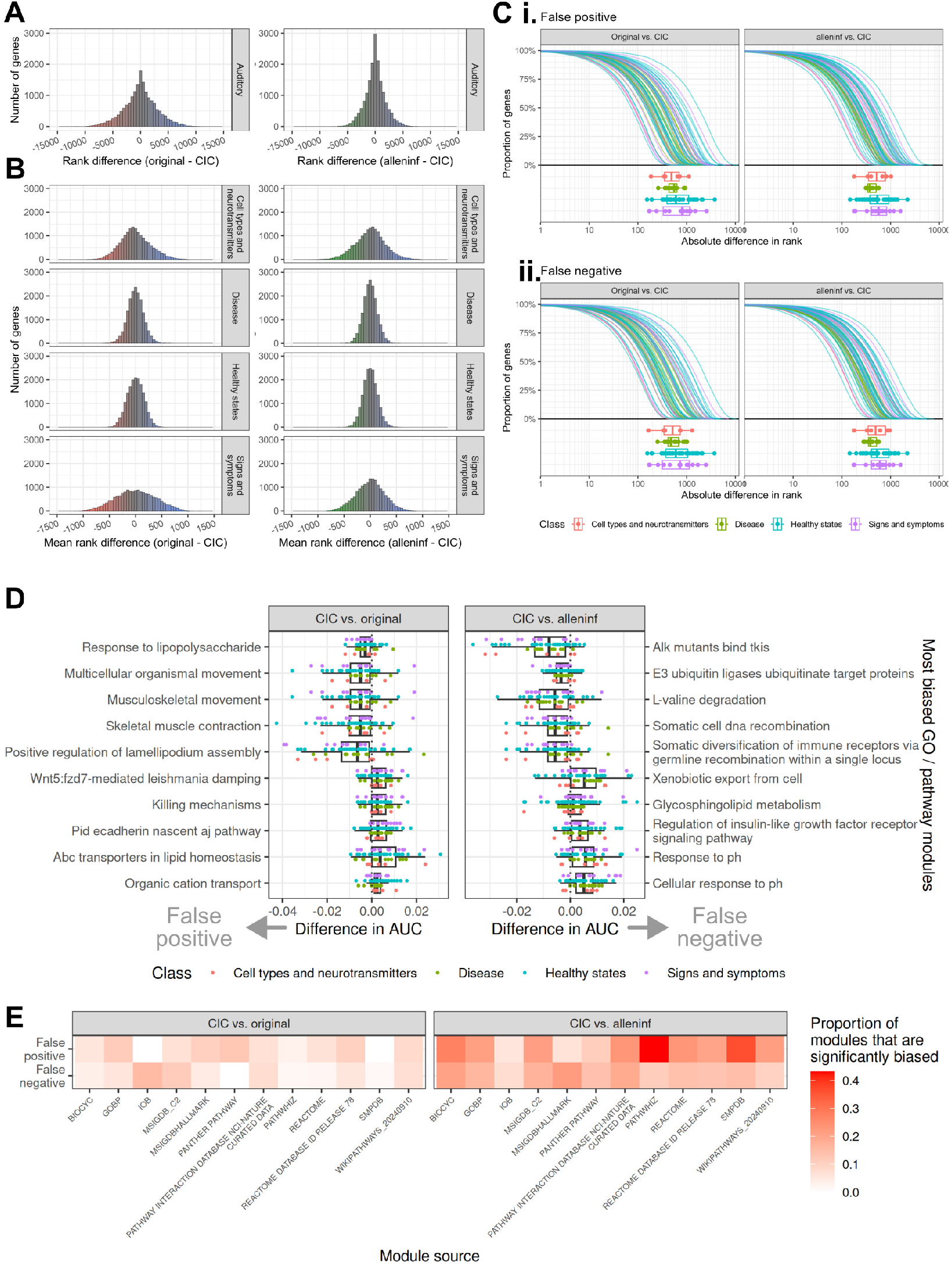
Predicted bias arising from coordinate choice. (**A**) To generalize bias across diverse neuroimaging findings, distributions of rank differences were similarly computed for 66 other spatial phenotypes obtained using NeuroQuery (shown is an example rank difference distribution for one NeuroQuery map, “auditory”). (**B**) Rank differences were averaged over classes of NeuroQuery maps. (**C**) Cumulative distributions for individual maps—separated by (**i**) false positive and (**ii**) false negative genes— are coloured by NeuroQuery class; to demonstrate the spread of rank differences over genes, the points at which 20% of genes have a greater absolute rank difference are marked for each map as points below. (**D**) Biases are also computed at the level of modules, and the most biased gene ontology and pathway modules are shown. (**E**) Numbers of significantly biased modules are presented as a function of module source.

Averaging across all 66 NeuroQuery maps, 15% of genes had an absolute mean rank difference greater than 250 when comparing original to CIC coordinates, and 10% of genes had an absolute mean rank difference greater than 250 when comparing alleninf to CIC coordinates (figure S12A(i)), suggesting the presence of genes that are both more (false positives) and less (false negatives) likely to be selected for in imaging-transcriptomics studies, regardless of the spatial phenotype. When using original coordinates, false positive genes tend to be expressed in the cortex and brainstem, while false negative genes tend to be expressed in the cerebellum. When using alleninf coordinates, false positive genes tend to be expressed throughout the brain and more strongly in the brainstem (figure S12A(ii)). These genes are enriched for a variety of biological processes and pathways, including developmental, regulatory, and metabolic processes (figure S12B).

We also examined coordinate-related biases at the level of gene modules, using gene ontology (GO) biological process modules (*28, 29*) in addition to pathway modules from a variety of sources (*30*). For each of the 66 NeuroQuery maps, we simulated typical spatial gene enrichment analysis studies separately using each of the three coordinate sets, and obtained enrichment effect sizes (quantified using the AUC metric) for all pathway and GO biological process modules (Figure 8D shows the most biased modules). When comparing differences in enrichment due to coordinate choice, we found that approximately 6% of GO biological process modules were false negatives and 11% false positives when using original coordinates, and 16% false negatives and 21% false positives when using alleninf coordinates. At the extreme end, 38 of 88 PathWhiz modules examined were identified as false positives (43%) when using alleninf coordinates (Figure 8E).

## Discussion

In this study, we provide new spatial coordinates (“CIC” coordinates, defined relative to the MNI-space ICBM152 nonlinear symmetric 2009c template (*5*)) for tissue samples from the Allen Human Brain Atlas (AHBA). We compare these coordinates to two previously described sets of coordinates (original coordinates from the Allen Institute (*1*) and updated coordinates from the ALLENINF software package (*11*)), and show that CIC coordinates provide a more accurate description of the anatomy from which gene expression data is derived. As we have further shown, differences in coordinates can lead to substantial differences in both regional and voxelwise measures of gene expression, which in turn can lead to either an under- or over-representation in genes of interest when looking for genes that correlate with neuroimaging phenotypes, along with biological process and pathway modules in downstream enrichment analyses.

### Implications for future studies

Applied to future imaging-transcriptomics studies, using CIC coordinates would presumably lead to fewer false positives in genes of interest being identified, and therefore better identification of the processes, pathways, and cell types that underlie phenotypes seen using neuroimaging. Furthermore, in cases where neuroimaging is used as a tool to precisely locate pathology and anatomical targets of treatment, CIC coordinates make linking gene expression data possible. For example, in deep brain stimulation where millimeters matter (*31*), CIC coordinates are better positioned to localize corresponding tissue samples and better understand the transcriptomic correlates of treatment response. Thus, CIC coordinates substantially improve the AHBA, a widely-used resource, and stand to make a positive impact in future studies.

### Interpreting accuracy

Although we showed that CIC coordinates performed better than original and alleninf coordinates across many tests of accuracy, we never obtained perfect classification or zero error. Given the nature of the data, we suggest that this might not be possible. For instance, in examining intensity correlations, Spearman correlations were far from unity. Even if perfect point-to-point correspondence were achieved between donor brains and template, any level of image noise that results in voxel intensity variation would also necessarily decrease the correlation (*18*). What matters however are relative measures of accuracy (quantified by Spearman correlations in the image intensity analyses)—as opposed to absolute measures of accuracy—which we show to be generally most optimal for CIC coordinates. Similarly in the interpolation task, though CIC coordinates resulted in the highest AUC, these were still far from maximum, because core neurons also project out of subregions of certain thalamic non-core nuclei. Some level of parvalbumin expression outside core nuclei is therefore expected, and so perfect identification of core nuclei based on the expression of PVALB would not be possible.

### Choice of ground truth

Our definition of ground truth are the MRI coordinates and expert annotations, both provided by the Allen Institute. We note that some imprecision might exist in MRI coordinates, which themselves are derived from a landmark-based image registration process that aligned dissection site annotations on Nissl-stained images to the donor MRI (*1*). Large sizes of tissue samples dissected may further add to ambiguity due to partial volume effects that arise, particularly for subcortical samples that were obtained via laser microdissection (*1*). Given that manual tagging is generally accepted as a form of ground truth, we took the MRI coordinates derived from this landmark-based registration as the true positions of tissue samples relative to donor brain anatomy. In the case of expert annotations, structure labels will to some extent be dependent on the neuroanatomist involved. In particular, samples derived from edges of structures might be annotated with a structure label that would be different under different neuroanatomists. In assessing anatomical location of coordinates, we used the AHRA and compared structure assignments with expert annotations. Although the AHBA and AHRA are from the same source, they notably define structures in a different manner, with the former relying on the Human Brain Atlas ontology and the latter using the Developing Human Brain Atlas ontology. The relations between the two ontologies made in this study are based on structure names and their positions within the ontology tree. Matched structures might not be spatially identical resulting in imperfect comparisons particularly at the boundaries of structures where there might be ambiguities (e.g. a sample truly labeled as being from long insular gyri (LIG) might fall within the planum polare (PLP) as defined by the AHRA, because of a varying definition of the LIG-PLP boundary). This can set an upper bound on accuracy metrics in all analyses where coordinate-based placement of samples within AHRA structures are compared to AHBA annotations. Thus, in the analysis comparing cortical lobe assignments to true dissection lobe, the number of mislocated samples may not reach zero, because of a misspecified ground truth. At present, there is no publicly-available MRI-based parcellation of the structures defined within the Human Brain Atlas, preventing further validation using the same structure definitions contained within the AHBA.

### Choice of template

We carried out all analyses in relation to the sym09c template (*5*). This required an additional transformation from original and alleninf coordinates from nlin6asym to sym09c space, a potential further source of error. To ensure this is not the case, we repeated the intensity correlation analyses in nlin6asym space, where only CIC coordinates are transformed (with the inverted version of the same transformation that original and alleninf coordinates were subject to, and therefore subject to the same source of registration error). We observed little difference in intensity correlations, implying that this error is negligible. Though the choice of template over which we compared coordinates is not important, it is an important factor in the generation of coordinates, i.e. as the target to which donor brains are aligned to. The sym09c template (produced from an alignment of the same ICBM 152 brains following a newer pipeline) provides a cleaner and sharper registration target that could explain the improvement seen with CIC coordinates. For a given set of coordinates, template choice could also account for the varying levels of accuracy seen across different donor brains. To accurately localize AHBA samples, the anatomy described by the template must be reflective of donor brains. The six donors represent both sexes, a wide range of ages, and multiple ethnicities; therefore some donor brains will presumably align better to the template because of similarities in biological characteristics with the ICBM 152 brains from which the templates are created. We examined whether this was the case, but the limited sample size of six AHBA donors prevented us from making formal inferences. Brain bank-based imaging initiatives (*32*) are well positioned to produce templates that are better suited for post-mortem imaging data, and to address this previous question.

### Sources of error

Despite showing that CIC coordinates perform best across multiple tests of accuracy, these coordinates are derived from image registration of noisy images, and there still are samples that are incorrectly localized. Inaccurate placement of coordinates can be due to two broad classes of errors: (1) errors that can be minimized with methodological refinement, and (2) inherent errors due to differences in data sources that cannot be overcome. The first class captures errors that might be attributed to parameters (relating to MRI contrast), choices in imaging (e.g. imaging post-mortem brains in cranio vs. ex cranio due to constraints at each of the two sites), and image registration. For instance, for cortical samples, surface registration techniques might offer an improvement in donor brain cortical alignment and AHBA cortical sample localization, especially when combined with our multispectral volume registration. Different sets of algorithms and/or registration parameters that tune the extent of local and global deformations might also provide additional improvement; an exploration of these choices is outside the scope of this study however. More generally, manual refinement of coordinate placement might further improve accuracy. The second class of errors relate to the existence of homologous points in donor and template brains. Any attempt at identically mapping sample locations from donor brains to specific points in other brains must make the fundamental assumption that such pairs of points exist, regardless of their methodological approach. Perfect localization of AHBA samples to anatomy—whether derived from image registration or from manual placement—requires perfect homology between source and target brains at sampling locations, which as mentioned before, might not be present between individual AHBA donors and any MNI-space template built from a group of unrepresentative subjects. Beyond differences between donor brains and template, individual differences in donor anatomy—from cortical folding (*33*) to variation in areal definitions on the cortical surface (*34*)—will likely increase the number of mislocated samples in relation to any static representation of group-averaged brain anatomy.

### Recommendations

In our analyses, we have used expert annotations along with a variety of other techniques (e.g. gene expression similarity) to quantify the accuracy of each sample’s location and assignment to a brain structure. This leads to the option of further filtering away mislocated samples before use, similar to previous recommendations (*4, 8*). There are many reasonable ways to do this; our preference is to use CIC coordinates remapped to grey matter and select samples with the following constraints: (1) following Arnatkevičiūtė et al. (*4*), samples must be shifted less than 2 mm, (2) expert annotations and sample position within the AHRA must match at a coarse level, and (3) for cortical samples, expert annotations and sample position within the AHRA must match at the level of lobes. A more restrictive filtering step of selecting samples to provide a group-wise intensity correlation above a threshold (e.g. at least 0.6 averaged over *T*_1_- and *T*_2_-weighted images) can be further carried out in each subject to improve accuracy. Additionally, given that gene expression is highly bilateral, coordinates can be mirrored to the opposite hemisphere. Finally, we note that an alternative option of directly aligning donor brains to a better suited template is another option (*35*).

Beyond suggesting filtering coordinates before use, we also recommend that future studies clearly describe datasets used. In examining previous studies to understand past use of coordinate sets, almost half of the studies we examined did not explicitly report the source of AHBA coordinates. Furthermore, as described in this manuscript and in others, there exist multiple templates that describe anatomy in MNI-space (*13*). Despite these templates having differing representations of anatomy, many studies do not make this distinction clear, leading to the possibility that there is an additional mismatch between the template to which neuroimaging findings are characterized on and the template on which coordinates describe AHBA sample locations. We therefore call on future studies to clearly report all methods used when linking AHBA data to neuroimaging.

### Summary

Neuroimaging has tremendously benefited from the Allen Human Brain Atlas. Many studies that connect neuroimaging findings with gene expression have only been made possible due to the provision of MNI coordinates that localize AHBA gene expression data to specific points in a reference brain. Yet, prior to this study, the accuracy of these coordinates were never formally validated. In this manuscript, we report a revised set of MNI-space coordinates for tissue samples from the Allen Human Brain Atlas. These new “CIC” coordinates provide an improved localization of AHBA tissue samples in relation to a standard neuroanatomical reference. In contrast to previously-reported coordinates, the calculation of these coordinates follow from an enhanced image registration process of post-mortem donor brains (that uses multispectral inputs along with manual correction steps), with the image registration target being the more modern and documented ICBM152 nonlinear symmetric 2009c MNI-space template. Using a variety of independent tests, we have verified that CIC coordinates provide a more accurate description of the true anatomical locations of tissue samples, and that the choice of coordinates can impact conclusions of neuroimaging studies that use the AHBA. CIC coordinates, along with code used to generate them, are made openly available for use.

## Supporting information

Manuscript figures (high resolution versions)

## Acknowledgments

We thank all researchers that provided information on AHBA coordinate usage in our survey of the *imaging-transcriptomics* literature.

## Author contributions

Conceptualization, Y.Y., V.L., J.C.L, G.A.D., and M.M.C.; methodology, Y.Y., V.L., A.T., B.K., and G.A.D.; investigation, Y.Y., Y.L., L.F., V.L., A.T., and G.A.D; writing-–original draft, Y.Y., V.L., and Y.L.; writing-–review & editing, Y.Y., Y.L., L.F., V.L., A.T., B.K., A.R.K., J.C.L., Y.Z., G.A.D., and M.M.C.; funding acquisition, Y.Y., A.R.K., J.C.L., and M.M.C.; resources, L.F., B.K., and Y.Z.; supervision, Y.Y., A.R.K., J.C.L., and M.M.C.

## Data and materials availability

CIC coordinates are available as data S1, on GitHub (https://github.com/CoBrALab/AllenHumanGeneMNI), and are additionally deposited at Zenodo under DOI:https://doi.org/10.5281/zenodo.3677132. All data reported in this paper are publicly available; additionally, guidance on acquiring/working with these data will be provided by the lead contact upon request.

All data processing and analyses, including the generation of CIC coordinates, were carried out using open source software. All original code has been uploaded to GitHub, and are publicly available as of the date of publication through public repositories. Specifically, code for generation of CIC coordinates are in https://github.com/CoBrALab/AllenHumanGeneMNI, and code for the analysis contained in this manuscript are in https://github.com/yohanyee/project-ahba-mni-coordinates. Any additional information required to reanalyze the data reported in this paper is available from the lead contact upon request.

## Supplementary materials

Materials and Methods

Supplementary Text

Figures S1 to S35

Tables S1 to S17

Data S1 to S17

## Supplementary Materials for

### Materials and Methods

#### Allen Human Brain Atlas

The Allen Human Brain Atlas (AHBA) consists of genome-wide gene expression values obtained from tissue that is either manually macro-dissected (mostly cortex) or laser microdissected (subcortex and brainstem) and subjected to a microarray analysis (*1*). A total of 3702 tissue samples were obtained from six donors (IDs: H0351.1009, H0351.1012, H0351.1015, H0351.1016, H0351.2001, H0351.2002). Donor brains were obtained from two sites (NICHD Brain and Tissue Bank, University of Maryland or UCI Psychiatry Brain Donor Program and Functional Genomics Laboratory, University of California, Irvine), with subjects ranging in age from 24 years to 57. Of the six donors, five were male, and of these, three were classified as White or Caucasian, and two were classified as Black or African American. The single female subject (H0351.1015) was classified as Hispanic.

*T*_1_- and *T*_2_-weighted images were acquired ex vivo for all donors, with data for brains acquired from UCI (H0351.1009, H0351.1012, H0351.1015, and H0351.1016) imaged *ex cranio*. Tissue samples were obtained from blocks that also underwent Nissl staining; dissection sites were annotated on Nissl images and transformed onto subject *T*_1_-weighted magnetic resonance (MR) images using a landmark-based registration process (*1*). Thus, for each donor, sample coordinates are originally reported as voxel coordinates relative to their MRI scan and are further annotated based on expert determination of the structures they are derived from. Expert structure annotations are chosen from the Allen Institute’s Human Brain Atlas (HBA) ontology that organizes brain structures within a hierarchical tree of regions. We take these coordinates (relative to donor brain images) and expert annotations as the ground truth.

Microarray data (including MRI and MNI coordinates reported by the Allen Institute) were downloaded from https://human.brain-map.org/static/download. Gene expression values contained within these downloaded data are already normalized, allowing for comparisons across subjects, samples, and genes. *T*_1_- and *T*_2_-weighted MR images of donor brains were downloaded from URLs found at https://human.brain-map.org/mri_viewers/data.

#### Existing coordinates describing AHBA tissue sample locations

Through affine and nonlinear alignment of donor brains to an MNI-space template, the Allen Institute further provides sample locations in MNI-space (*1*). Subsequently, Gorgolewski et al. (*11*) re-ran this subject to MNI-space template registration and thereby derived an alternate set of MNI coordinates for each sample (provided within the alleninf software package). The original and alleninf coordinates appear to be derived from donor brain image alignments to the MNI152 6th generation nonlinear asymmetric (“nlin6asym”) template (*13*), the default template with the FMRIB Software Library (FSL).

#### Allen Human Reference Atlas

Each sample can be placed into a brain structure within an anatomical atlas based on its spatial coordinates. To assign each sample a brain structure, we used the Allen Human Reference Atlas (*20*) (AHRA), a voxelwise parcellation provided relative to the MNI152 nonlinear 2009b symmetric (“sym09b”) template brain at 0.5 mm^3^ resolution. The AHRA outlines 141 structures (57 cortex, 47 subcortex, 11 brainstem, 4 cerebellum, 22 white matter or ventricles) that are described within the Allen Institute’s Developing Human Brain Atlas (DHBA) tree-based ontology. AHRA structure labels can be visualized in Figure 4A. The AHRA was used due to its three-dimensional parcellations that are of high quality in all three axes. Additionally, labels are drawn over the sym09b template, which allows for accurate registration to the sym09c template (over which most of our analyses are carried out) due to these templates being highly similar. Finally, the AHRA and AHBA originate from the same source (Allen Institute), and similarities in structure ontologies at a coarse level allow for coordinate-based structure annotations derived from the AHRA to be compared to expert annotations of tissue samples’ structural origins contained within the AHBA. The AHRA can be downloaded from: https://community.brain-map.org/t/allen-human-reference-atlas-3d-2020-new/405.

#### MNI templates

Three MNI templates are used in this study:

- MNI152 6th generation nonlinear asymmetric (“nlin6asym”) template, consisting of a *T*_1_-weighted image at a 0.5 mm^3^ resolution. Original and alleninf coordinates describe tissue sample locations relative to nlin6asym anatomy.
- MNI152 nonlinear 2009b symmetric (“sym09b”) template, consisting of *T*_1_- and *T*_2_-, and PD-weighted images at a 0.5 mm^3^ resolution. AHRA labels outline structures relative to sym09b anatomy.
- MNI152 nonlinear 2009c symmetric (“sym09c”) template, consisting of *T*_1_- and *T*_2_-, and PD-weighted images at a 1 mm^3^ resolution. Our new CIC coordinates describe tissue sample locations relative to sym09c anatomy.

All three templates describe a reference “average” brain constructed from 152 subjects. As these templates are constructed from the same source data and are highly similar, templates can be spatially aligned to each other well, with minimal image registration error. Unless otherwise specified, all analyses are carried out in relation to the sym09c template, after transforming all datasets to this template (further details below).

#### Generation of CIC coordinates

Donor brains were individually aligned to the MNI152 nonlinear 2009c symmetric (“sym09c”) template, using both *T*_1_- and *T*_2_-weighted images available for all donors and for the template.

Prior to alignment, all donor brain *T*_1_- and *T*_2_-weighted images were converted to MINC file format, and image intensities were bias-field corrected using the sqrt*(T*_1_/*T*_2_) method (*14*) or tissue-class N4 bias-field correction using the `N4BiasFieldCorrection` tool (*15*). Images were converted back to NIFTI following bias-field correction for volumetric registration to the template using the `antsRegistration` tool that is part of the Advanced Normalization Tools (ANTs) framework (*16*).

Two separate classes of alignment strategies were used depending on the AHBA donor. For five out of the six donors: H0351.1009, H0351.1012, H0351.1015, H0351.2001, and H0351.2002, alignment was achieved using a simple pipeline consisting of two separate `antsRegistration` calls (one relating to affine registration, the other relating to nonlinear registration). In the affine registration, processed and masked brain images were passed through an initial moving transform, followed by a rigid transform (gradientStep=0.5), a similarity transform (gradientStep=0.1), and two affine transforms (gradientStep=0.1 and gradientStep=0.5), with the mattes metric being optimized. In the nonlinear registration, registration was initialized with the output of the affine registration, and optimized the cross correlation metric using the symmetric normalization (*17*) (SyN) transform (gradientStep=0.1, updateFieldVarianceInVoxelSpace=3, totalFieldVarianceInVoxelSpace=0). For donor H0351.1016, the above strategy was not sufficient to produce an alignment of high quality due to the subject’s cerebellar anatomical shape and positioning. A modified strategy that separately registered the cerebellum was used: first, the cerebellum was masked out and the donor brain (without the cerebellum) were affine-aligned to the template (without the cerebellum); next, the donor cerebellum was affinely and nonlinearly aligned to the template cerebellum (after initializing the registration with the output of the previous affine transform); finally, a nonlinear registration of the full brain image was aligned to the template (after initializing the registration with the output of the previous two transforms). Parameters chosen for this alignment strategy were similar to those used for the other five donors. Parameter choices, including those listed above, followed from a series of prior registrations over which parameters were tuned to provide the most optimal alignment between images via manual evaluation. All alignments between donor and template brain images passed visual quality control.

In all instances, both *T*_1_- and *T*_2_-weighted image and template data were used to compute the registration metric being optimized, and each modality was equally weighted. Together, for each subject, these resulted in single transformations based on these multispectral optimizations that— further concatenated across all stages—map every point in the subject brain to the template. Using these transformations, sample locations within subject MR images were mapped to the template via `antsApplyTransformsToPoints`.

Scripts that exactly describe these registration steps and allow CIC coordinates to be reproduced are available on GitHub: https://github.com/CoBrALab/AllenHumanGeneMNI. CIC coordinates produced from the aforementioned steps are available both within the linked repository and as data S1. Compared to previous AHBA coordinates, main differences in the CIC coordinates are that: (1) they are provided in relation to a more modern MNI-space template (MNI152 nonlinear 2009c symmetric), (2) they are derived from multispectral image registrations applied to bias field-corrected and masked images, and (3) careful alignment was achieved through a stringent set of steps that included parameter tuning for optimal registration, thorough visual quality control, and manual refinement in cases where further improvement was necessary (i.e. subject H0351.1016).

#### Templates and coordinates

We derived CIC coordinates from donor brain image alignments to the newer MNI152 nonlinear 2009c symmetric (“sym09c”) template. Previous original and alleninf coordinates are understood to be provided in relation to the MNI152 6th generation nonlinear asymmetric template, though to the best of our knowledge, this is not explicitly described in the literature. To allow for comparisons between coordinate sets and with an anatomical atlas, we further aligned these two templates to the high resolution MNI152 nonlinear 2009b symmetric (“sym09b”) template over which the Allen Human Reference Atlas (AHRA) is defined.

These alignments were also carried out using ANTs (*16*) (tool: `antsRegistration`), and performed with an initial moving transform, followed by an affine transform (gradientStep=0.1), followed by a symmetric normalization (*17*) (SyN) transform (gradientStep=0.1, updateField-VarianceInVoxelSpace=3, totalFieldVarianceInVoxelSpace=0). In aligning sym09b to nlin6asym, we used the *T*_1_-weighted templates. In aligning sym09b to sym09c, we used *T*_1_-, *T*_2_-, and PD-weighted images. In both cases, the registration metric optimized was mutual information (32 bins). The specific `antsRegistration` calls can be found at https://github.com/yohanyee/project-ahba-mni-coordinates. We visually validated that the quality of this alignment was good.

To allow for fair comparisons between coordinates’ positions over these two different representations of a reference brain, we subsequently transformed each set of coordinates to the other template by concatenating (and inverting) transformations to and from the sym09b template. Alignment to sym09b also allowed us to transform AHRA atlas labels to each template and examine coordinate positions within structures using their native templates. Thus, for both nlin6asym and sym09c templates, we obtained template-specific coordinates for each of the three coordinate sets, along with template-specific AHRA structure labels. Primary analyses comparing coordinate sets were performed in relation to both templates to ensure that results are not dependent on the choice of the template.

#### Remapping of coordinates to grey matter

We observed that in all three coordinate sets, a large proportion of grey matter samples were positioned either in white matter or outside the brain, albeit less so when using the CIC coordinates. This has been acknowledged by others (*4*), and repositioning sample locations to the nearest cortical tissue has been suggested as part of preprocessing pipelines. Following this suggestion, for all three coordinate sets, we proceeded to map all samples truly originating within grey matter to their nearest grey matter voxel within the MNI sym09c template. Nearest grey matter voxels were determined using the fast marching method (*37*) (as implemented in the `scikit-fmm` package for `Python`) to compute Euclidean distance to all grey matter voxels.

#### Relating AHBA and AHRA structures

Both the AHRA and AHBA region definitions are contained within tree-based ontologies (*1, 20*), but notably, different ontologies are used (AHBA ontology: “Human Brain Atlas”, StructureGraph ID: 10; AHRA ontology: “Developing Human Brain Atlas”, StructureGraph ID: 16), and a one-to-one mapping of structures within the AHBA and AHRA is not possible in certain areas of the brain such as the hippocampus. The tree-based definitions allow for structures to be grouped together as coarser regions however, allowing for one-to-one mappings at different levels of granularity and in certain substructures, thus making the AHRA an ideal atlas to use. In this manuscript, AHBA and AHRA structures are compared in several different cases: (1) at a coarse level, (2) cortical lobes, and (3) thalamic subdivisions. In each case, mappings were manually ascertained using structure names and their positions within their respective ontological hierarchy. Mappings are provided in data S4 (coarse level mappings), data S6 (cortical lobes), and data S14 (thalamic subdivision mappings).

#### Gene expression

We used gene expression values from a filtered set of 17205 probes (corresponding to 16678 unique gene symbols, 16606 unique Entrez IDs, 16685 gene names; see data S17). Probe selection is described in Zeighami et al. (*38*) which follows methods in Hawrylycz et al. (*39*).

#### Meta-analytic brain maps

In order to assess the practical impacts that coordinate set choice imparts, we used NeuroQuery (*12*), a meta-analytic tool that, given text strings, produces predicted spatial brain maps based on a model trained on prior literature. Prior to any analysis, we produced a list of interesting terms meant to capture a range of spatial brain imaging phenotypes. 66 terms in total were identified across four categories (healthy states, signs and symptoms, disease, and cell types and neurotransmitters), and are listed in data S16. For each of these terms, we generated a spatial map using the `neuroquery` package for `Python`. Maps, generated in relation to the nlin6asym template, were subsequently transformed to resampled to the sym09c template for analysis using the transformations described above.

#### Survey of AHBA coordinate use within neuroimaging studies

To estimate the prevalence of the different AHBA coordinate sets in prior neuroimaging research, we obtained a representative sample of studies through a PubMed database search (June 1st, 2023) using the following keywords: “(Allen Human Brain Atlas OR AHBA) AND (gene expression OR transcriptome OR microarray OR enrichment) AND (MRI OR neuroimage)”. To be clear, these search terms were chosen to provide a sample of similar studies from which we could calculate the proportion that each coordinate set was used; they do not—and are not meant to—capture every neuroimaging study that uses the AHBA.

The search resulted in 160 potentially relevant articles, and 97 were excluded for mismatches in type, topic, and content (Figure S14). The 63 studies included were neuroimaging studies that utilized the AHBA microarray data to examine gene expression or gene enrichment of specific regions of interest in an unbiased manner, without focusing on specific cell types or disease contexts. The studies providing workable gene enrichment data were used for the NeuroQuery analyses.

Among the 63 included studies, we determined their usage of coordinates by thoroughly reviewing the articles and their supplementary materials. We recorded the choice of coordinates when explicitly stated in the article. For studies that did not provide explicit information on the set of coordinates used, we made the following assumptions:

- If a study did not specify how the AHBA microarray data was mapped to its own data and did not cite any other set of coordinates, we assumed the original coordinates provided by AHBA were used.
- If a study cited the `abagen` toolbox (*8*), we assumed the alleninf coordinates were used.
- If a study cited a pipeline for spatial transcriptomics but without recommending a specific set of coordinates, we labeled it as unsure.

Additionally, some coordinate usage information was provided by the authors through email correspondence.

We note that this survey was carried out in 2023, and since then, CIC coordinates have been used in additional research studies (*40*).

### Quantification and statistical analysis

#### Determination of affine-only alignment for original AHBA coordinates

Affine-only alignment of donor brains to the MNI152 6th generation nonlinear asymmetric (“nlin6asym”) template was determined by plotting MRI voxel coordinates of all samples against original MNI-space world coordinates. Perfect collinearity in this relationship points to an alignment that has no nonlinear component.

To examine the extent of nonlinear warping associated with the generation of original coordinates for samples from the two donor brains found to have a nonlinear component, and to compare this to nonlinearities associated with alleninf and CIC mappings, we defined a nonlinearity index as the geometric mean of 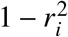, where 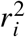 is the adjusted *R*^2^ value of the linear regression predicting the -axis world coordinate from the *i*-axis voxel coordinate.

#### Comparison of original, alleninf, and CIC coordinates

We first visualized sample position by rendering random samples’ locations over their respective image. Sample positions based on original and alleninf coordinates were rendered over the nlin6asym template, positions based on CIC coordinates were rendered over the sym09c template, and sample locations within donor brains were rendered over the donor’s MR *T*_1_-weighted image.

To understand how different the three coordinate sets are, and whether any two coordinate sets provide a more similar representation of sample location, we computed the pairwise distance for each sample separately, after transforming original and alleninf coordinates to the sym09c template. Distributions of distances and their correlations were subsequently computed.

#### Validation of neuroanatomical accuracy based on MRI image intensities

In the intensity-based correlation analysis, we correlated intensity values of donor MR images with intensity values of the template under sample locations. Intensity data were extracted using `RMINC`, an `R` package for working with the MINC file format. Sample positions, represented as world coordinates, were converted to image-specific voxel coordinates and extracted using the `mincGetVoxel` function. Intensity values were extracted for both sym09c templates and nlin6asym templates after transforming the appropriate coordinate sets to each template, and for all available modalities (*T*_1_- and *T*_2_-weighted for sym09c, *T*_1_-weighted for nlin6asym).

Relationships between subject image and template were quantified using Spearman’s correlation, separately for each subject. To compute confidence intervals, we additionally bootstrapped this correlation 1000 times by resampling with replacement. This procedure was repeated for subsets of samples grouped by the coarse brain structure from where they were dissected (cortex, subcortex, brainstem, cerebellum). Additionally, this procedure was also repeated using grey matter-remapped coordinates.

In order to determine the most optimal set of coordinates for each subject, the coordinates providing the highest Spearman correlation was chosen (implying that subject-template alignment was best); if other coordinate sets whose Spearman correlation fell within the 95% confidence interval of this optimal set, they were considered equivalent.

Finally, we also examined the relationship between the quality of donor brain alignment with the template (as approximated by the Spearman correlation) and donor biological and sampling characteristics. We used linear mixed effects models to predict (Fisher-transformed) spearman correlation values *ρ* given donor characteristic (age; sex; ethnicity; handedness; source) as a fixed effect; these slopes were constrained but intercepts were allowed to vary (random effect) across the different coordinate sets and MR contrasts (`R` formula notation):

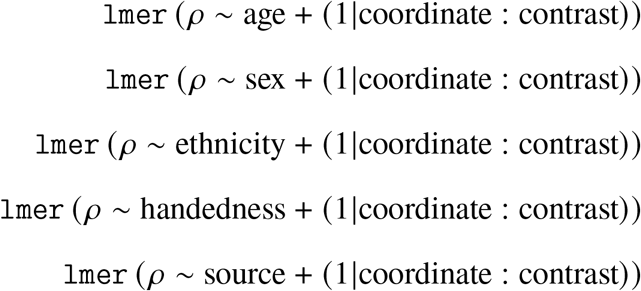

#### Validation of neuroanatomical accuracy based on atlas structure annotations

In this analysis, expert annotation of structures that samples were dissected from were compared to the anatomical structure that the sample is placed within the AHRA by its coordinates. Given that expert annotations were based on the HBA (Allen Institute’s ontology StructureGraph ID: 10) whereas the AHRA structures are based on the DHBA (Allen Institute’s ontology StructureGraph ID: 16), a mapping between the HBA and DHBA had to be made, as described in the Relating AHBA and AHRA structures subsection. At a coarse level, five shared structures could be identified: cortex, subcortex, brainstem, cerebellum, white matter and ventricles. White matter and ventricles were grouped together due to their distinct positions and low sample size. A sixth label, “Outside brain mask”, was considered when a sample’s spatial coordinates placed it outside any AHRA label. Thus, for each sample and coordinate set, a sample contained two annotations drawn from this list, one based on expert annotation (ground truth) and one based on its spatial coordinate relative to the AHRA.

Agreements between expert annotations and anatomical locations within the AHRA atlas were assessed using confusion matrices, which described the number of samples shared between true (expert) annotations and coordinate-based annotations. Separately for groups of samples truly drawn from each coarse structure, accuracy of coordinates were assessed using the false negative rate (proportion of misclassified samples within each ground truth category). These measures were computed both individually for each donor and grouped across all six donors.

Finally, we repeated this analysis using grey matter-remapped coordinates. Given the large number of samples that were remapped to the cortex, we also carried out a secondary analysis examining concordance at the level of cortical lobes, where mapping between the HBA and DHBA were possible.

#### Validation of neuroanatomical accuracy based on similarities in gene expression profiles

The goal of this analysis was to validate coordinate-based sample placement within AHRA structures by examining how similar samples’ gene expression profiles were to structures’ expected gene expression profiles. For this analysis, we restricted ourselves to samples that were placed into different AHRA structures when using different sets of coordinates. Samples that were placed in separate AHRA structures when comparing (a) original and CIC coordinates and (b) alleninf and CIC coordinates were identified. For each such sample and comparison of coordinate sets, we identified every other sample located in the two different structures. Then for each gene, we computed the mean gene expression across all identified samples (not including the sample being examined) in each structure. Gene expression of the sample being examined was correlated with the two mean gene expression profiles of the two structures associated with the two coordinate sets being compared, with the idea that there is a greater likelihood that the sample being examined was drawn from the structure with which there is a higher correlation. To quantify differences between coordinate sets, we computed differences between these expression profile correlations, after applying a Fisher transformation. The analysis was repeated using coordinates relative to the nlin6asym template, and also with grey matter-remapped coordinates.

#### Validation of neuroanatomical accuracy based on discriminability of cortical structures

Similar to the above analysis, we tested how well transcriptome-based positioning of samples within neighbouring but transcriptionally-distinct anatomical structures reflected coordinate-based positioning (Supplementary Text). Our approach was to select pairs of nearby structures, assign samples to each based on their gene expression profiles, and subsequently test how well the samples’ coordinate-based positions within AHRA predict their expression-based assignment.

To carry out this approach, we began by a priori choosing pairs of nearby structures that could be identified within both the AHBA and AHRA. All pairs of structures identified were cortical:

- Precentral gyrus and postcentral gyrus
- Occipital pole and cuneus
- Cuneus and lingual gyrus
- Superior and inferior temporal gyrus
- Superior and middle temporal gyrus
- Middle and inferior temporal gyrus
- Superior and middle frontal gyrus
- Medial and lateral orbital gyrus

Next, we selected samples that were within either structure based on ground-truth (AHBA) annotations, and assigned samples to either structure by training a LASSO (*41*) model to predict numerically-encoded structure assignment (0 or 1) given the expression values of all genes and ground-truth (AHBA) annotation. 10-fold cross-validation was used, and the selected sparsity level (i.e. number of genes) minimized cross-validation error. We used the `biglasso` package (*42*) for `R` (`cv.biglasso`), with binomial family function, elastic net penalty (elastic net mixing parameter *α* = 0.5). We obtained class predictions for samples using this model and compared these to expert annotations, and discarded any pairs where predictions were discordant with ground truth *(>* 10% misclassified samples; these represent transcriptomically-instinguishable pairs). For remaining pairs of structures, we explicitly assigned these predicted structures to samples as their expression-based regional identity. Using these gene expression-based assignments of structures, we examined the proportion of samples misclassified (i.e. the proportion of samples where their expression-based structure class assignment did not match their coordinate-based structure assignment) as a function of coordinate set, both pooled over all subjects and also within individual subjects. We also conducted a receiver-operator-curve (ROC) analysis by varying thresholds over models’ predicted response values (ranging between 0 and 1) and computing binary overlap with coordinate-based structure assignment. This analysis was repeated using grey matter-remapped coordinates.

#### Validation of neuroanatomical accuracy based on identifiability of thalamic organization

We designed an interpolation task in which we tested which set of coordinates provided the most expected interpolation pattern of known gene expression. We relied on the core-matrix framework (*22*) that describes a spatially varying distribution of parvalbumin *(PVALB)*-expressing “core” neurons and calbindin *(CALB1)*-expressing “matrix” neurons. Under this framework, thalamic nuclei can be partitioned into those that contain high amounts of core neurons (and consequently a high *PVALB*/*CALB1* ratio), and those that do not. The goal of this analysis was to determine which coordinate set best identifies the spatial patterns of core nuclei (as described by the AHRA), based on the interpolated values of *PVALB* (AHBA probe name: A_23_P17844) and *CALB1* (AHBA probe name: CUST_16773_PI416261804).

For each sample, we acquired *PVALB* and *CALB1* expression and normalized expression values to lie between 0 and 1 through a centering and standard deviation scaling of values, followed by an arctan and subsequent affine transformation. The ratio *(PVALB*/*CALB1)* of these normalized values are interpreted as a fold change, with values greater than 1 indicating that samples are likely derived from core nuclei. First, we classified AHBA thalamic nuclei into “core”, “matrix”, or “mixed” groups based on literature, and confirmed that samples truly dissected from core nuclei has fold change greater than 1, and that samples from matrix nuclei had a fold change of less than 1. After confirming this, we proceeded to use samples’ coordinate-based positions to interpolate fold-change values over all voxels within the thalamus, and spatially compared this to AHRA segmentations of thalamic nuclei.

For interpolation, we used 175 samples that were truly dissected from the thalamus (Data S15), and further mirrored these across the midline, resulting in 350 spatial points (positioned differently by each coordinate set) to guide the interpolation. We used *k*-nearest neighbours interpolation *(k* ranging from 1—corresponding to nearest neighbour interpolation—to 25), where interpolated values are computed based on the inverse-distance-weighted values of *k* nearest samples. Spatial concordance between the interpolated *PVALB*/*CALB1* ratio and AHRA segmentations of core nuclei were examined using ROC curves constructed from thresholding fold change values, and were compared between coordinate sets. This analysis was also repeated using grey matter-remapped coordinates.

#### Validation of neuroanatomical accuracy under separate processing choices

We tested whether coordinate choice alters downstream imaging-transcriptomic results beyond the above processing choices. We focused on *GABRB1*, which shows higher expression in medial thalamic nuclei and lower expression laterally (*25, 26*) using previously published pipelines commonly employed in imaging-transcriptomic studies (*8, 25, 43*). Briefly, the thalamic atlas from the `Freesurfer5` (v8.1.0, released in sym09c space) was first registered to the nlin6asym space, using rigid and affine transformations (`Greedy6`, https://github.com/pyushkevich/greedy) obtained from a *T*_1_-weighted nlin6asym to *T*_1_-weighted sym09c template registration. This thalamic atlas was chosen due to its integration of *ex vivo* MRI-derived structural information and histology-informed parcellation of thalamic subnuclei (*36*). Samples within the thalamus were extracted using original, alleninf, and CIC coordinates, and processed via the `abagen` toolbox (*8*) (https://github.com/rmarkello/abagen) using default parameters. Microarray probes were first reannotated (*44*), with probes unmatched to a valid Entrez ID discarded. Probes were then filtered relative to background noise (probes with intensity less than the background in >=50% of samples across donors were discharged with an intensity-based threshold of 0.5), and selected based on differential stability across donors. Samples were assigned to the thalamus if their coordinates were within 2 mm of the parcel as per the standard parameter (*8*). As samples from the right hemisphere were only present in two subjects, only samples from the left thalamus (original and alleninf coordinates: n=125; CIC coordinate: n=131) were included in analysis.

Voxelwise *GABRB1* expression was calculated through universal kriging (`gstat`, R) for 3D spatially-aware interpolation as described in previous works (*25, 43*), producing three expression maps in three coordinate sets. Leave-one-out cross-validation was performed for each coordinate set. For each thalamic sample *i* = 1..*n*, we removed sample *i* and fit the same universal kriging model on the remaining *n* −1 samples. The predicted expression was then compared to the observed expression at the location *ŷ*_*i*_. We summarized error with the root mean squared error (RMSE), calculated as:

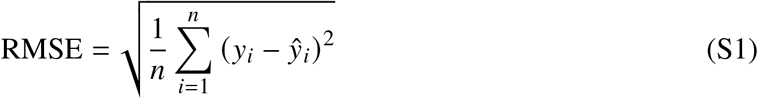

All voxelwise gene maps were registered back into the sym09c template space using the inverse transformation to ensure correspondence throughout analysis. Next, we computed local orientation with an adapted 3D structure tensor to characterize the direction of maximal expression change in each coordinate set (https://github.com/Bradley-Karat/Structure_Tensor_Python). Briefly, we computed the partial derivative along each axis of the expression map *I(x, y, z)* to obtain the variation in each direction and smoothed with a Gaussian kernel *(σ*_*d*_ = 1.0):

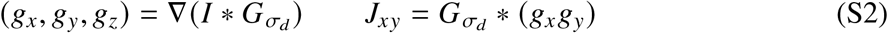

The partial derivatives are then combined and smoothed with a second Gaussian kernel *(σ*_*i*_ = 3.0) to construct a 3 × 3 structure tensor *(J)* matrix:

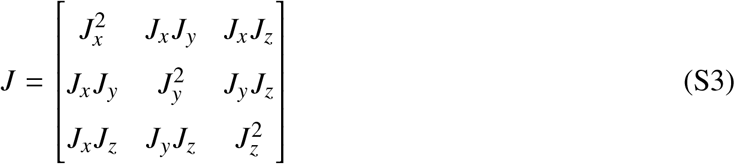

The direction of maximal expression change is given by the largest eigenvectors 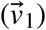 through eigendecomposition of *J* at each voxel. To remove edge artifacts, we interpolated voxelwise expression to pad the voxels surrounding the thalamic regions for structure tensor analysis, then masked with the thalamic parcellations to restore the original array shape. Cosine similarity was then performed to quantify the angular differences between the 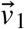 of different coordinates (Python v3.13.5):

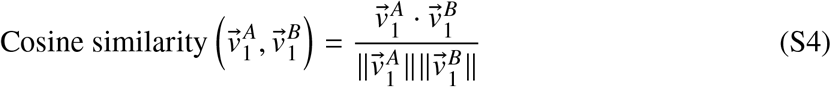

The result of cosine similarity ranges from −1 (opposite directions) to +1 (aligned directions), with 0 indicating orthogonality.

#### Observed effects of coordinate choice on a real study

We used data from Patel et al. (*27*), and for 15143 genes examined in that study, ranked these genes by correlation with phenotype using each of the three coordinate sets. For a given coordinate set, the gene with the highest positive correlation is ranked first, and the gene with the highest negative correlation is ranked last. For each gene, rank differences were computed between original and CIC coordinates (original rank - CIC rank), and alleninf and CIC coordinates (alleninf rank - CIC rank); genes with a positive rank difference have a higher correlation with the phenotype when using CIC coordinates as compared to previous coordinates, and vice versa. We further summarized the distributions of these rank differences by computing sample quantiles.

#### Predicted effects of coordinate choice on the field of imaging-transcriptomics

To predict the effects of coordinate choice in a more general context, we applied the above methods to the set of 66 meta-analytic brain maps obtained through NeuroQuery (*12*) (selected a priori to reflect a range of brain states and conditions). For each of the three coordinate sets, we extracted the activation values under reported sample locations, and for each of the 17205 probes, correlated these extracted activation values with spatial expression. For each probe, NeuroQuery map, and coordinate set, we computed the Pearson correlation between gene expression and coordinate-based activation values over all samples, and subsequently for each NeuroQuery map and coordinate set, ranked genes based on these correlations. Using these gene ranks, we predicted the effects that coordinate choice have on imaging-transcriptomic studies in (a) the genes that can be identified, and (b) downstream enrichment modules (GO terms and pathways).

At the level of genes, we searched for genes that produced consistently biased shifts (seen over multiple NeuroQuery maps) when using the original and alleninf coordinate sets as compared to CIC coordinates. For each gene, we computed the mean rank difference over all NeuroQuery maps and over subsets of NeuroQuery maps grouped by their classification, resulting in a distribution of genes in which genes at the tail ends of this distribution can be interpreted as being biased (i.e. they are more likely to be ranked higher/lower when using previous coordinate sets as compared to CIC coordinates). To interpret these genes, we tested for enrichment within gene sets obtained from the Bader lab at the University of Toronto (*30*) (https://baderlab.org/GeneSets; gene set: Human_GOBP_AllPathways_withPFOCR_no_GO_iea_October_01_2024_symbol.gmt, consisting of GO biological process terms and pathway modules, but excluding GO terms inferred from electronic annotation). Gene sets were restricted to those that contain between 10 and 200 genes (*45*), and the Mann-Whitney U-test (as implemented in the `tmod` package (*46*) for `R`) was used to compute enrichment from ranks.

At the level of enrichment modules, we used gene ranks obtained for each NeuroQuery map and coordinate set to directly produce a set of associated terms, mimicking typical imaging-transcriptomic studies. We looked for terms that were consistently more or less likely to be identified when using original and alleninf coordinate sets as compared to CIC coordinates. Module enrichment were determined in the same way as above (using the same gene set, filtering to between 10 and 200 genes, and using the U-test contained within the `tmod` package). Differences in AUC metric (returned by the U-test) as a result of using previous coordinates instead of CIC coordinates were computed; modules that consistently produced positive/negative differences can be interpreted as being less/more likely to be obtained in a general imaging-transcriptomic study when using original or alleninf coordinates, and thus indicate potential biases that may arise in downstream inferences when using previous coordinates. Modules that contained AUC difference values significantly different from zero (computed over all NeuroQuery maps and after p-value adjustment for multiple comparisons) were annotated as being false negatives or false positives, depending on the direction of AUC difference. Finally, we verified our predictions of bias by comparing mean AUC difference values against AUC differences observed in modules obtained from the ranked gene lists in the Patel et al. (*27*) study.

### Supplementary Text

#### Determination of affine-only alignment for original AHBA coordinates

Although the AHBA Microarray Survey technical whitepaper states that four brains undergo deformable registration via ANTs, original (MNI) coordinates can be mapped to MRI coordinates using an entirely affine transformation. We plotted MRI voxel coordinates against world coordinates and discovered perfect collinear relationships for four out of the six *ex cranio* donor brains (Figure S15).

#### Association of alignment quality with donor demographics

Recent studies have emphasized the importance of population-specific templates in accurate alignment of individual brain images to templates (*47, 48*). To this end, we examined whether the magnitude of Spearman correlations—a proxy for registration accuracy—could be related to donor-specific characteristics in our intensity-based analysis. Through linear mixed effects models (Figure 3C), we found a small positive effect of Hispanic ethnicity (estimate=0.206, t=3.183, adjusted p-value=0.025) after correcting for false discovery rate (*49*) across the different coordinate sets (original, alleninf, CIC) and MR contrasts (*T*_1_-weighted, *T*_2_-weighted). Intensity values obtained from coordinates derived from the Hispanic donor (H0351.1015) tended to be higher than other ethnicities. Sample sizes are too low to confidently generalize such relationships however, and the presence of collinearities (the only Hispanic subject is also the only female subject and brains from the NICHD source are all from Black or African American donors) makes interpreting these relationships challenging.

#### Effect of template choice on analysis

Original and alleninf coordinates are provided in relation to the MNI152 6th generation nonlinear asymmetric (“nlin6asym”) template, while CIC coordinates were derived from donor brain alignments to the MNI ICBM152 nonlinear symmetric 2009c (“sym09c”) template. All analyses comparing coordinate representations within this manuscript are carried out in relation to the same template, either by transforming original and alleninf MNI coordinates to the sym09c template via a nlin6asym to sym09c template alignment, or vice versa. Primary results described in the Results section are in relation to the sym09c template. To ensure that our results are robust and are not affected by errors within the additional nlin6asym to sym09c alignment, we repeated major analyses relative to the nlin6asym template. Here, despite the CIC coordinates undergoing an additional transformation, almost identical results were obtained. In the analyses comparing subject-template image intensity correlations under sampling locations, CIC coordinates remained better than (or equal to the best) other coordinate representations (Figure S16 and Figure S17).

#### Prediction of dissection structure from coordinate-based position within the AHRA

We note that the FNR is a measure of incorrectly categorized observations, given the true categories, and the many samples misplaced into the white matter/ventricles or outside the brain explains the high FNR observed. In practice, it is also helpful to know the converse: for example, if a sample is found to be located in the brainstem, what is the probability that it was actually dissected from the brainstem? This probability of correct classification given the observed category is the positive predictive value (PPV), equivalently expressed as the false discovery rate (FDR; 1-PPV), is also most optimal when using CIC coordinates (Figures S18-23). Excluding the high FDR for samples classified as white matter based on location within the atlas, the FDR was reasonable (<10% pooling across all subjects) compared to the inflated FNR seen for all coordinate sets, though using CIC coordinates resulted in a further lowering of the FDR in all except one comparison (vs. alleninf in the brainstem, where the FDR was almost equivalent).

#### Concordance between coordinate representations

Each coordinate representation (original, alleninf, CIC) generally placed tissue samples in distinct locations of MNI-space, with little similarity to the other two coordinate sets (Figure S24).

In general, assignment of samples to structures were quite different when using different coordinate sets, both across coarse structures and also within Figure 4A. At the coarse structure level, 56.4% of all samples (2087 of 3702 samples) were placed in different regional labels in at least one of the three coordinate representations, with this proportion being highest for samples truly dissected from the cortex (61.9%) and lowest for samples from the brainstem (40.6%). In instances where two of the three coordinate representations agreed, original coordinates were most discordant (770 of 3702 samples placed in a structure different from that which is obtained when using either of the other two coordinate representations), though these numbers were high for alleninf (523 samples) and CIC (506 samples) coordinates as well. Additionally, this concordance in sample placement between coordinate representations was subject dependent, with CIC coordinates being most discordant in sample placement for subjects H0351.1009 and H0351.1012, and alleninf coordinates for subjects H0351.2001 and H0351.2002 (Figures S25-27).

#### Gene expression-based separation of samples assigned to transcriptome-distinct regions through spatial coordinates

In an additional analysis of coordinate accuracy using gene expression data, we sought to determine how well each coordinate set is able to correctly place samples into nearby structures. In the previous analysis, correlations were generally high because all genes were used, regardless of whether a gene’s expression level differed between the two structures. Here, we explicitly assigned a regional identity to each sample based on a LASSO (*41*) model that learned the weights of a limited set of genes distinct in their expression between the two structures. We then tested whether each sample’s coordinate-based position under AHRA labels matched expression-based regional identity.

Specifically, we trained the LASSO model using expert annotations of tissue samples’ structures to determine a sparse set of marker genes that were most different in their expression levels between the precentral and postcentral gyrus. We found a set of 10 genes with distinct expression levels in this pair of structures *(CBLN2, CNTN6, SPHKAP, CTXN3, LGALS2, CREB3L3, BTNL9, TG, MET, TSHZ3)* that minimized cross-validation error (Figure S28), and confirmed that the gene expression-based LASSO model was able to separate samples obtained from the pre- and postcentral gyrus well (2% of samples were misclassified by the model as compared to ground truth annotations), indicating that these two regions are transcriptomically distinct. Then, we examined which set of coordinates would place samples within AHRA-defined regions that matched the LASSO-based annotations, and found that CIC coordinates performed the best at separating brain regions based on their samples’ gene expression profiles (Figure S29). We also confirmed this to be true with other neighbouring pairs of transcriptomically distinct cortical regions, namely: superior vs. inferior temporal gyrus (Figure S30), superior vs. middle temporal gyrus (Figure S31), middle vs. inferior temporal gyrus (Figure S32), superior vs. middle frontal gyrus (Figure S33), and medial vs. lateral orbital gyrus (Figure S34). CIC coordinates continue to separate transcriptomically distinct structures when using grey matter-remapped coordinates (Figure S35).

**Figure S1:**
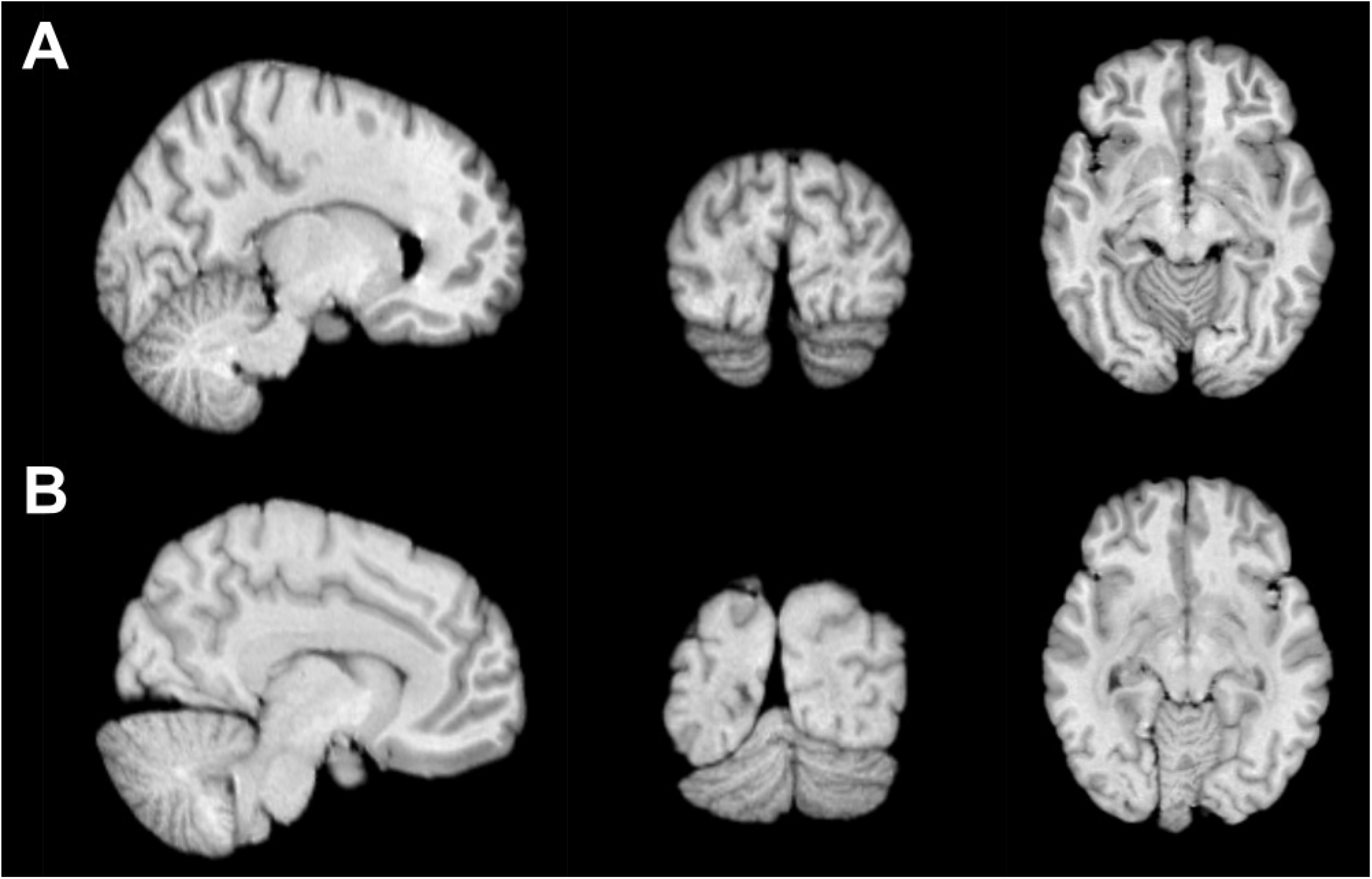
Abnormal deviation of cerebellar position requiring manual correction. Normal and abnormal positioning of the cerebellum is shown in each of the three axes for (A) subject H0351.1015 and (B) subject H0351.1016 respectively.

**Figure S2:**
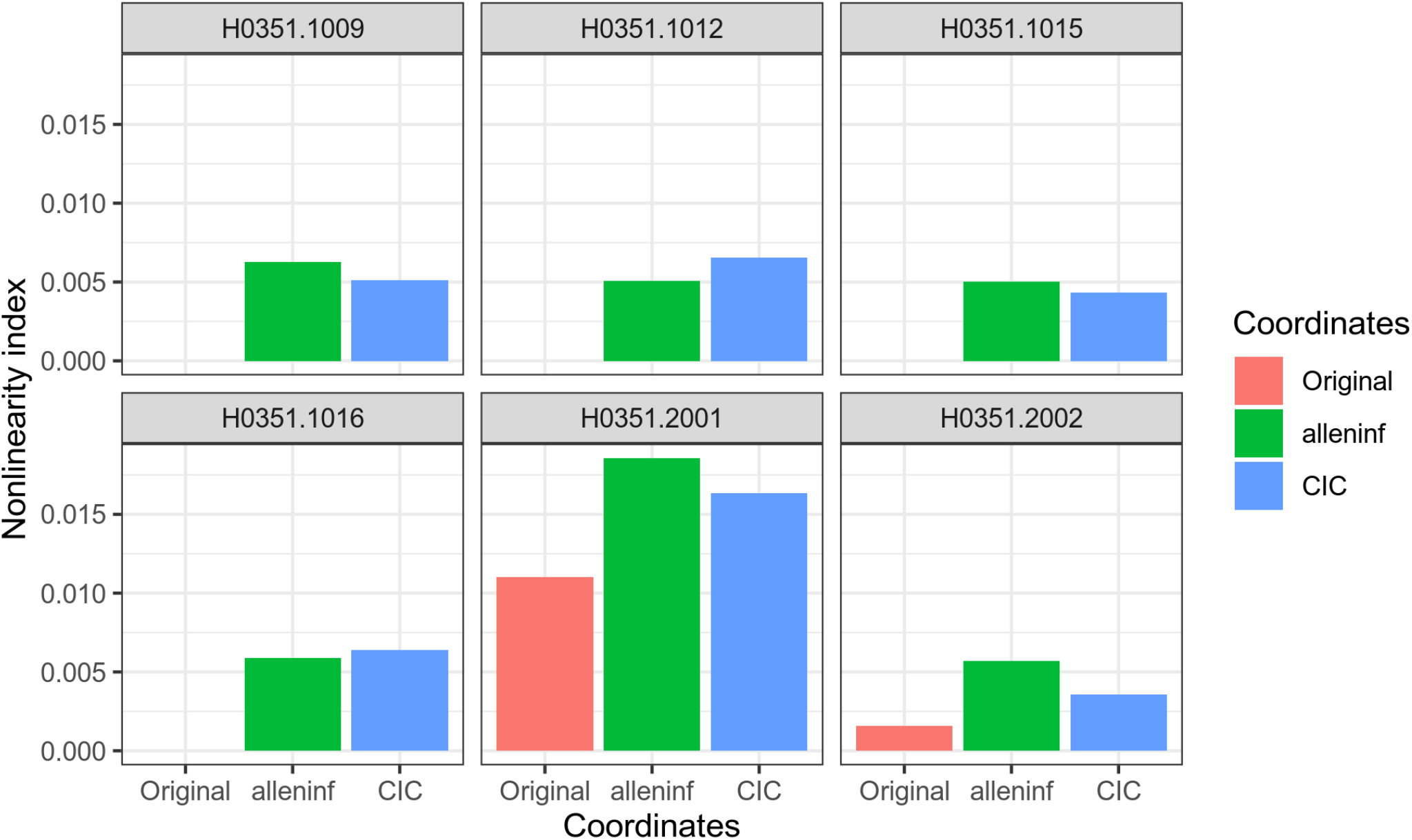
Extent of nonlinear deformations of donor brains used in MRI locations to MNI-space templates. For each subject, nonlinearity indices measuring the deviation from a linear relationship between MRI and MNI coordinates are computed separately for original, alleninf, and CIC coordinates. Briefly, the nonlinearity index is computed as the geometric mean of 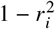 values obtained from three models, each predicting the MNI-space coordinates from MRI coordinates along an axis. See (*50*) for definition of the nonlinearity index.

**Figure S3:**
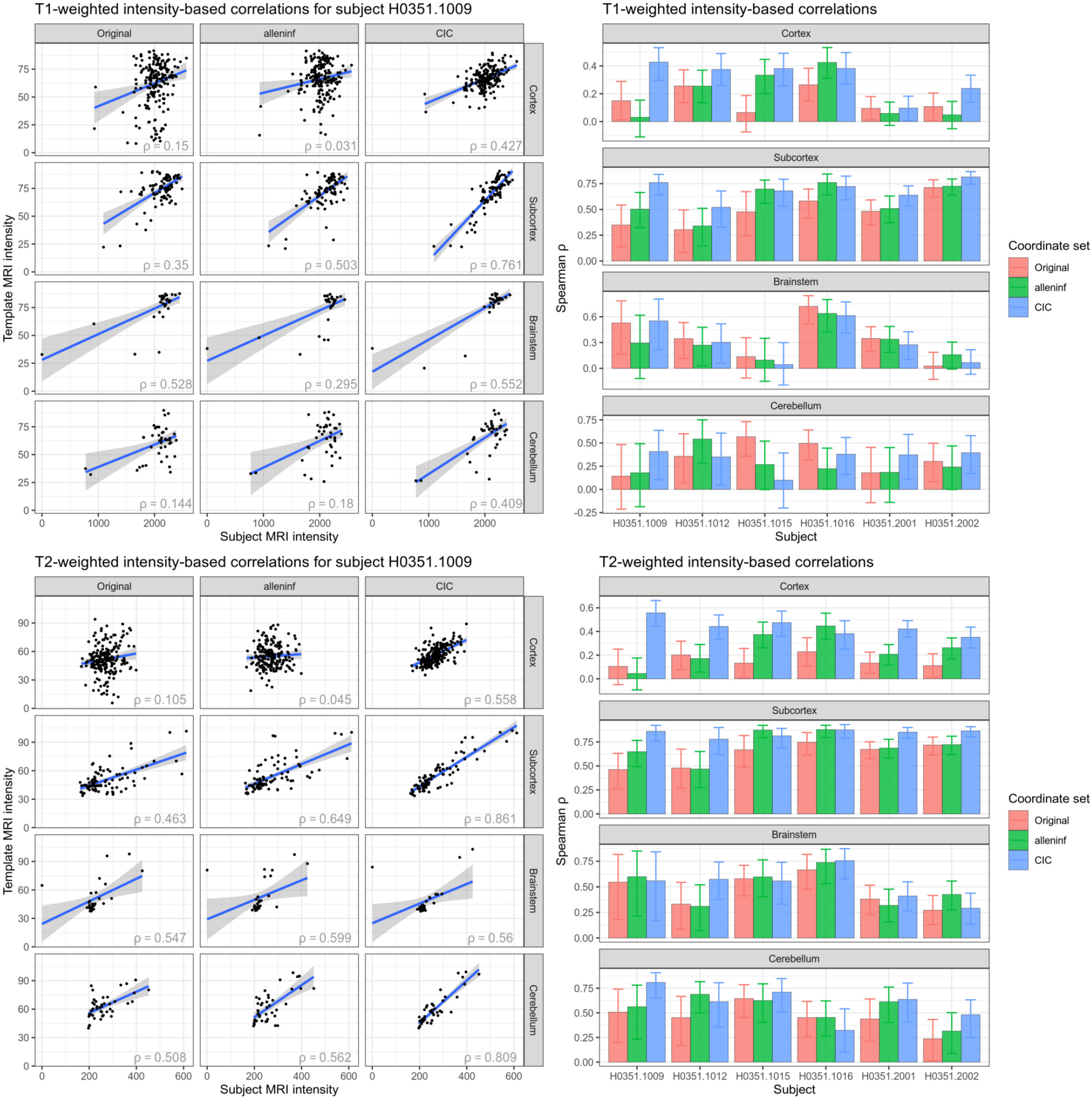
Intensity-based validation of registration quality examined within coarsely-defined substructures. Same as Figure 3, but carried out separately within large substructures of the brain, using samples dissected from the cortex, subcortex, brainstem, and cerebellum.

**Figure S4:**
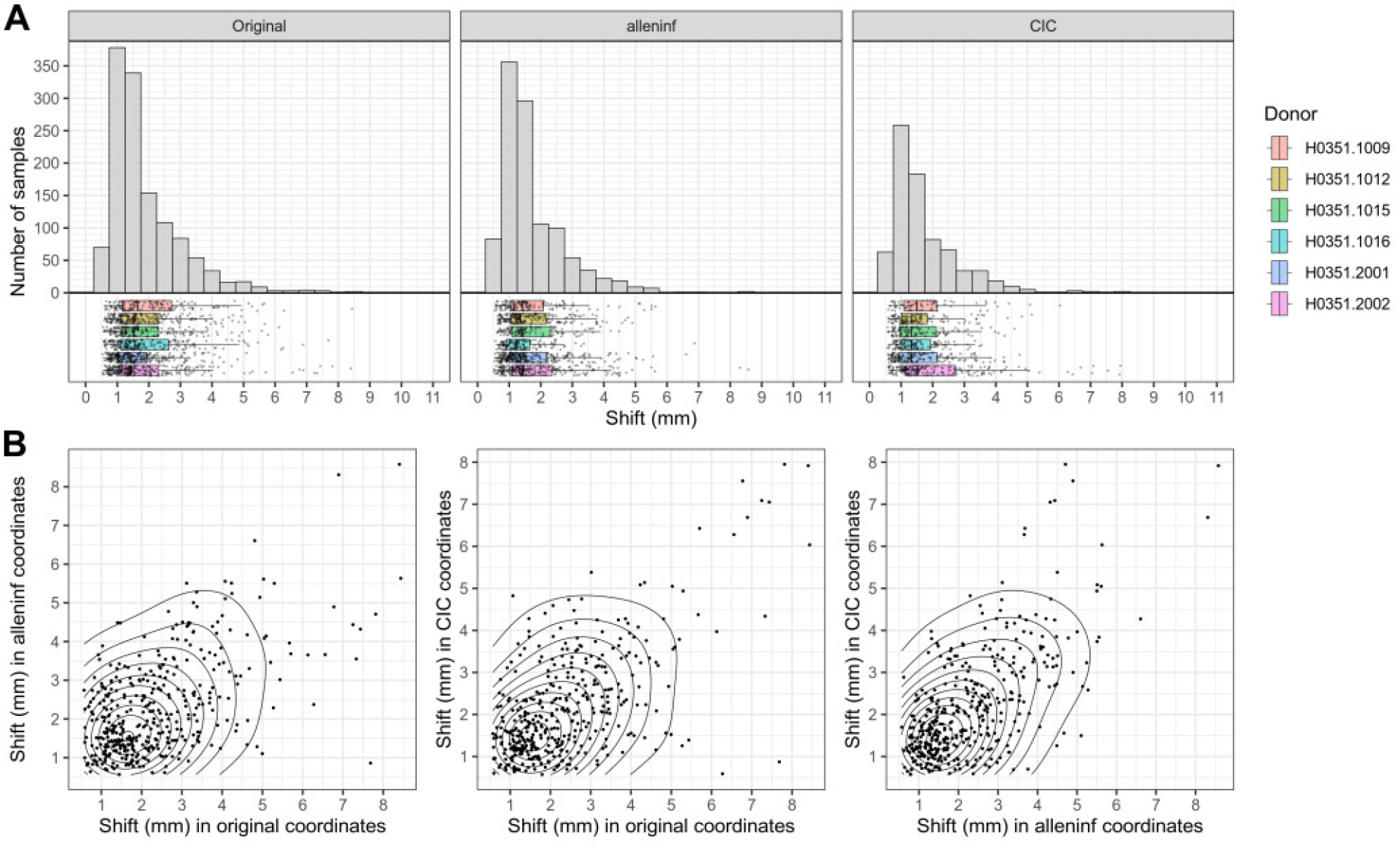
Shifts in sample coordinates after remapping samples to grey matter. (A) Distributions of distances shifted are shown for each of the three coordinate sets. (B) Distances are plotted against each pair of coordinate sets.

**Figure S5:**
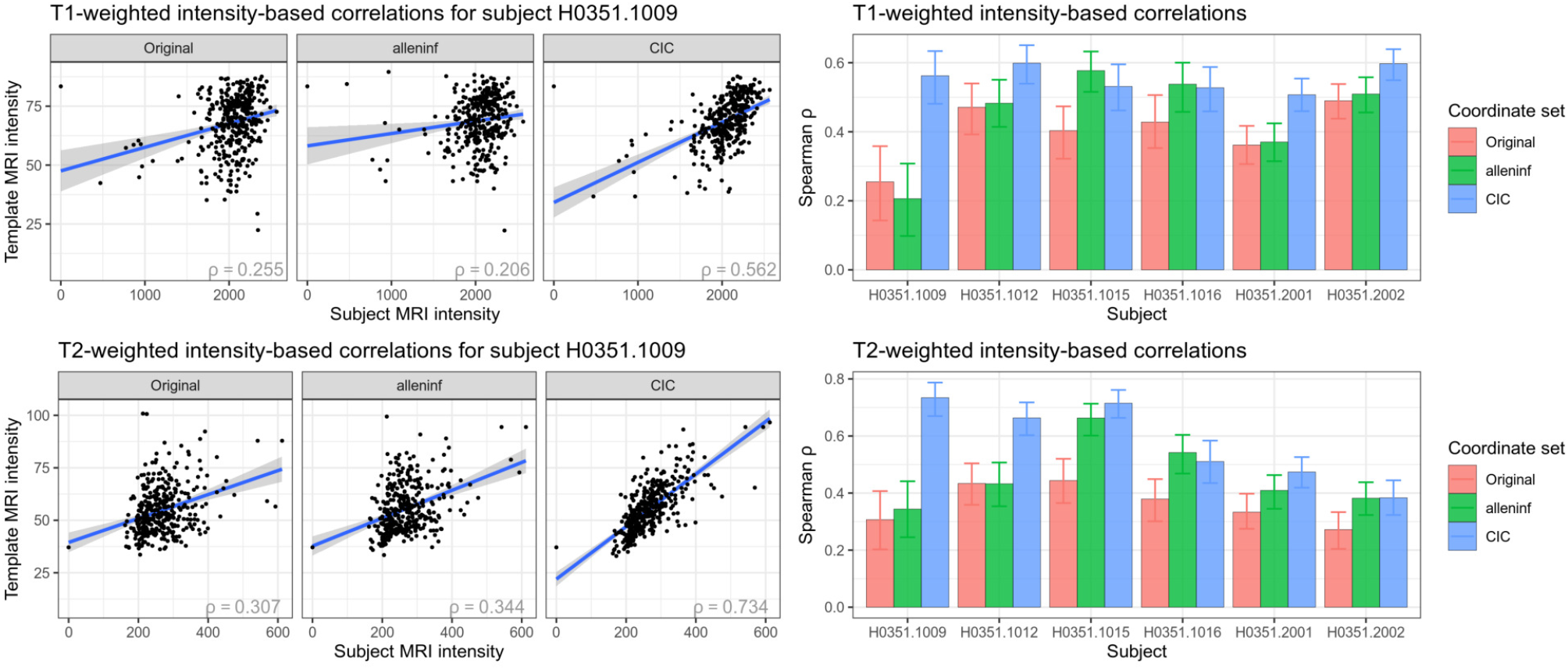
Intensity-based validation of registration quality using grey matter-remapped coordinates. Same as Figure 3, but using coordinates-based locations after shifting grey matter samples which were positioned in white matter or outside brain tissue to the nearest grey matter voxel.

**Figure S6:**
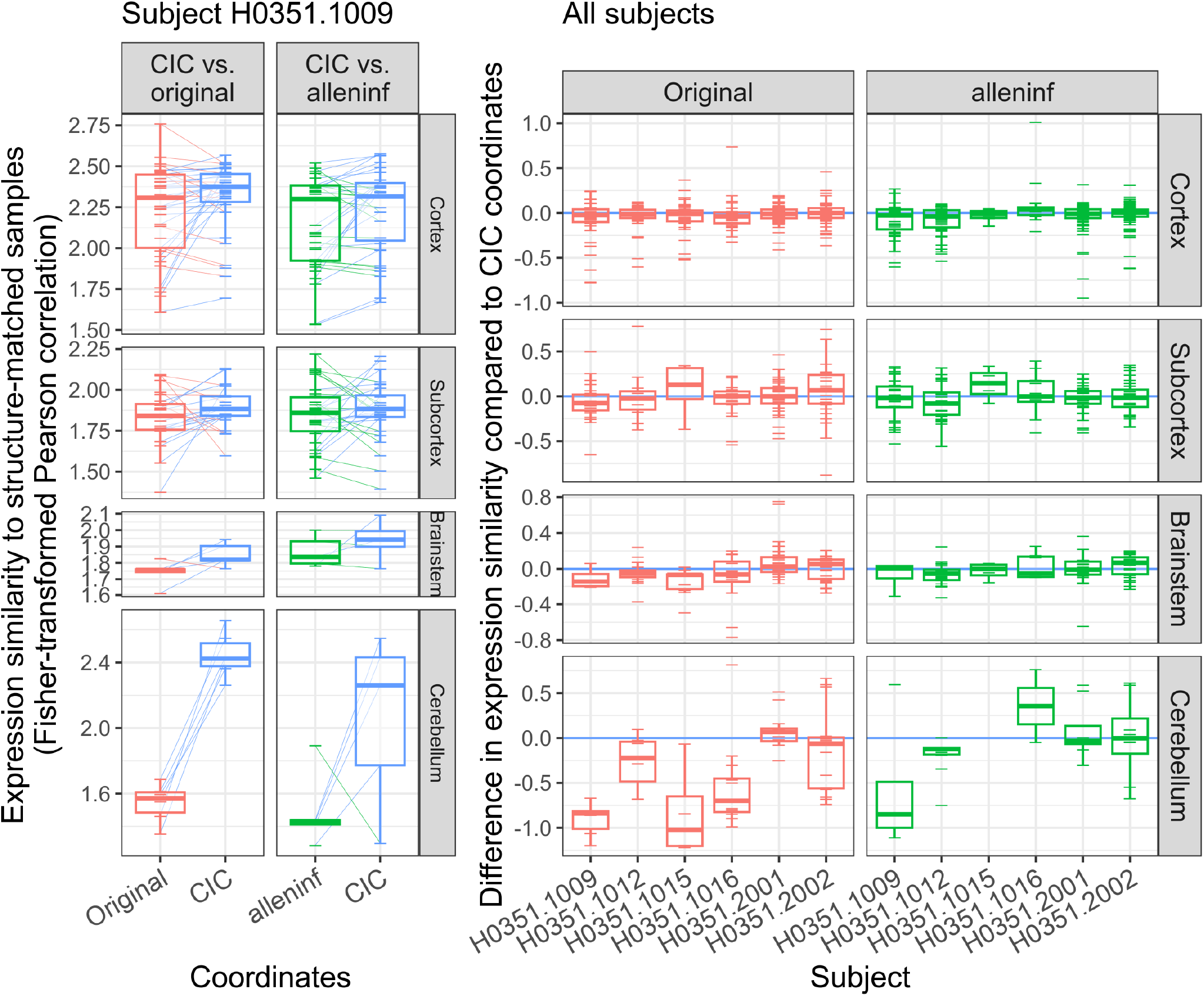
Expression similarity in individual subjects. Fisher-transformed correlation coefficients are compared between original and new (left column) and alleninf and new (right column) coordinate sets for one subject (left), with differences highlighted by lines joining the same samples. These differences are further shown for all subjects (right).

**Figure S7:**
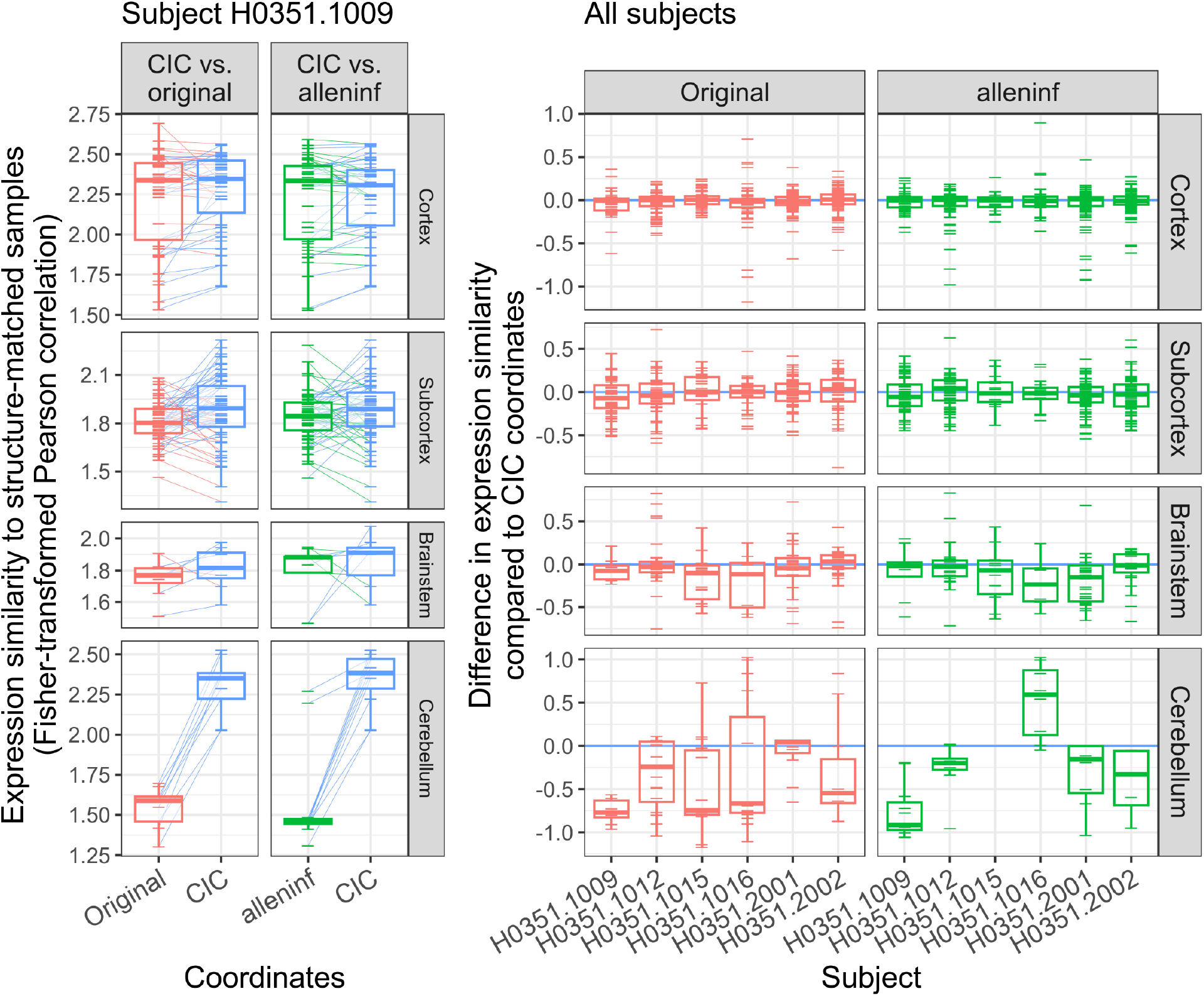
Expression similarity in individual subjects using grey matter-remapped coordinates. Same as Figure S6, but using coordinates-based locations after shifting grey matter samples which were positioned in white matter or outside brain tissue to the nearest grey matter voxel.

**Figure S8:**
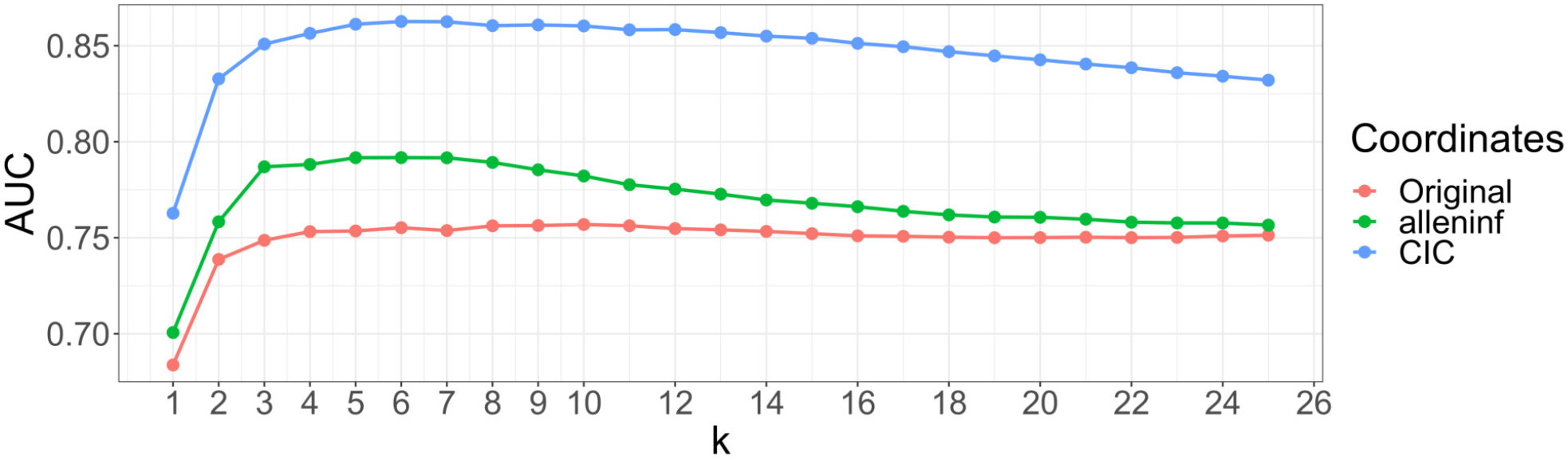
Area under the curve (AUC) as a function of *k*. The AUC reflects the extent that the interpolated *PVALB*/*CALB1* ratio overlaps with ground truth definitions, and is highest with CIC coordinates when using *k*-nearest neighbours interpolation for all *k*.

**Figure S9:**
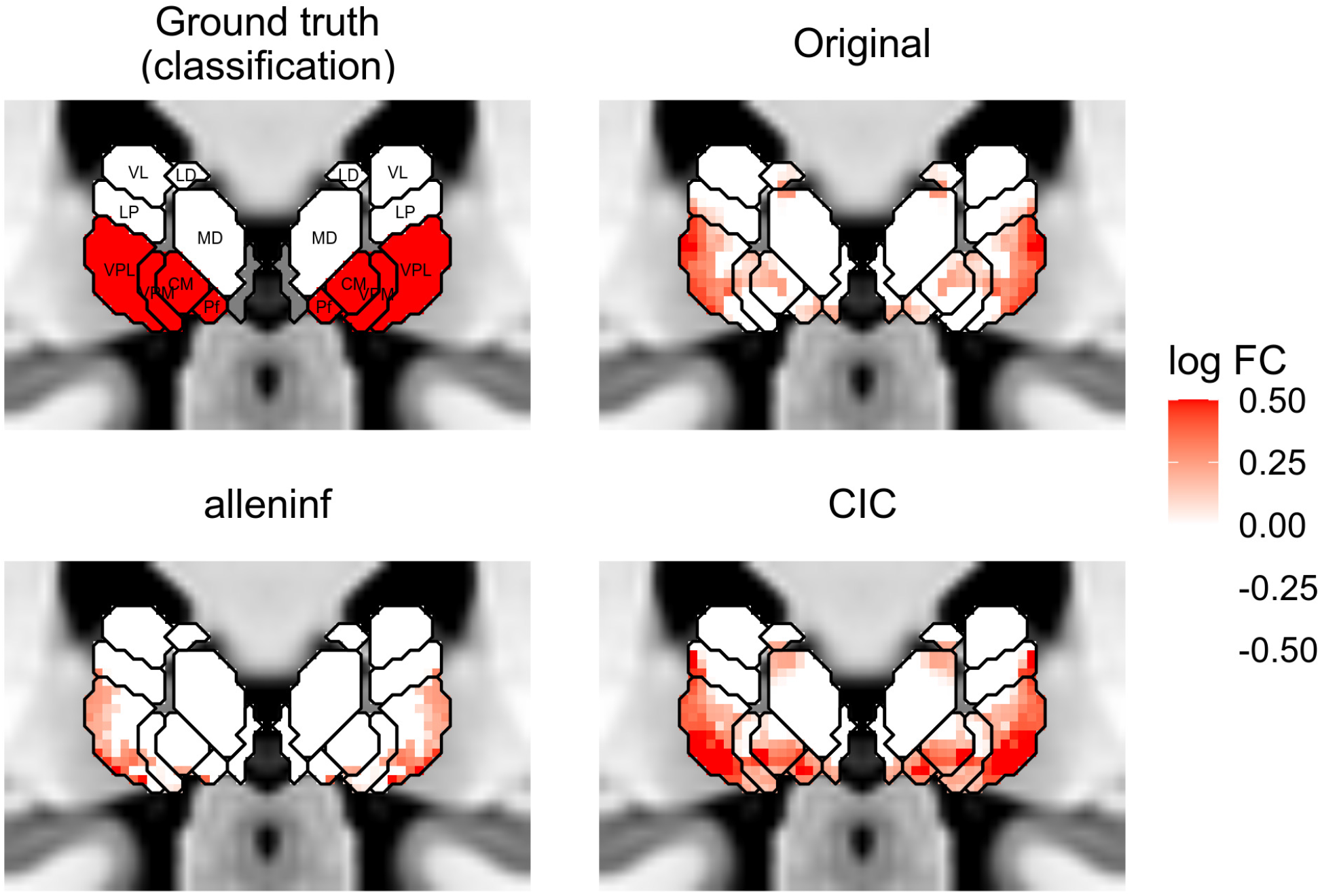
*PVALB*/*CALB1* interpolated using grey matter-remapped coordinates. Same as Figure 6D, but using coordinates-based locations after shifting grey matter samples which were positioned in white matter or outside brain tissue to the nearest grey matter voxel.

**Figure S10:**
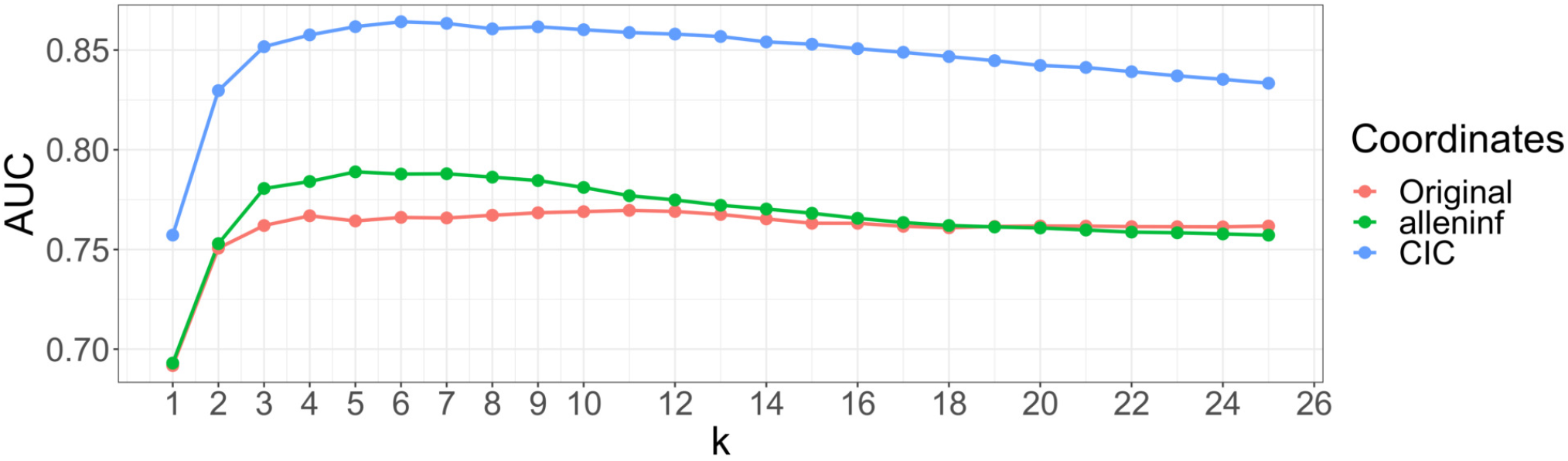
Area under the curve (AUC) as a function of k, using grey matter-remapped coordinates. Same as Figure S8, but using coordinates-based locations after shifting grey matter samples which were positioned in white matter or outside brain tissue to the nearest grey matter voxel.

**Figure S11:**
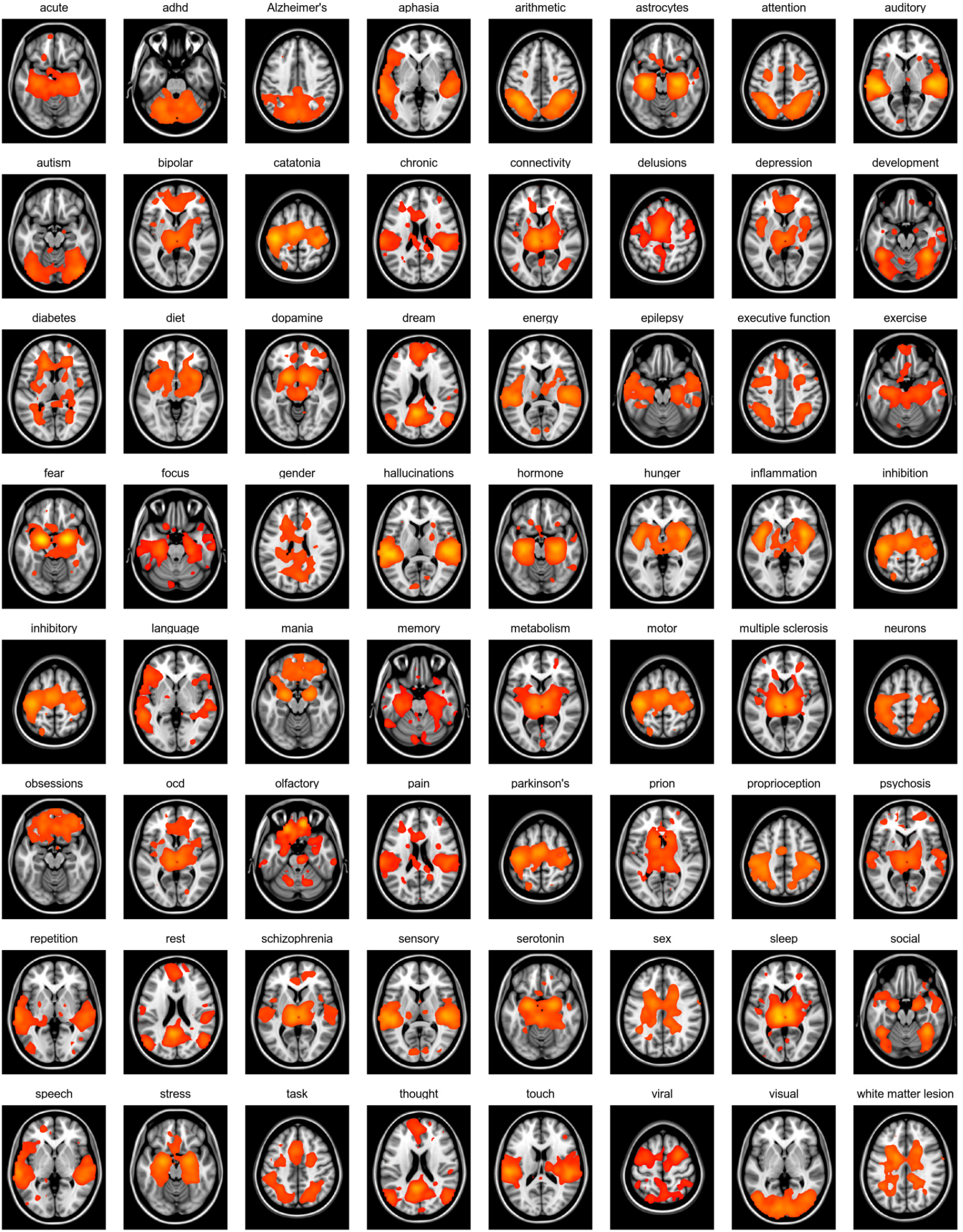
NeuroQuery maps used in this study. Axial slices show activation patterns associated with 64 of 66 NeuroQuery maps used in this study.

**Figure S12:**
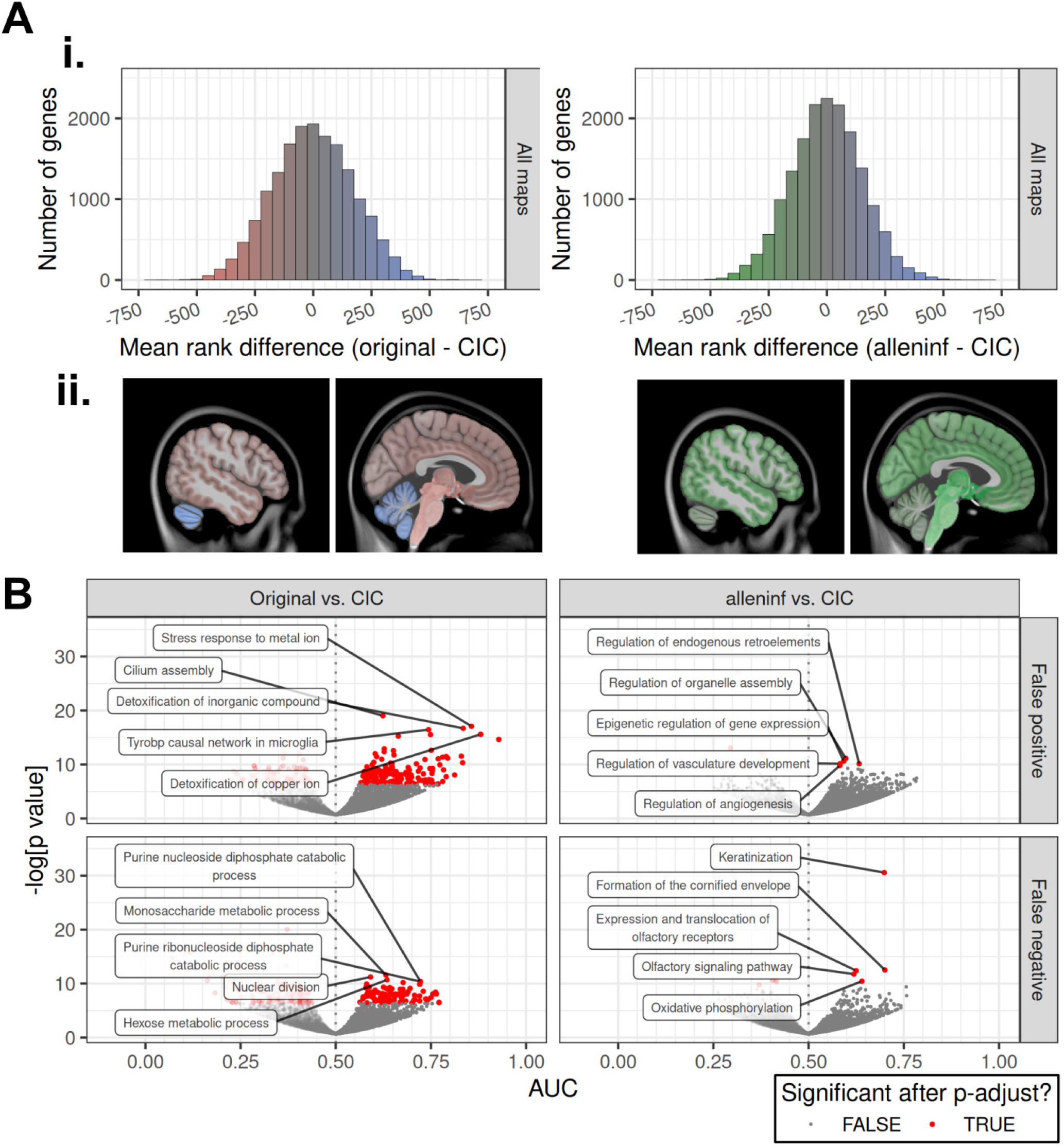
Gene rank differences in analyses of NeuroQuery maps due to coordinate choice. (A) Rank differences are averaged over all maps in (i), and are plotted over the brain after weighting by regional gene expression in (ii). (B) The gene ontology and pathway modules associated with most biased genes (i.e. those at the tail end of the distributions in (A) are displayed through volcano plots.

**Figure S13:**
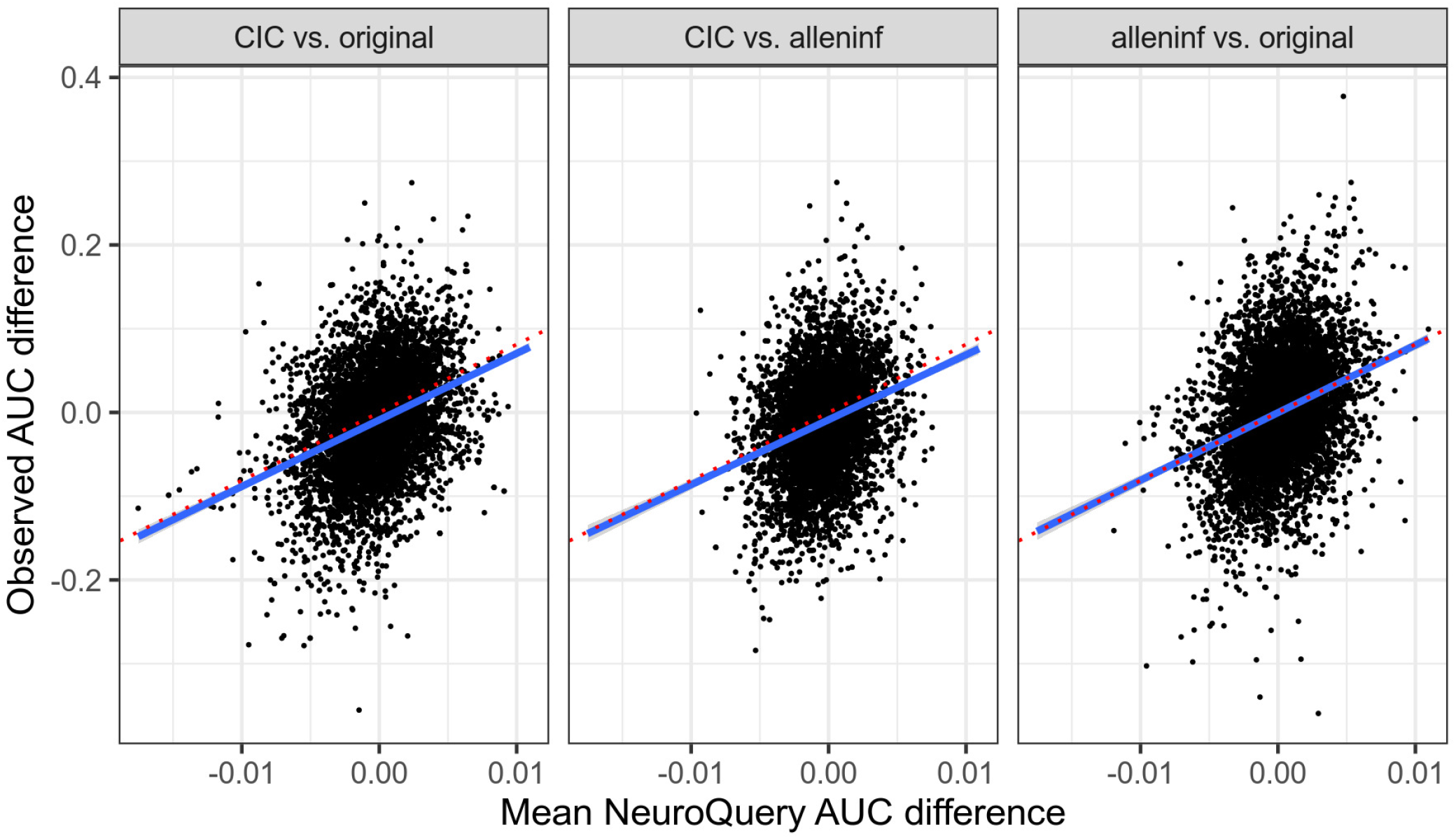
Comparison of predicted bias with observed bias, validating predictions made in the NeuroQuery analyses. Predicted bias (computed over all NeuroQuery maps) tracks with observed bias in the Patel et al. (*27*) study.

**Figure S14:**
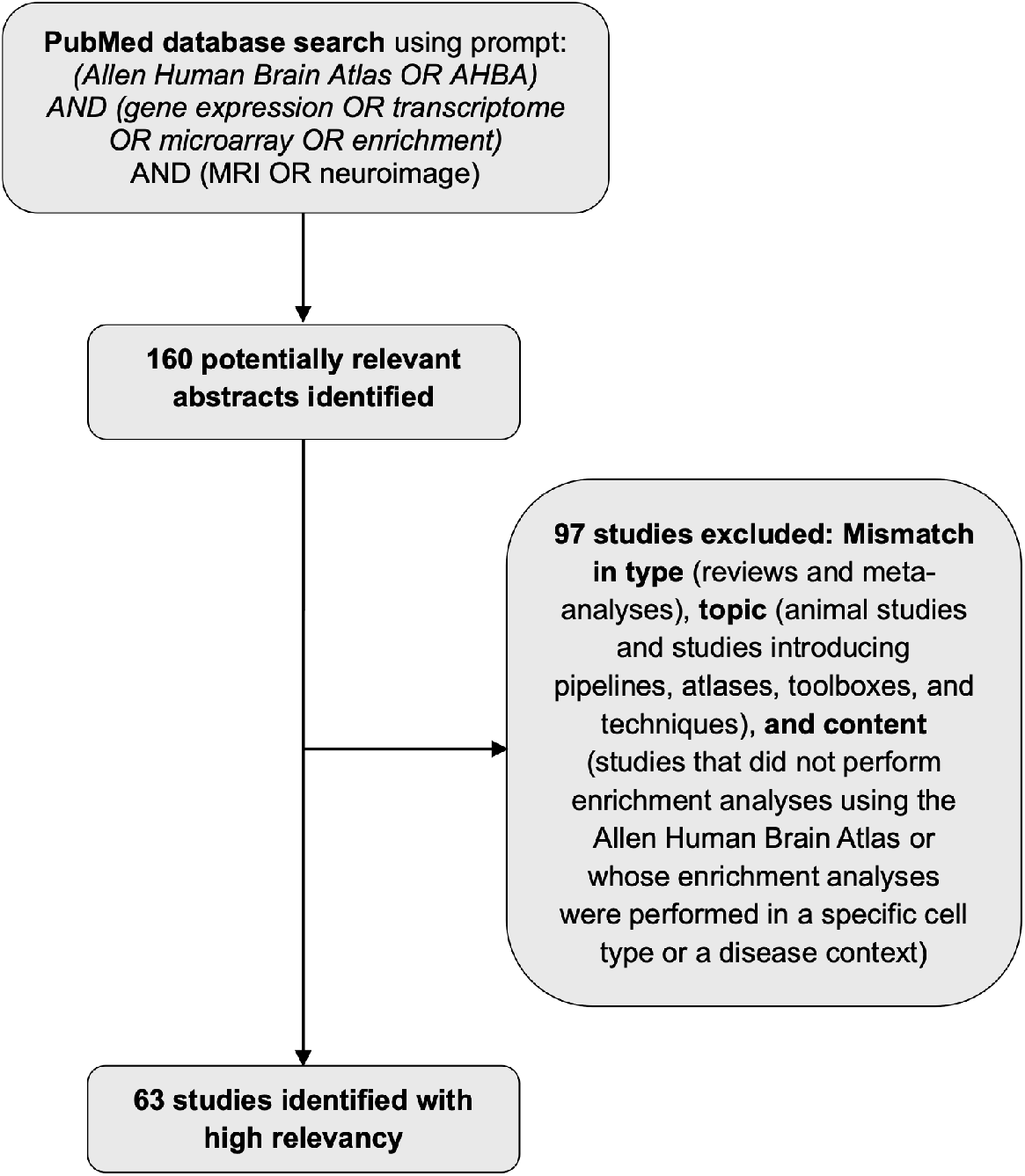
Selection of a representative sample of imaging-transcriptomic studies. Flow-chart depicting selection of imaging-transcriptomic studies examined for their use of AHBA sample coordinates.

**Figure S15:**
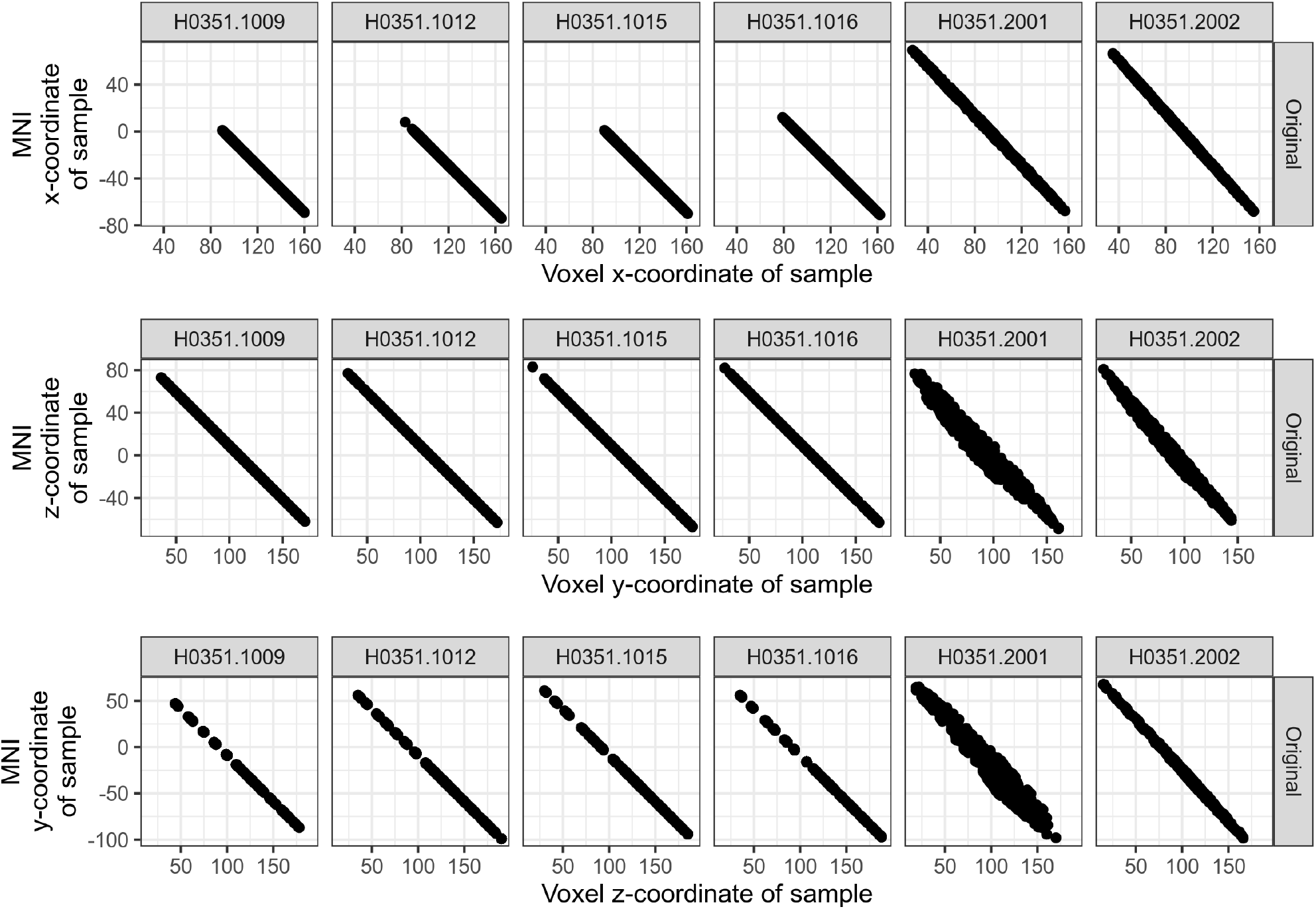
Affine-only alignment within original coordinates. For subject (columns), MRI *x*-, *y*-, and *z*-coordinates (rows) of tissue samples are plotted against originally reported MNI coordinates. An entirely linear relationship indicates is evidence of an affine-only transformation that maps MRI coordinates to MNI-space, and is seen for the four brains acquired *ex cranio* from UCI.

**Figure S16:**
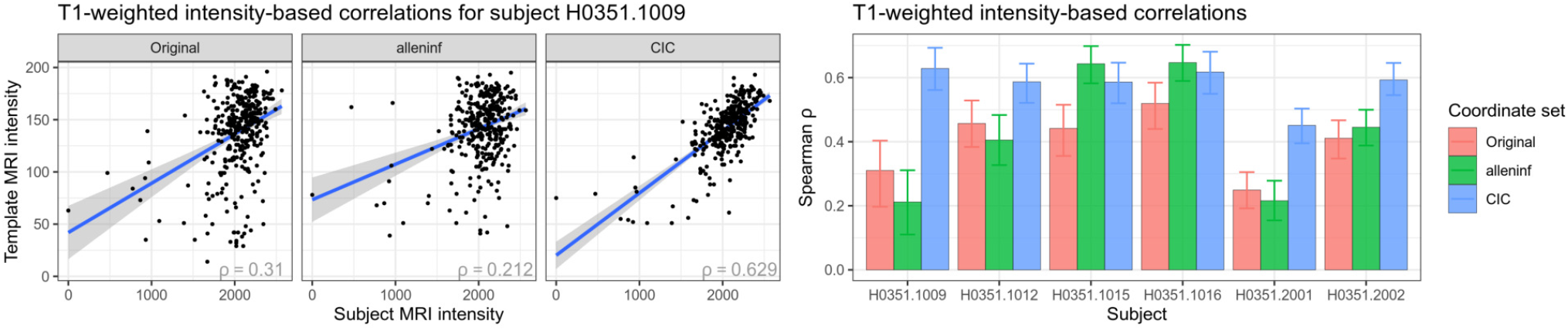
Intensity-based validation of registration quality using the MNI152 6th generation nonlinear asymmetric template. Same as Figure 3A, but with template intensity data derived from the MNI152 6th generation nonlinear asymmetric (“nlin6asym”) template instead of the MNI ICBM152 nonlinear symmetric 2009c. CIC coordinates were transformed to the nlin6asym template prior to analysis.

**Figure S17:**
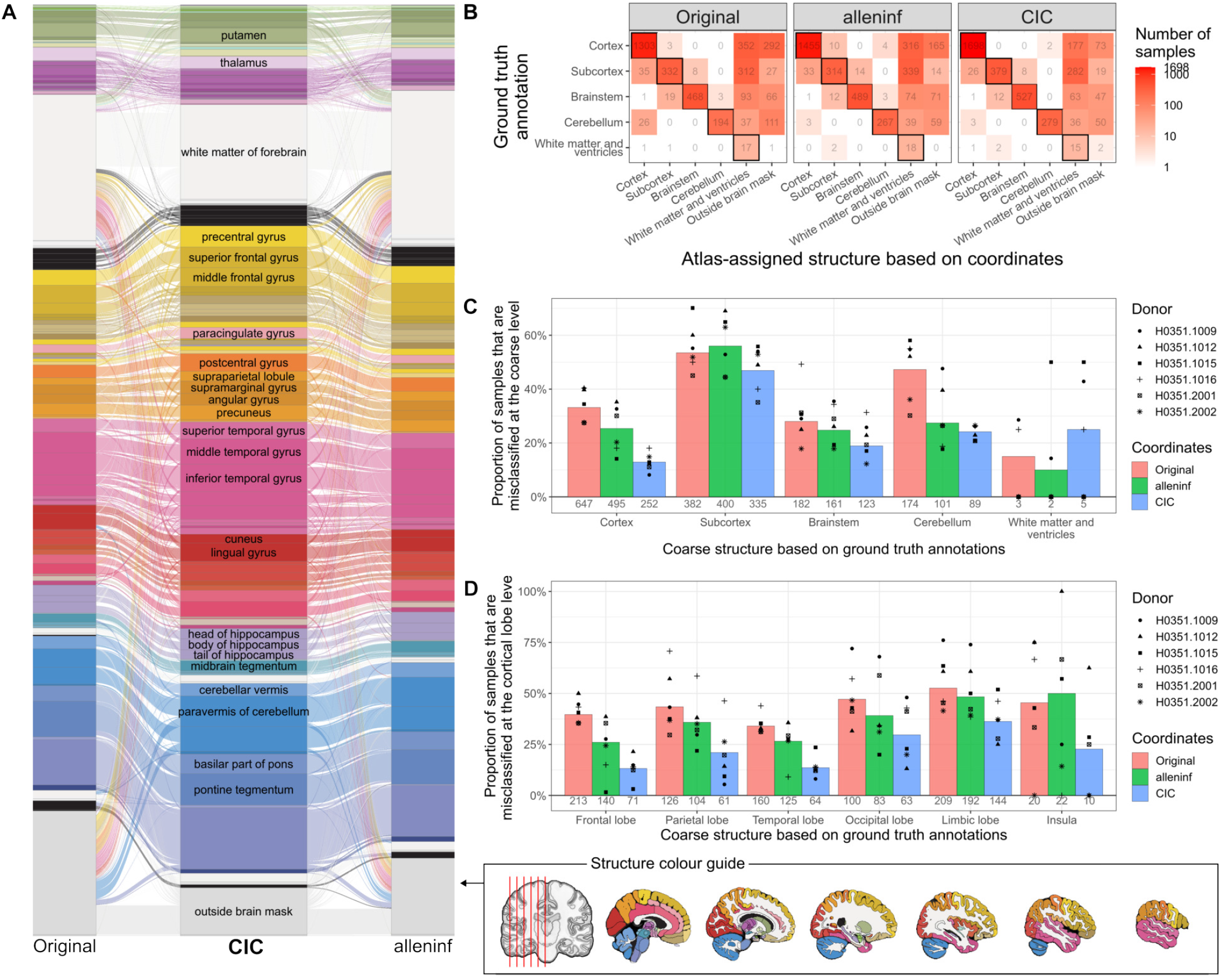
Assignment of samples to structures based on coordinate locations within the Allen Institute’s Human Reference Atlas (AHRA) relative to the nlin6asym template. Same as Figure 4, but with sample coordinates located relative to the nlin MNI152 6th generation nonlinear asymmetric (“nlin6asym”) template. CIC coordinates were transformed to the nlin6asym template prior to analysis.

**Figure S18:**
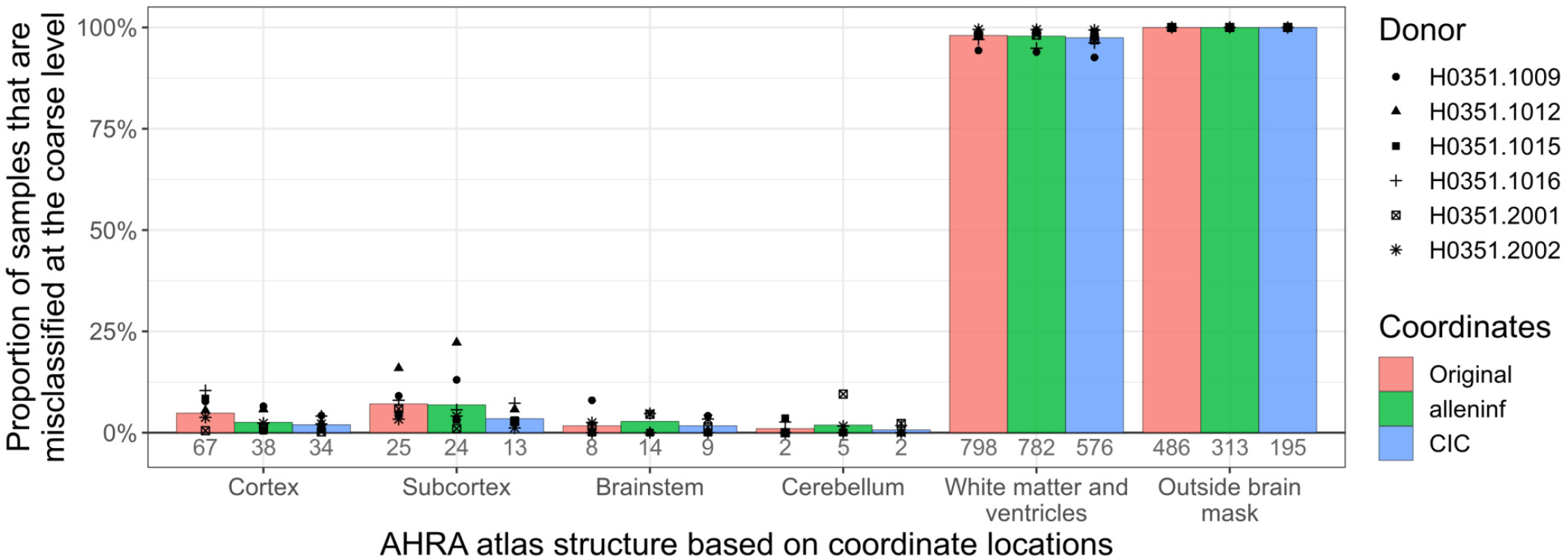
False discovery rate when predicting the coarse structure from which samples were dissected when using coordinate-based positioning within the AHRA (relative to the sym09c template). Same as Figure 4C, but with mislabeling quantified as the false discovery rate (1-PPV).

**Figure S19:**
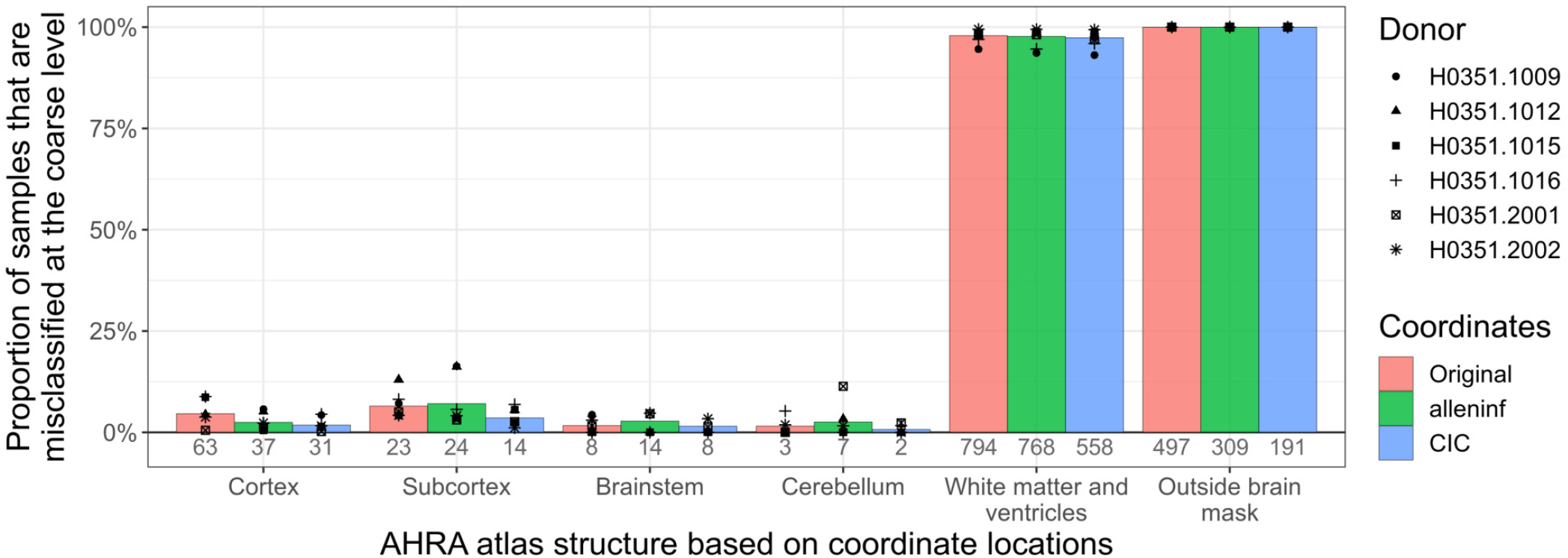
False discovery rate when predicting the coarse structure from which samples were dissected when using coordinate-based positioning within the AHRA (relative to the nlin6asym template). Same as Figure 4C, but with mislabeling quantified as the false discovery rate (1-PPV), and using sample coordinates located relative to the nlin MNI152 6th generation nonlinear asymmetric (“nlin6asym”) template. CIC coordinates were transformed to the nlin6asym template prior to analysis.

**Figure S20:**
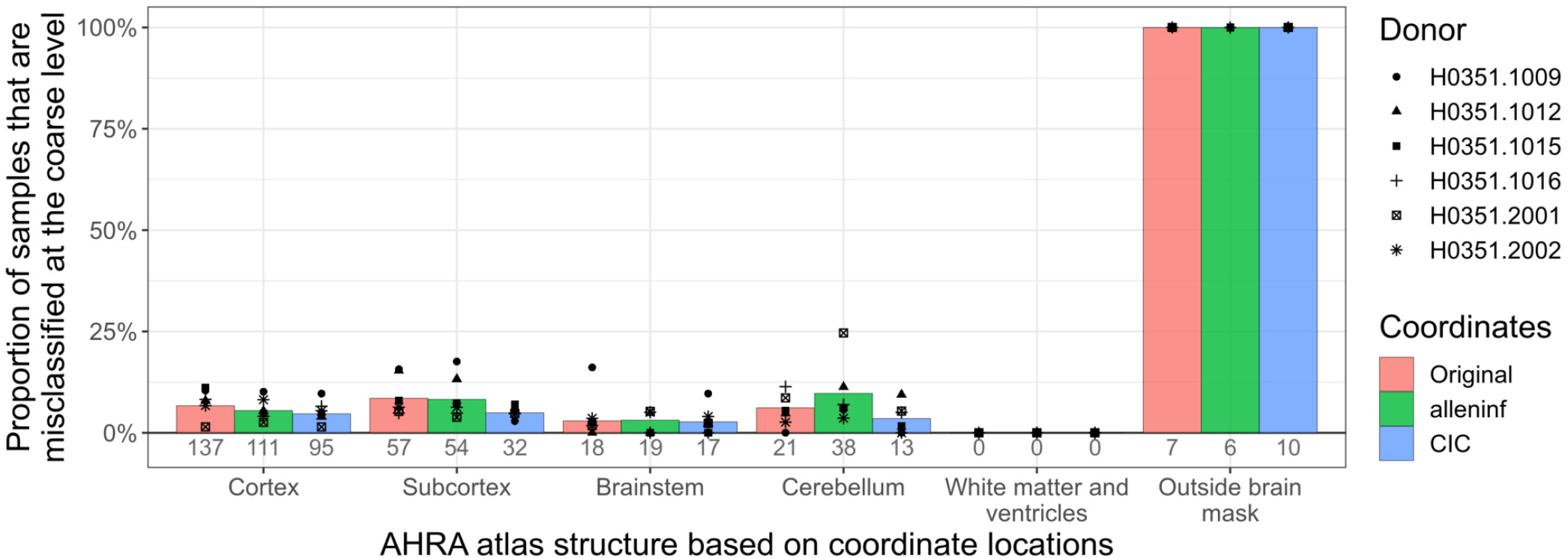
False discovery rate when predicting the coarse structure from which samples were dissected, using grey matter-remapped coordinates. Same as Figure 4C, but with mislabeling quantified as the false discovery rate (1-PPV), and using coordinates-based locations after shifting grey matter samples which were positioned in white matter or outside brain tissue to the nearest grey matter voxel.

**Figure S21:**
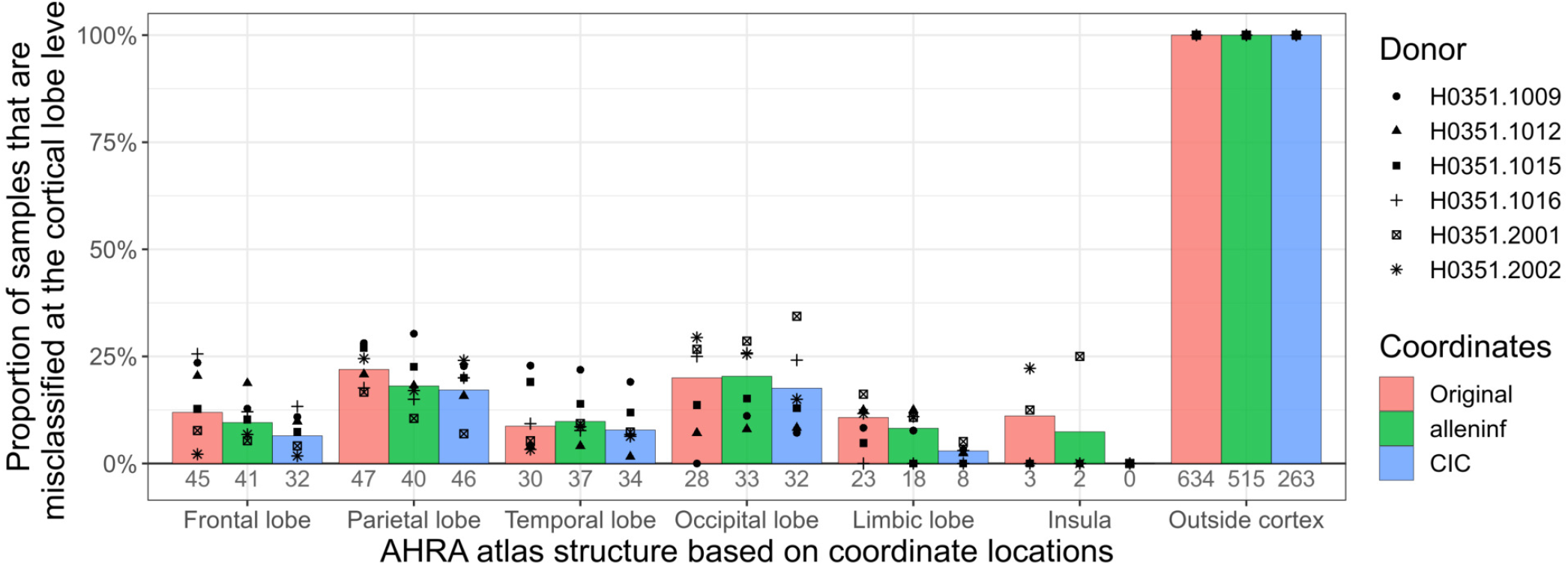
False discovery rate when predicting the cortical lobe from which cortical samples were dissected when using coordinate-based positioning within the AHRA (relative to the sym09c template). Same as Figure 4D, but with mislabeling quantified as the false discovery rate (1-PPV).

**Figure S22:**
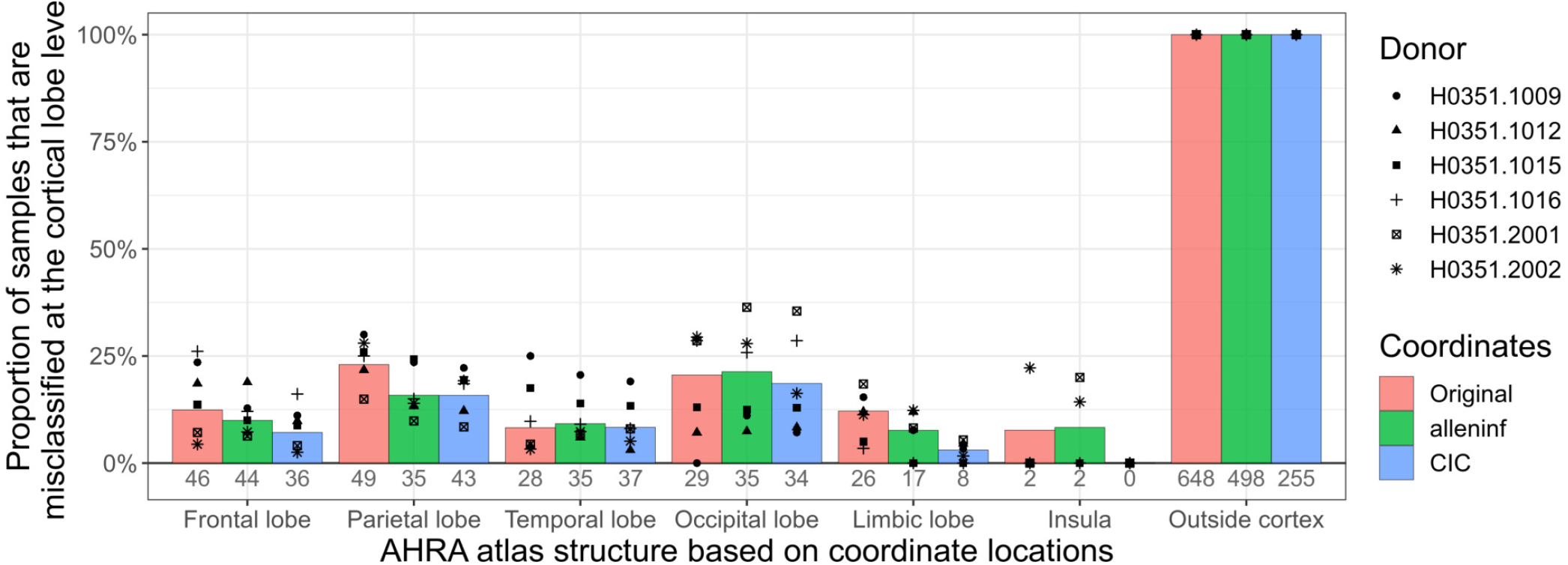
False discovery rate when predicting the cortical lobe from which cortical samples were dissected when using coordinate-based positioning within the AHRA (relative to the nlin6asym template). Same as Figure 4D, but with mislabeling quantified as the false discovery rate (1-PPV), and using sample coordinates located relative to the nlin MNI152 6th generation nonlinear asymmetric (“nlin6asym”) template. CIC coordinates were transformed to the nlin6asym template prior to analysis.

**Figure S23:**
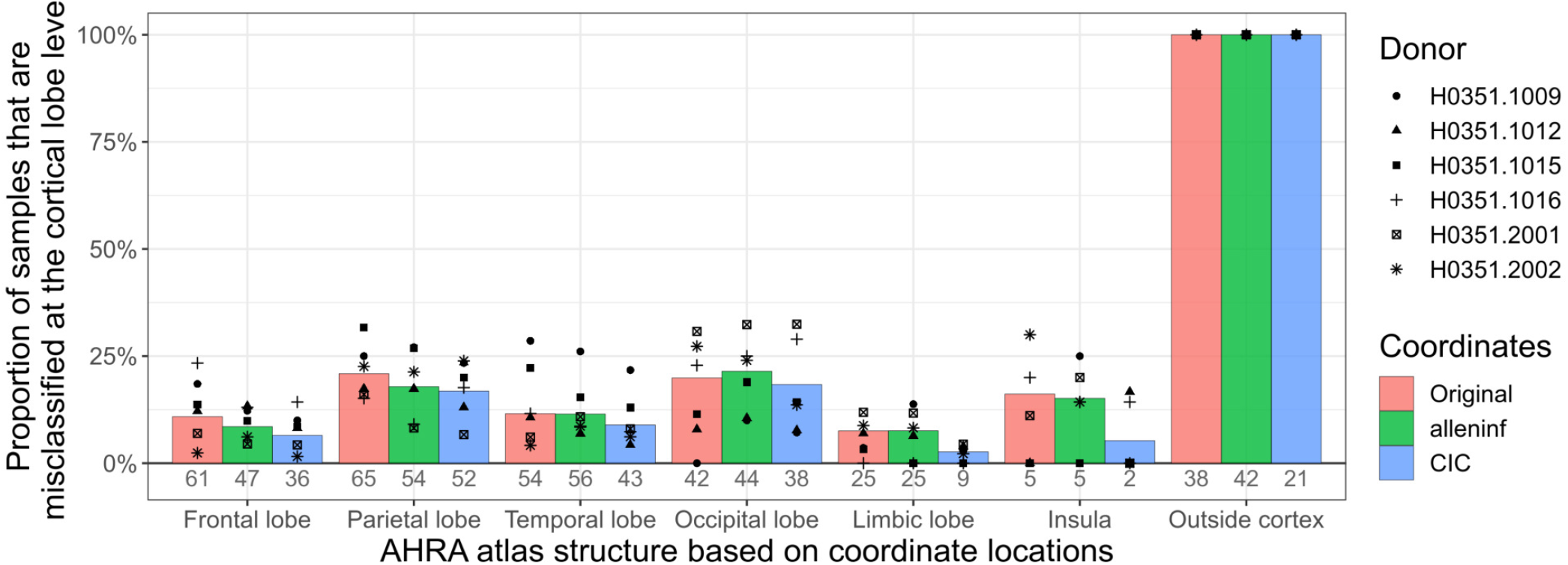
False discovery rate when predicting the cortical lobe from which cortical samples were dissected, using grey matter-remapped coordinates. Same as Figure 4D, but with mislabeling quantified as the false discovery rate (1-PPV), and using coordinates-based locations after shifting grey matter samples which were positioned in white matter or outside brain tissue to the nearest grey matter voxel.

**Figure S24:**
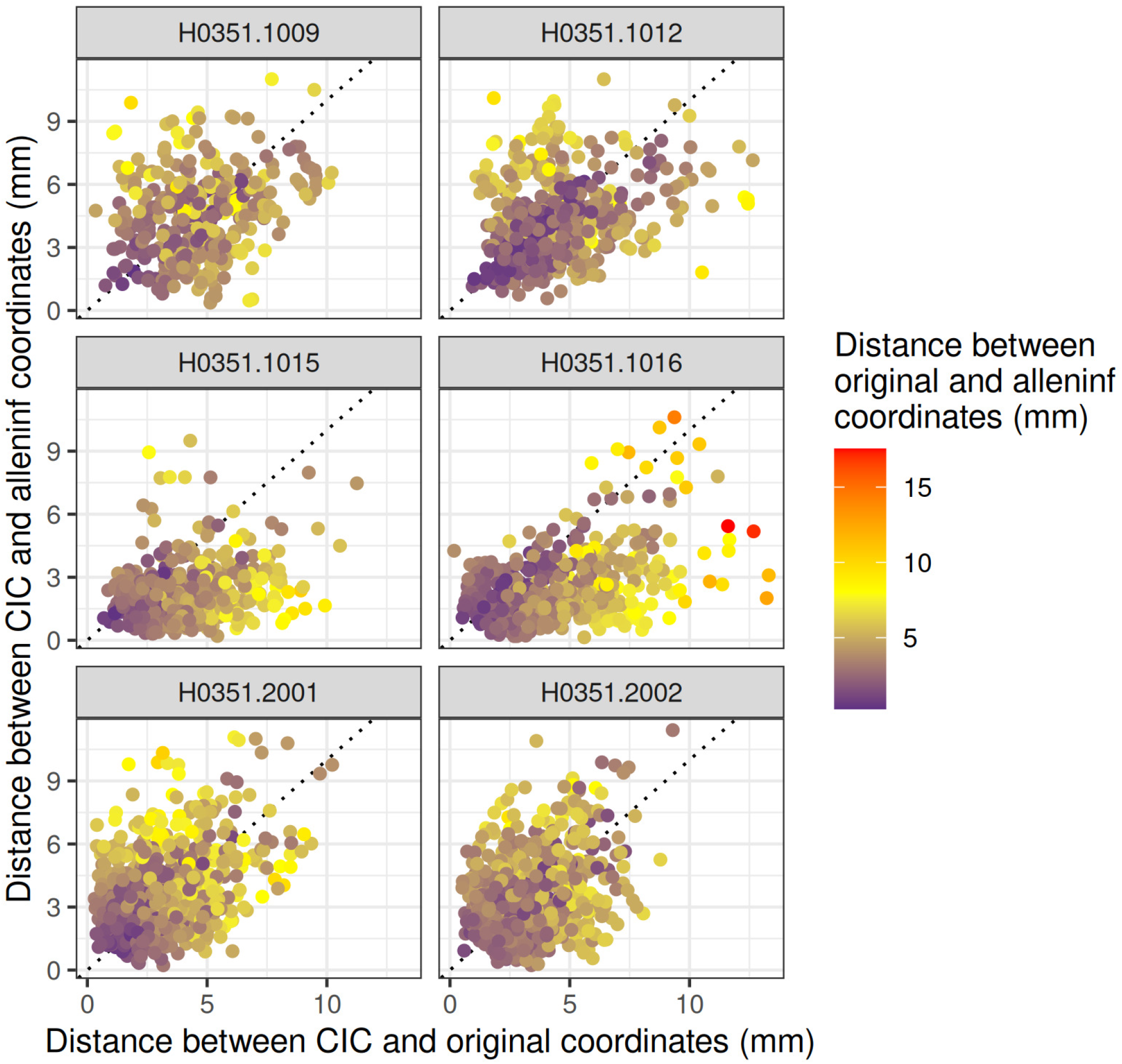
Similarity between coordinate representations. Distances between MNI-space locations of samples given by CIC and original coordinates are plotted against distances between MNI-space locations given by CIC and alleninf coordinates. Distances between locations given by alleninf and original coordinates are represented by colours. Each point is a sample, and data are separated by subject.

**Figure S25:**
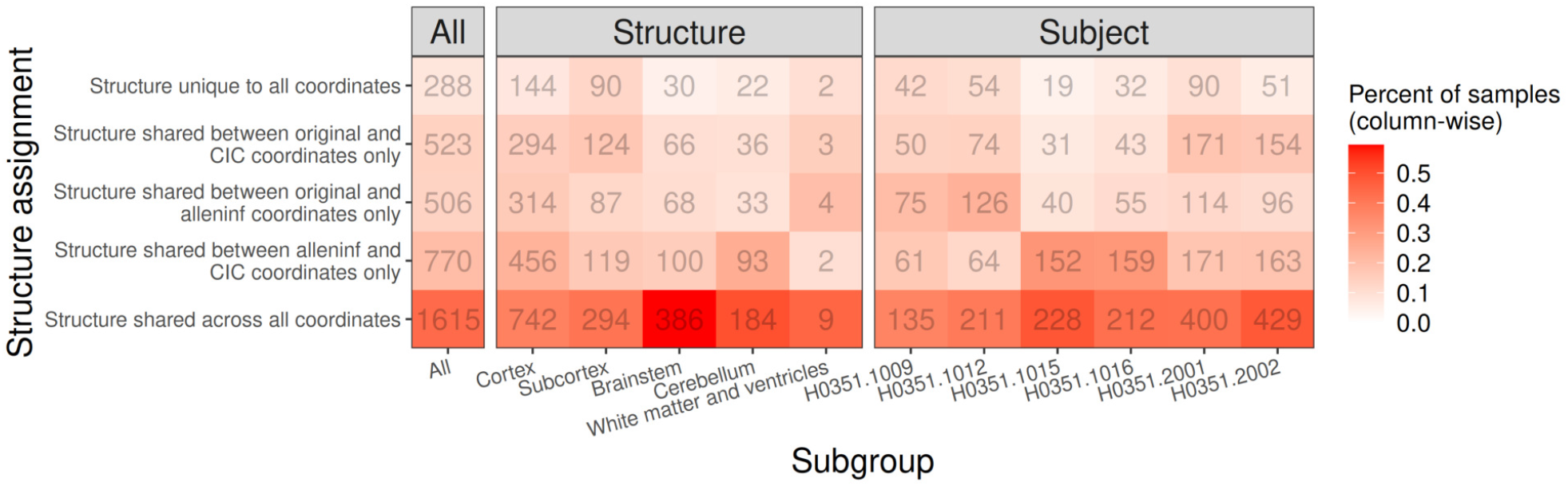
Concordance between different coordinate representations in their placement of samples within coarse neuroanatomical structures (defined by AHRA relative to the sym09c template). The number of samples that are placed in the same coarse structure according to different sets of coordinate representations (rows) are listed for all samples, separated by true dissection structure, and by subject. Colours represent proportions of samples computed within each subgroup of samples (i.e. column-wise).

**Figure S26:**
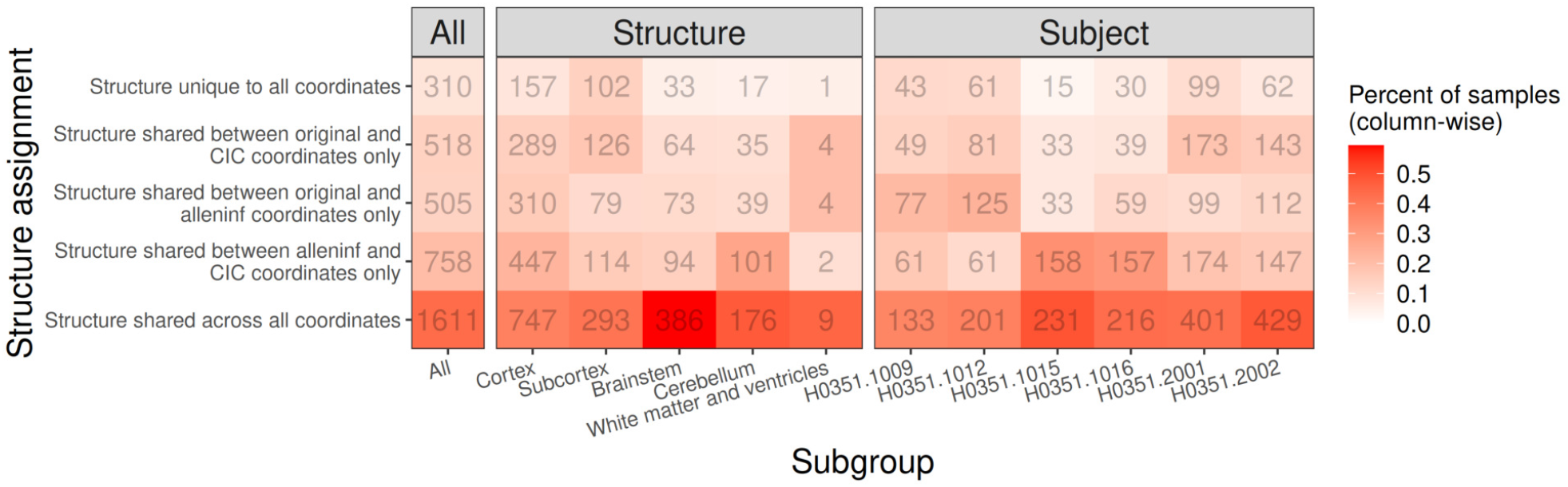
Concordance between different coordinate representations in their placement of samples within coarse neuroanatomical structures (defined by AHRA relative to the nlin6asym template). Same as Figure S25, but with sample coordinates located relative to the nlin MNI152 6th generation nonlinear asymmetric (“nlin6asym”) template. CIC coordinates were transformed to the nlin6asym template prior to analysis.

**Figure S27:**
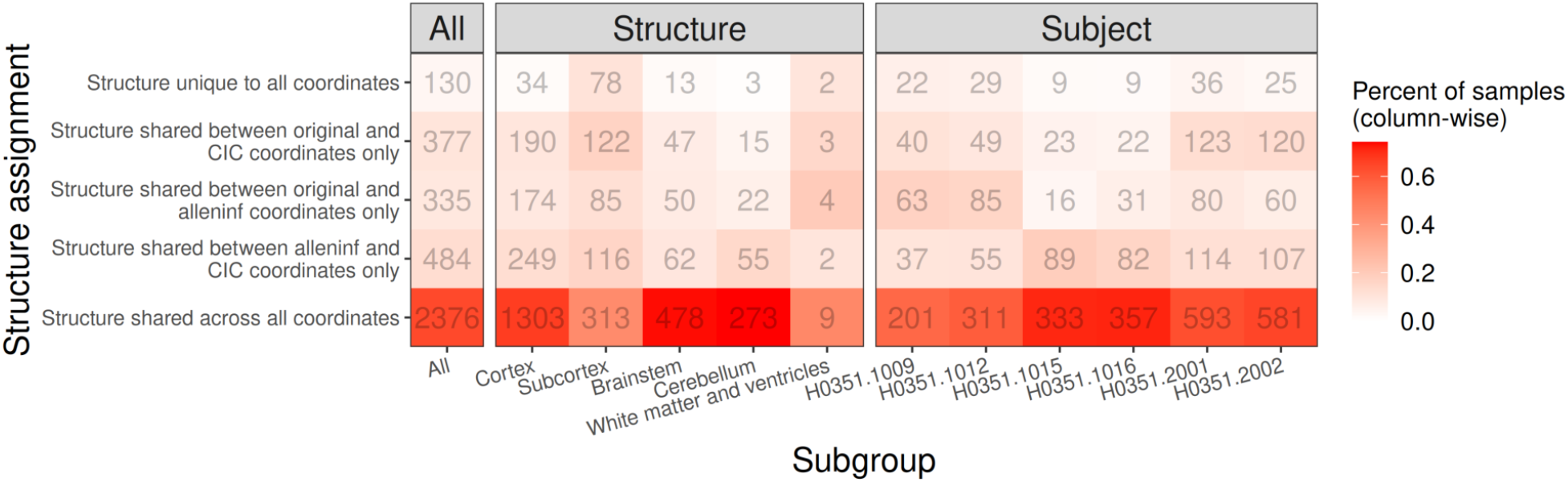
Concordance between different coordinate representations in their placement of samples within coarse neuroanatomical structures, using grey matter-remapped coordinates. Same as Figure S25, but using coordinates-based locations after shifting grey matter samples which were positioned in white matter or outside brain tissue to the nearest grey matter voxel.

**Figure S28:**
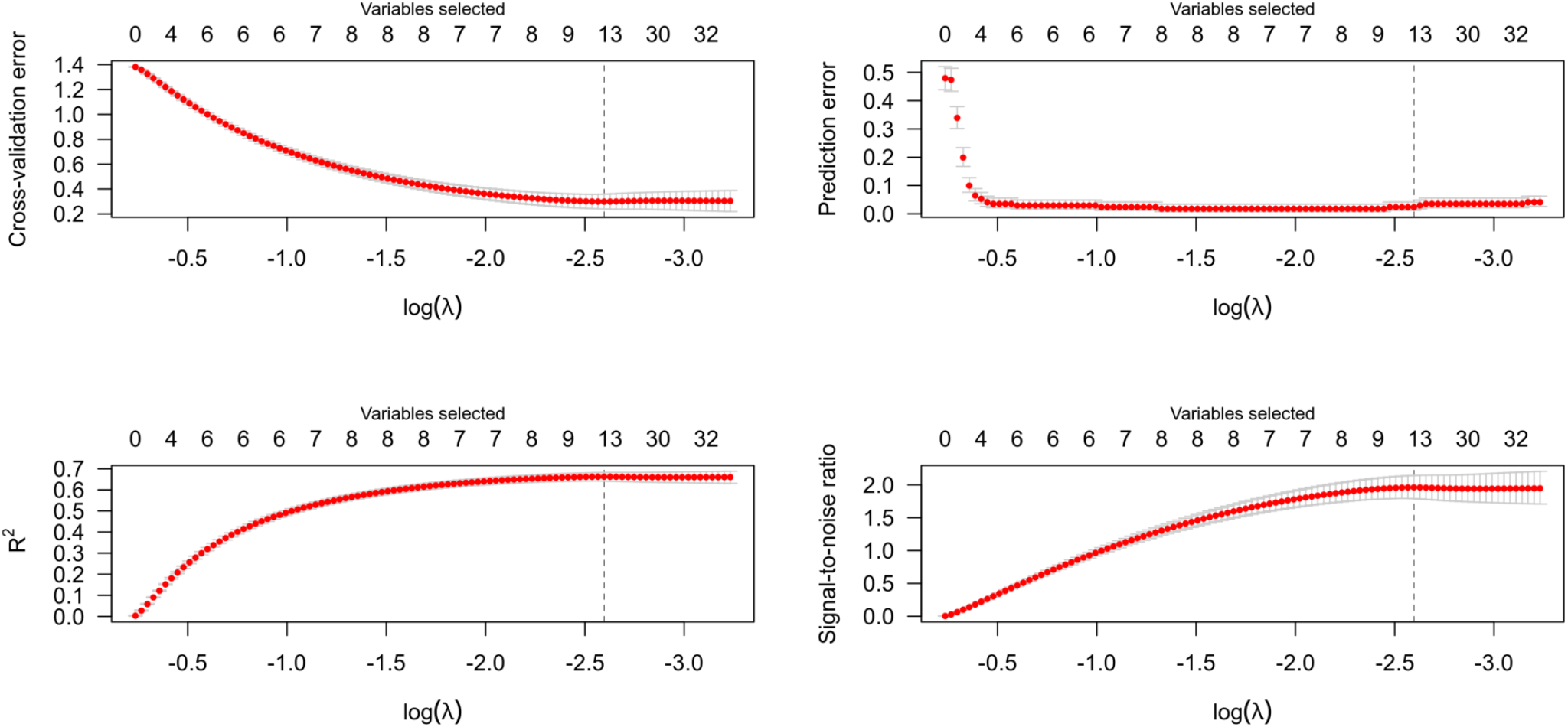
LASSO model fit and variable selection results. LASSO prediction of samples’ structure assignment given sample gene expression. Tissue samples dissected from only the precentral and postcentral gyrus were selected, and a sparse set of marker genes (that optimally predicted which of the two structures each sample was dissected from) were determined by minimizing cross-validation error.

**Figure S29:**
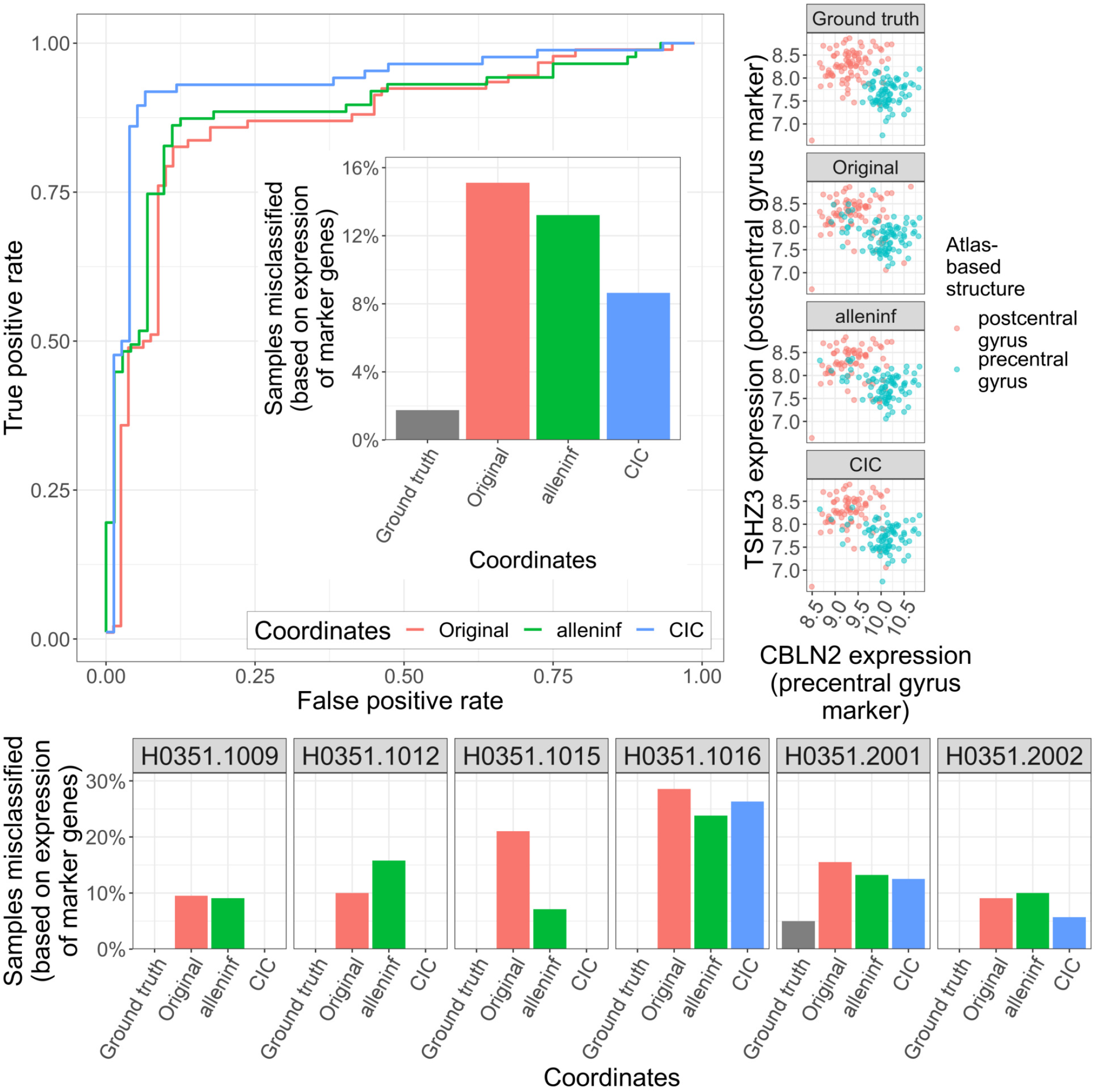
Expression-based separation analysis for precentral gyrus vs. postcentral gyrus. ROC curve demonstrating the extent that coordinate-based structure assignment can classify samples into their gene expression-based structure assigment (top left), along with the proportion of samples misclassified (inset) and separated by subject (bottom). A visualization of the ability of the top two genes to separate samples based on their AHBA coordinates are shown on the top right.

**Figure S30:**
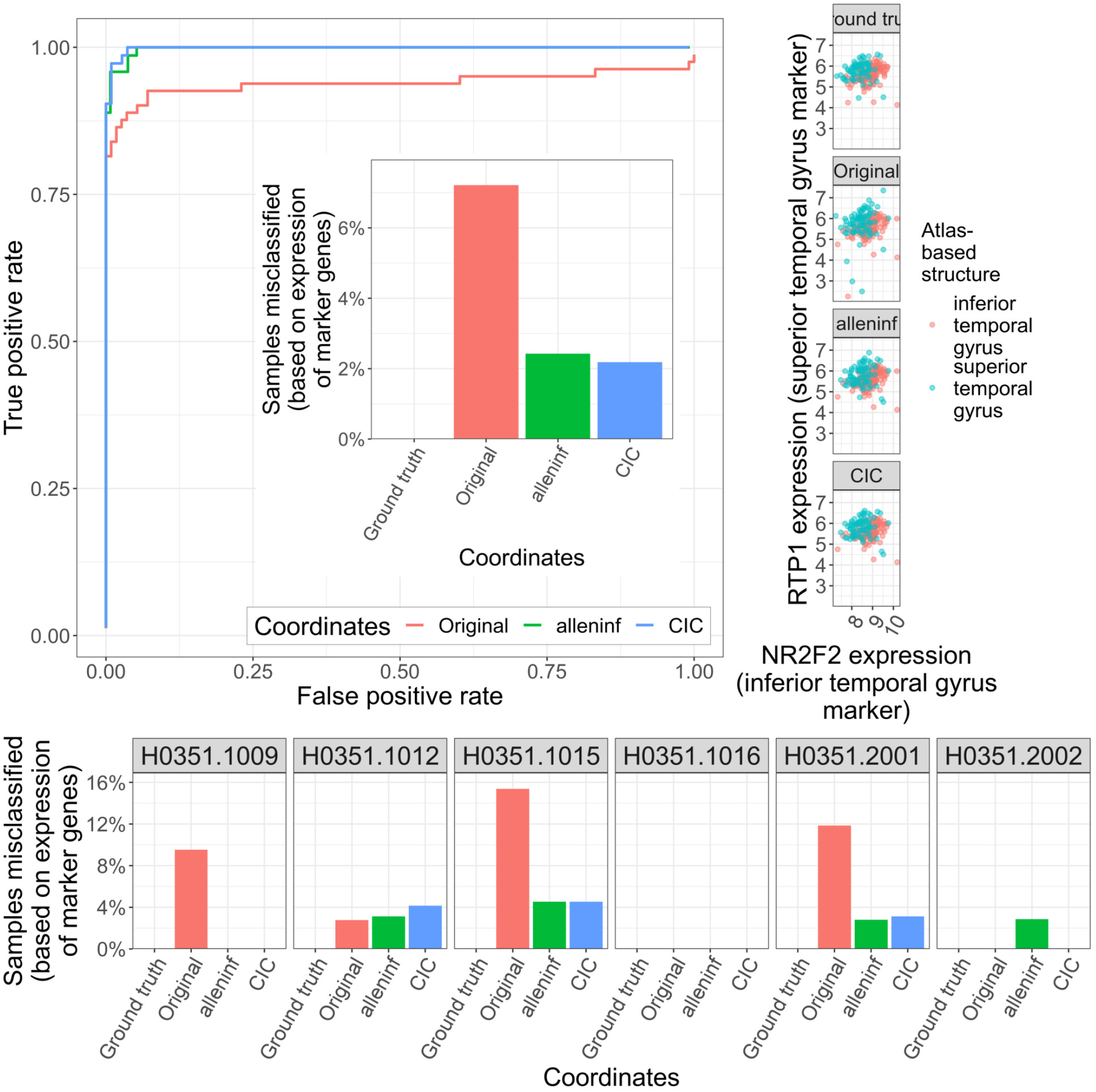
Expression-based separation analysis for superior temporal gyrus vs. inferior temporal gyrus. Same as Figure S29, but comparing the superior temporal gyrus vs. inferior temporal gyrus.

**Figure S31:**
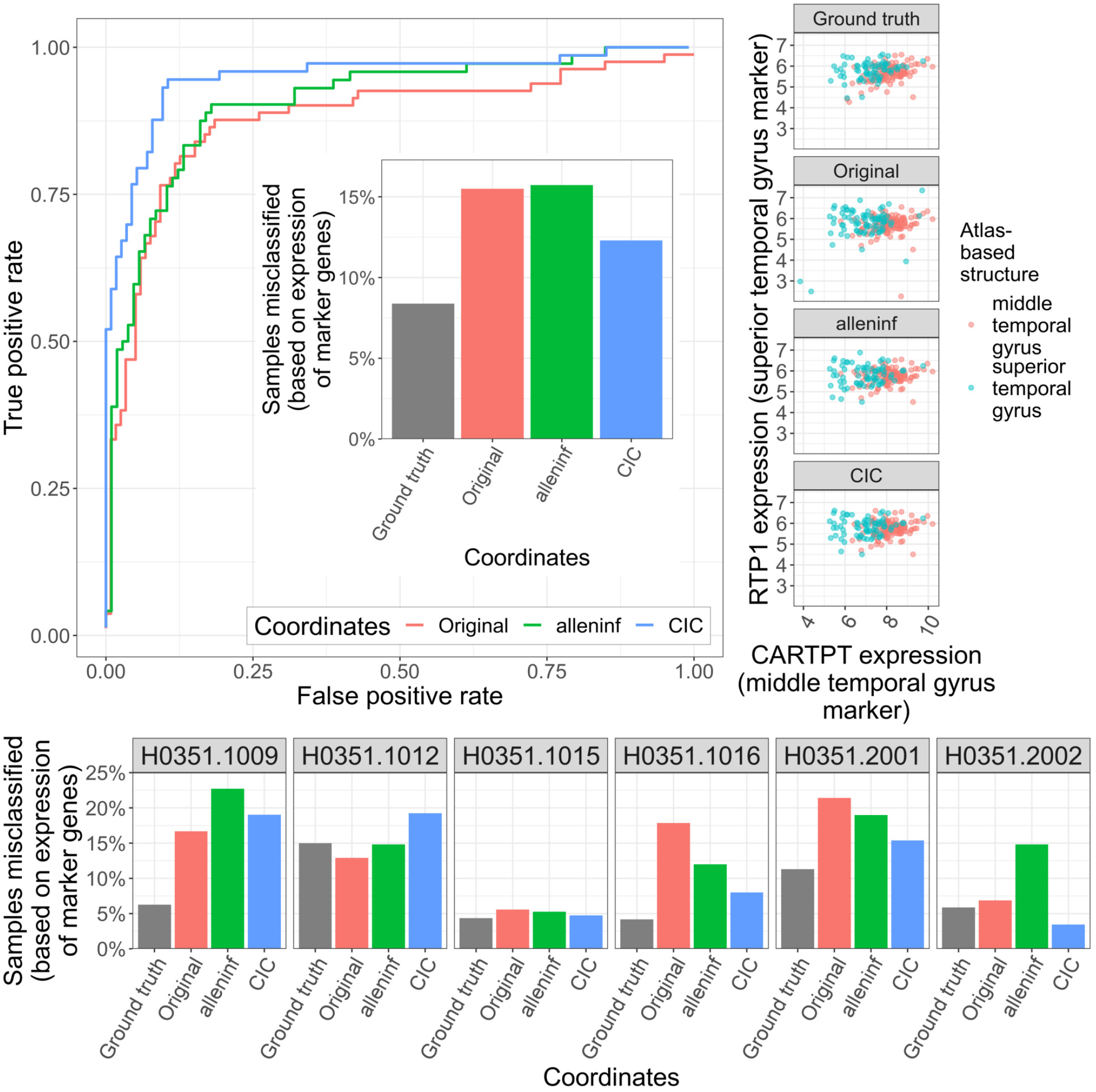
Expression-based separation analysis for superior temporal gyrus vs. middle temporal gyrus. Same as Figure S29, but comparing the superior temporal gyrus vs. middle temporal gyrus.

**Figure S32:**
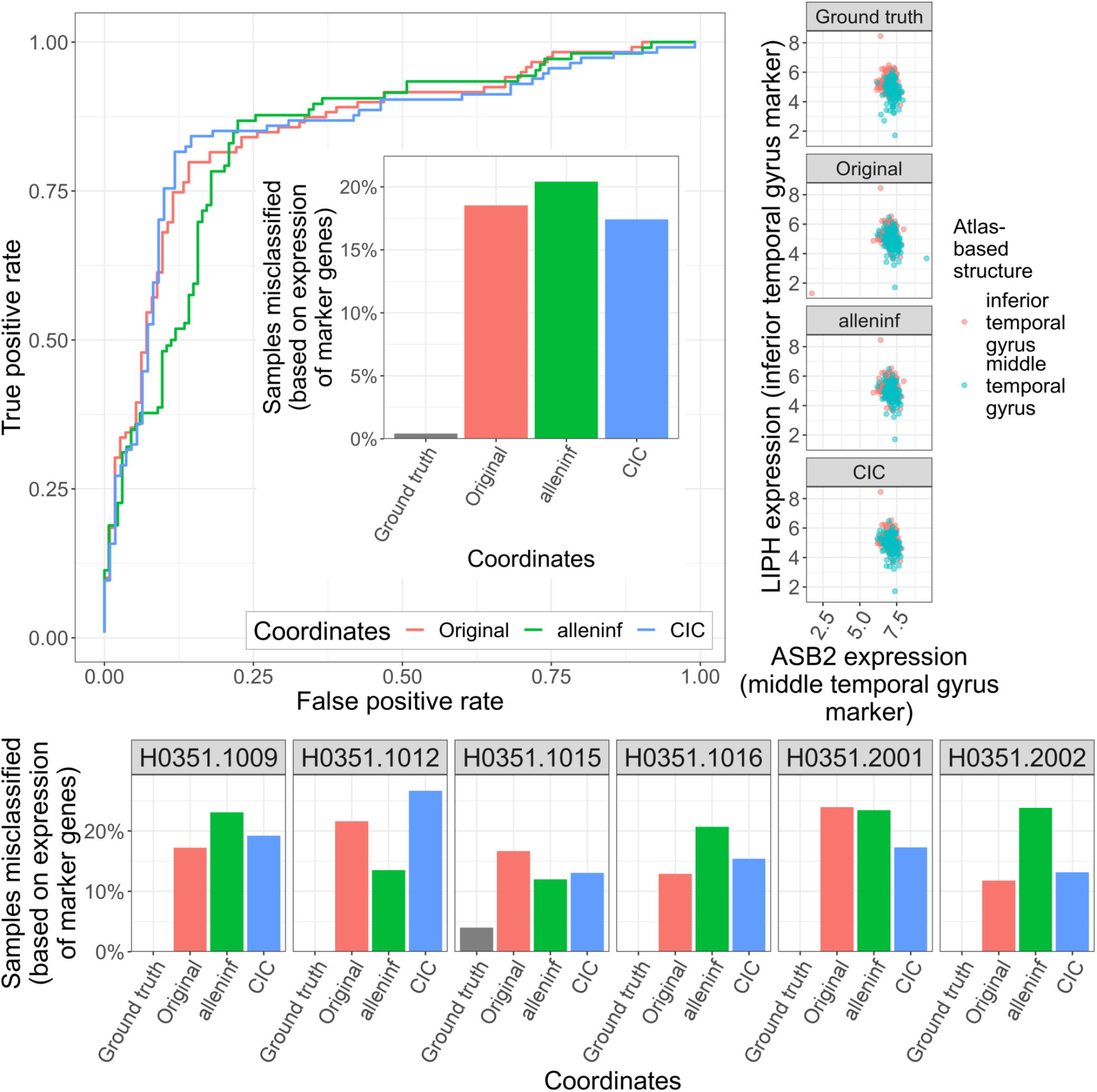
Expression-based separation analysis for middle temporal gyrus vs. inferior temporal gyrus. Same as Figure S29, but comparing the middle temporal gyrus vs. inferior temporal gyrus.

**Figure S33:**
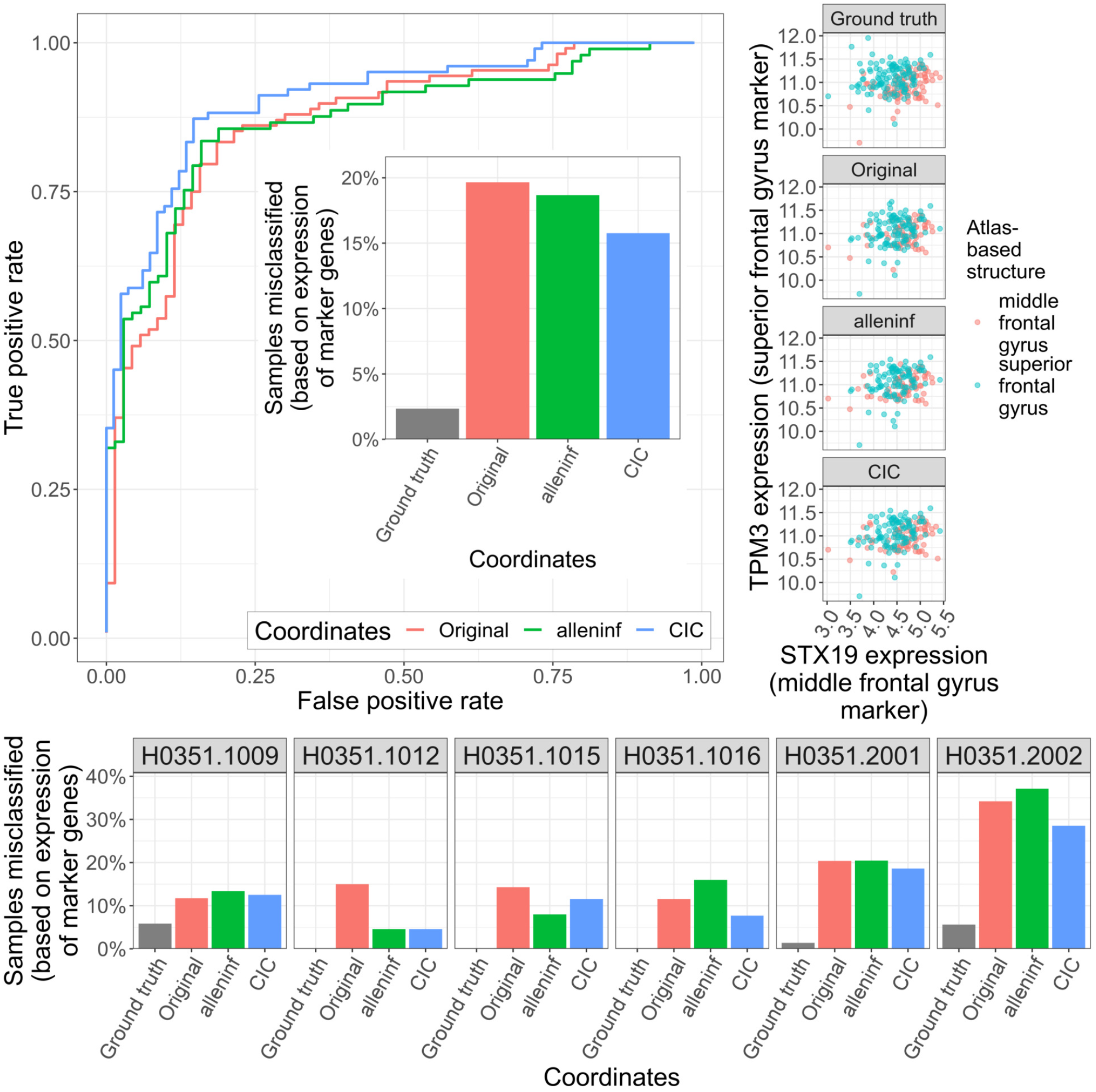
Expression-based separation analysis for superior frontal gyrus vs. middle frontal gyrus. Same as Figure S29, but comparing the superior frontal gyrus vs. middle frontal gyrus.

**Figure S34:**
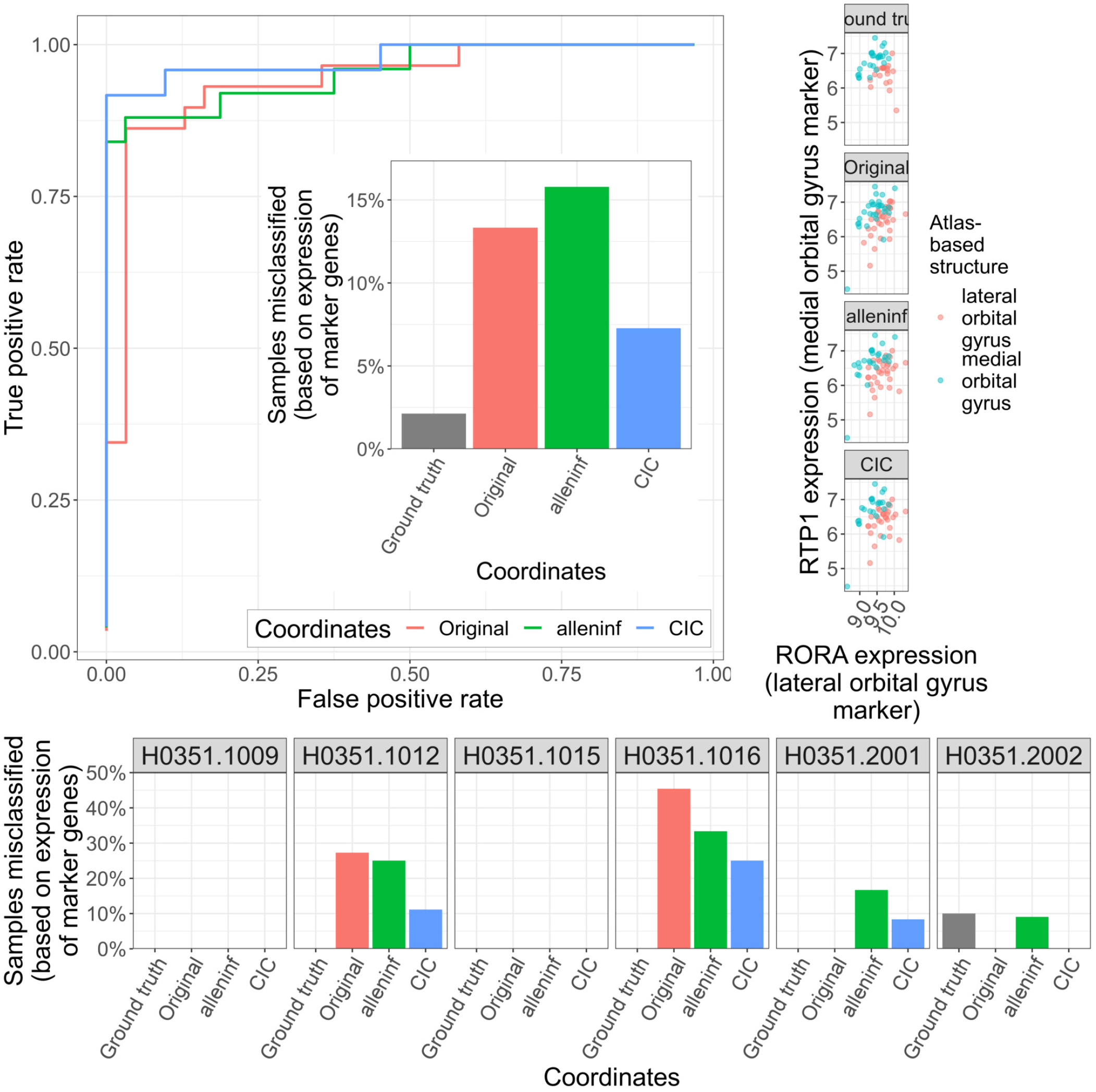
Expression-based separation analysis for lateral orbital gyrus vs. medial orbital gyrus. Same as Figure S29, but comparing the lateral orbital gyrus vs. medial orbital gyrus.

**Figure S35:**
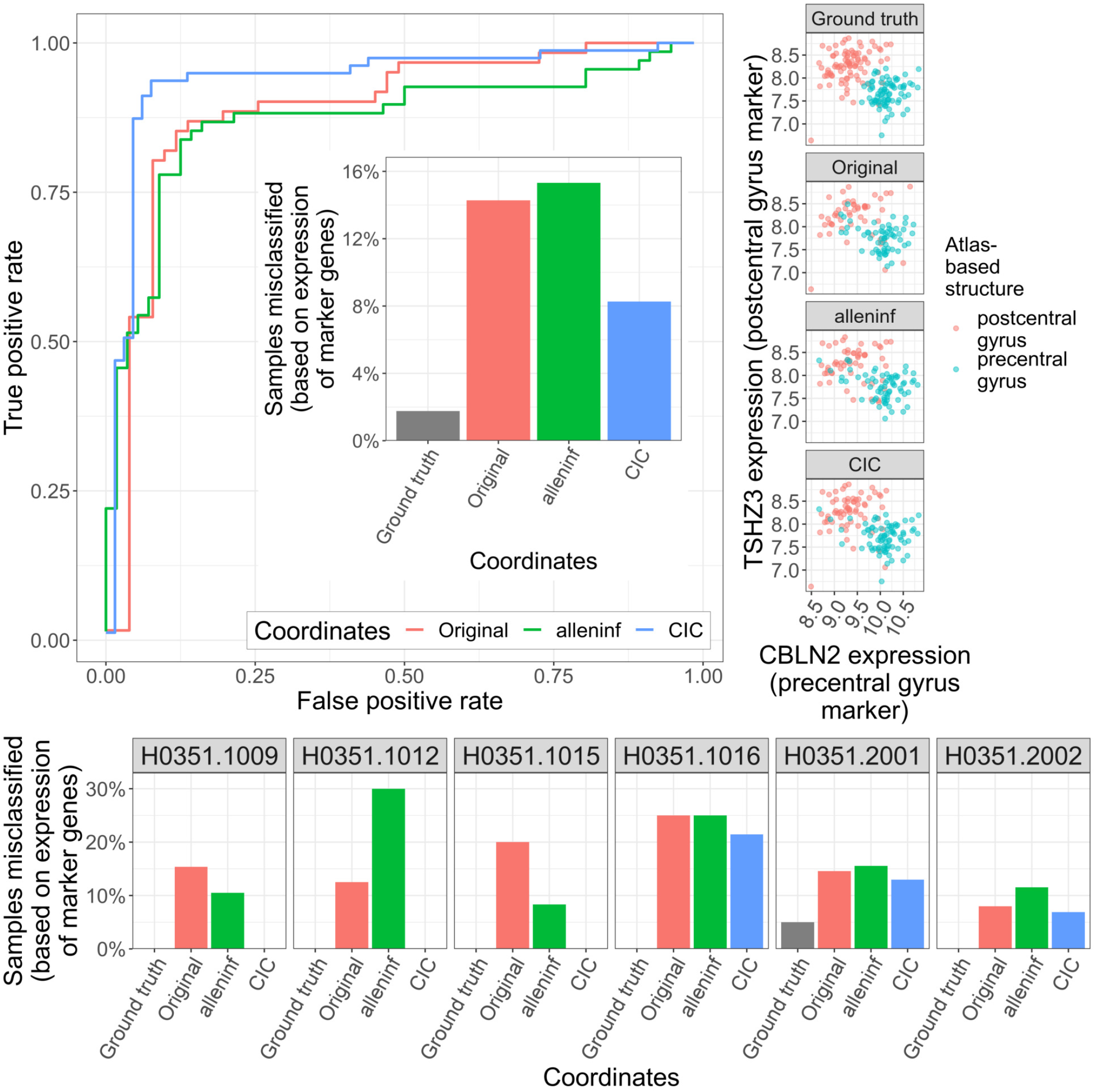
Expression-based separation analysis for precentral gyrus vs. postcentral gyrus, using grey matter-remapped coordinates. Same as Figure S29, but using coordinates-based locations after shifting grey matter samples which were positioned in white matter or outside brain tissue to the nearest grey matter voxel.

**Caption for Table S1. Median distances between new coordinates and previously-reported coordinates**. For each donor, the median distance between i) CIC and original coordinates and ii) CIC and alleninf coordinates are reported.

**Caption for Table S2**. *T*_1_**- and** *T*_2_**-weighted intensity-based measures of subject-template alignment accuracy**. For each subject, modality, and coordinate representation, provided are Spearman correlations summarizing the relationship between the intensities of the template image and template-aligned subject image. Image intensities are determined at the locations of all samples obtained from that subject using either original, alleninf, or CIC coordinates. Spearman correlations are computed and provided separately for both *T*_1_- and *T*_2_-weighted image sets.

**Caption for Table S3. Misclassification in coordinate-based placement of samples into coarse anatomical structures**. Numbers and proportions of samples (in)correctly classified in the coarse anatomical structure that they are dissected from, based on the placement of their coordinates within the AHRA.

**Caption for Table S4. Counts of samples’ true coarse structures and coordinate-assigned coarse structures**. For each true coarse (AHBA) structure that samples are dissected from, the number of samples assigned to each AHRA coarse structure based on each of the three coordinate representations are listed.

**Caption for Table S5. False negative rate (FNR) for coarse structure classification**. For each true coarse (AHBA) structure that samples are dissected from, the false negative rate in coordinate-based structure assignments (associated with each of the three coordinate representations) are listed.

**Caption for Table S6. Misclassification in coordinate-based placement of samples into cortical lobe structures**. Numbers and proportions of samples (in)correctly classified in the cortical lobes that they are dissected from, based on the placement of their coordinates within the AHRA.

**Caption for Table S7. Counts of samples’ true cortical lobes and coordinate-assigned cortical lobes**. For each true cortical lobe (AHBA) structure that samples are dissected from, the number of samples assigned to each AHRA cortical lobe based on each of the three coordinate representations are listed.

**Caption for Table S8. False negative rate (FNR) for cortical lobe classification**. For each true cortical lobe (AHBA) structure that samples are dissected from, the false negative rate in coordinate-based structure assignments (associated with each of the three coordinate representations) are listed.

**Caption for Table S9. Effects of remapping grey matter tissue sample coordinates to the nearest grey matter voxel**. The number of samples gained by each AHRA structure after remapping grey matter sample coordinates to the nearest grey matter voxel are provided.

**Caption for Table S10. Summary statistics for distance shifts due to remapping grey matter tissue sample coordinates to the nearest grey matter voxel**. Mean, median, and maximum distances that coordinates are shifted after remapping grey matter sample coordinates to the nearest grey matter voxel are provided for each of the three coordinate representations, along with the numbers and proportions of samples that shift above 2, 5, and 8 mm.

**Caption for Table S11**. *T*_1_**- and** *T*_2_**-weighted intensity-based measures of subject-template alignment accuracy, after remapping grey matter sample coordinates to the closest grey matter voxel**. For each subject, modality, and coordinate representation, provided are Spearman correlations summarizing the relationship between the intensities of the template image and template-aligned subject image. Image intensities are determined at the locations of all samples obtained from that subject using either original, alleninf, or CIC coordinates after remapping grey matter sample coordinates to the closest grey matter voxel. Spearman correlations are computed and provided separately for both *T*_1_- and *T*_2_-weighted image sets.

**Caption for Table S12. Misclassification in coordinate-based placement of samples into coarse anatomical structures, after remapping grey matter sample coordinates to the closest grey matter voxel**. Numbers and proportions of samples (in)correctly classified in the coarse anatomical structure that they are dissected from, based on the placement of their coordinates within the AHRA after remapping grey matter sample coordinates to the closest grey matter voxel.

**Caption for Table S13. AUC as a function of** *k*. The area under the ROC curve (AUC) was constructed from comparing an AHRA-based mask of core nuclei with a predicted mask built from thresholding a coordinate-based *k*-nearest neighbour interpolation of gene expression data within the thalamus. Here, AUC values are provided as a function of *k*.

**Caption for Table S14. AUC in the expression interpolation analysis**. AUC values are provided for *k* = 5.

**Caption for Table S15. AUC in the expression interpolation analysis, using grey matter-remapped coordinates**. AUC values are provided for *k* = 5, using grey matter-remapped coordinates.

**Caption for Table S16. AUC as a function of** *k*, **using grey matter-remapped coordinates**. The area under the ROC curve (AUC) was constructed from comparing an AHRA-based mask of core nuclei with a predicted mask built from thresholding a coordinate-based *k*-nearest neighbour interpolation of gene expression data within the thalamus, after remapping grey matter sample coordinates to the closest grey matter voxel. Here, AUC values are provided as a function of *k*.

**Caption for Table S17. Summary statistics for interpolation analysis using alternative processing decisions**. Mean, median, interquartile range, minimum, and maximum are reported for each structure and coordinate representation.

**Caption for Data S1. CIC coordinates**. A table (3702 rows corresponding to tissue samples) listing CIC coordinates alongside previous (original and alleninf) coordinates, along with sample metadata. Coordinates are provided in relation to both templates (MNI ICBM152 nonlinear symmetric 2009c and MNI152 6th generation nonlinear asymmetric) analyzed here.

**Caption for Data S2. Representative sample of imaging-transcriptomics literature**. A table (63 rows corresponding to studies) demonstrating coordinate use patterns.

**Caption for Data S3. Coordinate-based structure assignments**. A table (3702 rows corresponding to tissue samples) listing the AHRA structures that samples are placed in based on MNI coordinates, along with sample metadata. Structure assignments are provided in relation to MNI coordinates and the AHRA atlas mapped to both templates (MNI ICBM152 nonlinear symmetric 2009c and MNI152 6th generation nonlinear asymmetric) analyzed here.

**Caption for Data S4. Structure mappings relating the AHBA and AHRA at a coarse level**. A table (4336 rows corresponding to AHRA-defined structures) that relate each AHRA structure to a coarse structure that is also identifiable within the AHBA.

**Caption for Data S5. True (AHBA) and coordinate-based (AHRA) assignment of samples to coarse structures**. A table (11106 rows corresponding to tissue samples × 3 coordinates) associating each tissue sample and coordinate choice to the AHRA structure that it falls within.

**Caption for Data S6. Structure mappings relating the AHBA and AHRA at the level of cortical lobes**. A table (82 rows corresponding to AHRA-defined cortical structures) that relate each AHRA cortical structure to a lobe that is also identifiable within the AHBA.

**Caption for Data S7. True (AHBA) and coordinate-based (AHRA) assignment of cortical samples to cortical lobes**. A table (5850 rows corresponding to cortical tissue samples × 3 coordinates) associating each cortical tissue sample and coordinate choice to the AHRA structure that it falls within.

**Caption for Data S8. CIC coordinates after remapping grey matter sample coordinates to the closest grey matter voxel**. A table (3702 rows corresponding to tissue samples) listing grey matter-remapped CIC coordinates alongside previous (original and alleninf) coordinates, along with sample metadata. Coordinates are provided in relation to the MNI ICBM152 nonlinear symmetric 2009c template.

**Caption for Data S9. Coordinate-based structure assignments after remapping grey matter sample coordinates to the closest grey matter voxel**. A table (3702 rows corresponding to tissue samples) listing the AHRA structures that samples are placed in based on grey matter-remapped MNI coordinates, along with sample metadata. Structure assignments are provided in relation to the MNI ICBM152 nonlinear symmetric 2009c template.

**Caption for Data S10. Similarity between each samples’ gene expression profile and the mean gene expression profile of all other samples in the same coordinate-assigned AHRA region**. A table (22212 rows corresponding to tissue samples × 3 coordinates × 2 templates (sym09c and nlin6asym) that lists the similarity (as the Pearson correlation *r)* between the selected tissue sample’s gene expression profile and the average gene expression profile of all other tissue samples assigned to the same AHRA region based on their coordinates.

**Caption for Data S11. Tissue samples used for the expression similarity analysis**. A table (2254 rows from Data S10) representing the subset of tissue samples used in the expression similarity analysis. These are samples placed in different regions when comparing original and CIC, and alleninf and CIC coordinates.

**Caption for Data S12. Similarity between each samples’ gene expression profile and the mean gene expression profile of all other samples in the same coordinate-assigned AHRA region, after remapping grey matter sample coordinates to the closest grey matter voxel**. A table (11106 rows corresponding to tissue samples × 3 coordinates that lists the similarity (as the Pearson correlation *r)* between the selected tissue sample’s gene expression profile and the average gene expression profile of all other tissue samples assigned to the same AHRA region. Structure assignments are based on grey matter-remapped coordinates in relation to the MNI ICBM152 nonlinear symmetric 2009c template.

**Caption for Data S13. Tissue samples used for the expression similarity analysis, after remapping grey matter sample coordinates to the closest grey matter voxel**. A table (2254 rows from Data S12) representing the subset of grey matter-remapped tissue samples used in the expression similarity analysis. These are samples placed in different regions when comparing original and CIC, and alleninf and CIC coordinates, after remapping grey matter sample coordinates to the closest grey matter voxel.

**Caption for Data S14. Structure mappings relating the AHBA and AHRA thalamus subdivisions**. A table (211 rows corresponding to AHRA-defined thalamus substructures) that relate each AHRA thalamus subdivision to a subdivision identifiable within the AHBA.

**Caption for Data S15. Thalamic tissue samples**. A table (350 rows corresponding to 175 tissue samples dissected from the thalamus along with their positions in the contralateral hemisphere) describing the samples and their coordinates used in the interpolation analysis.

**Caption for Data S16. NeuroQuery terms**. A table (79 rows corresponding to an *a priori* search of diverse terms) and associated matches.

**Caption for Data S17. AHBA probes**. A table (17205 rows corresponding to AHBA probes) describing the probe IDs, names, and associated genes used in expression analyses.

