## Supplementary figures and images for "Accurate spatial localization of Allen Human Brain Atlas gene expression data for human neuroimaging"

### bias_gene_ranks.png

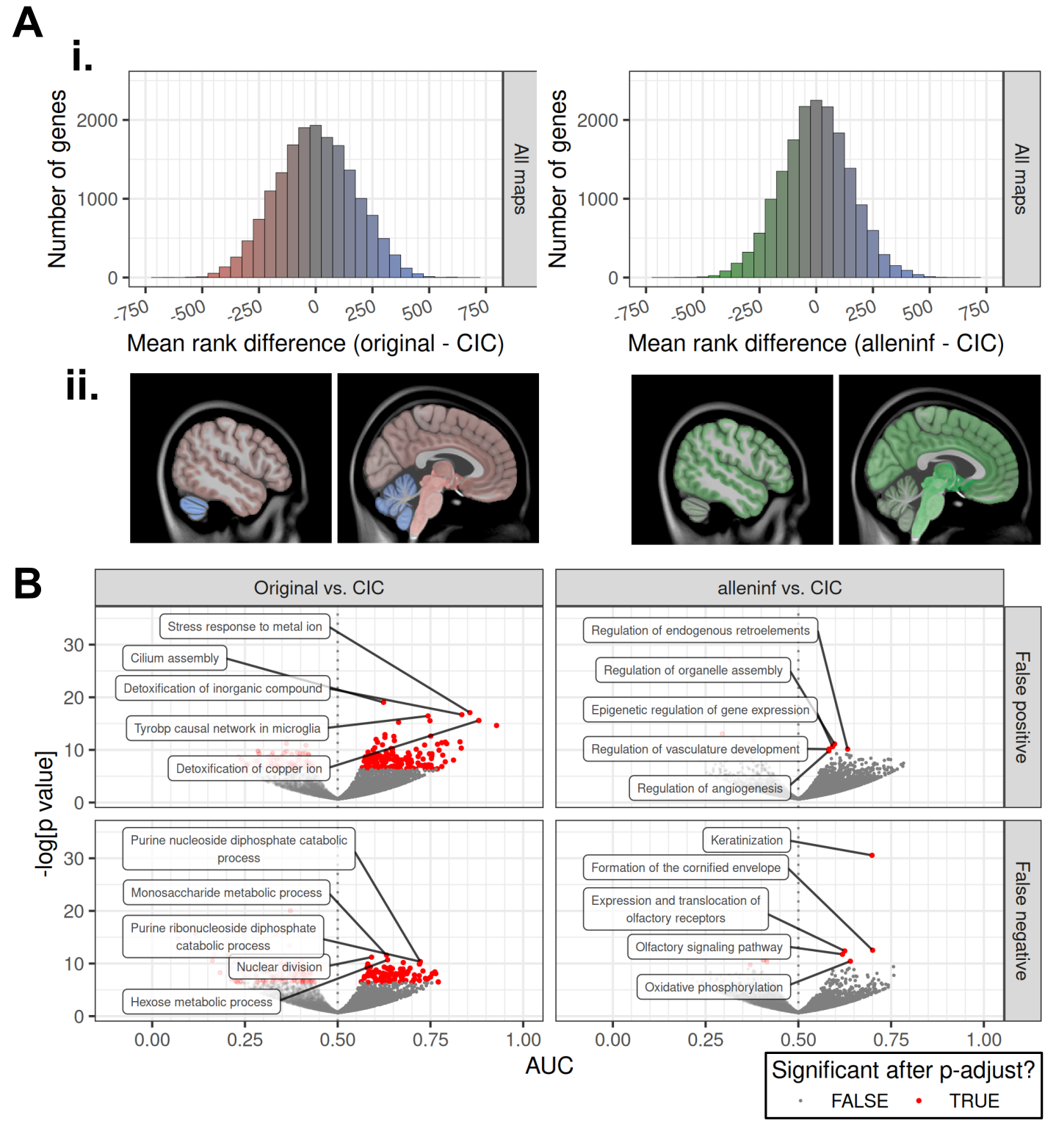

### bias_neuroquery_panel.png

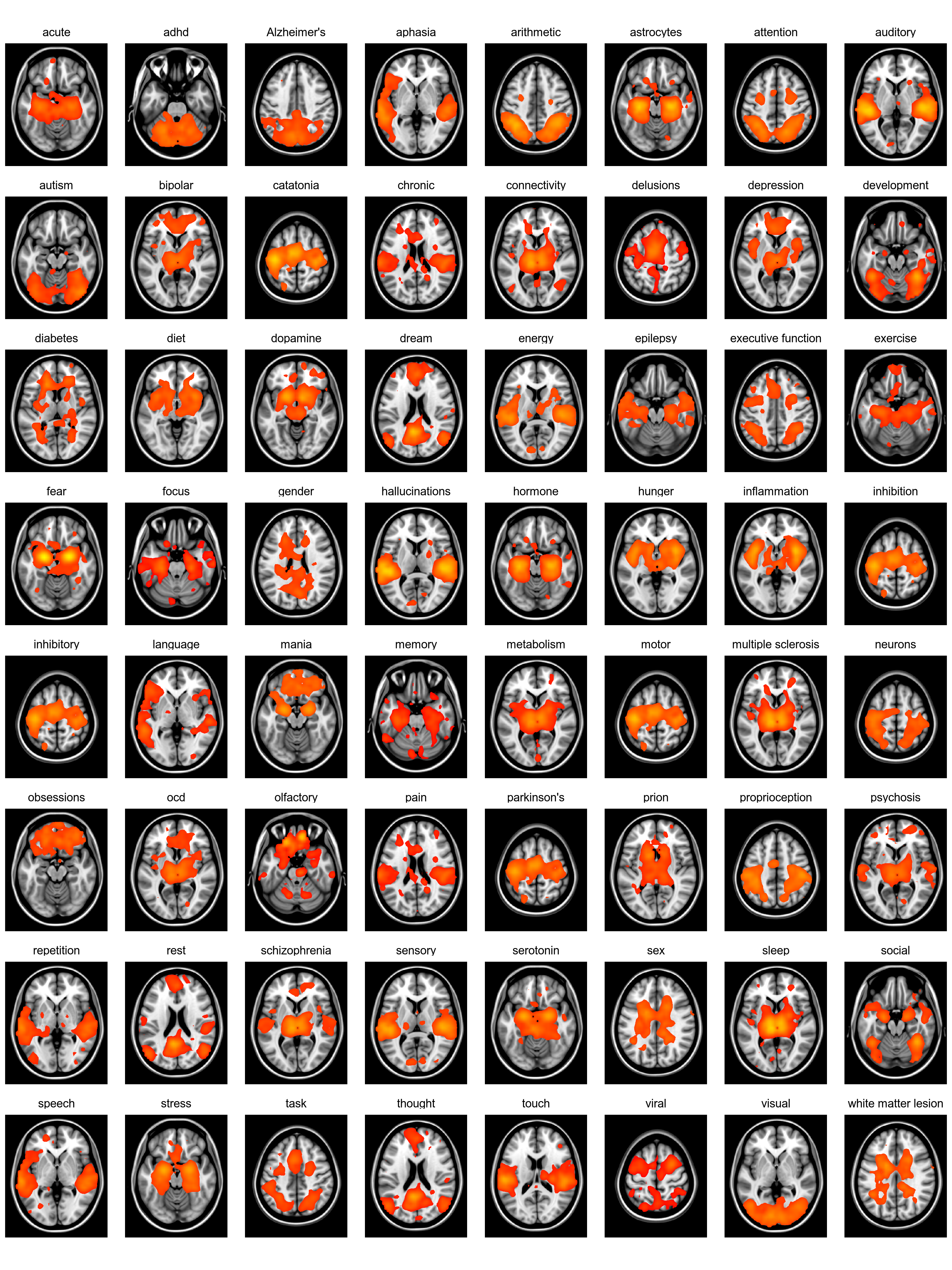

### bias_validate_modules.png

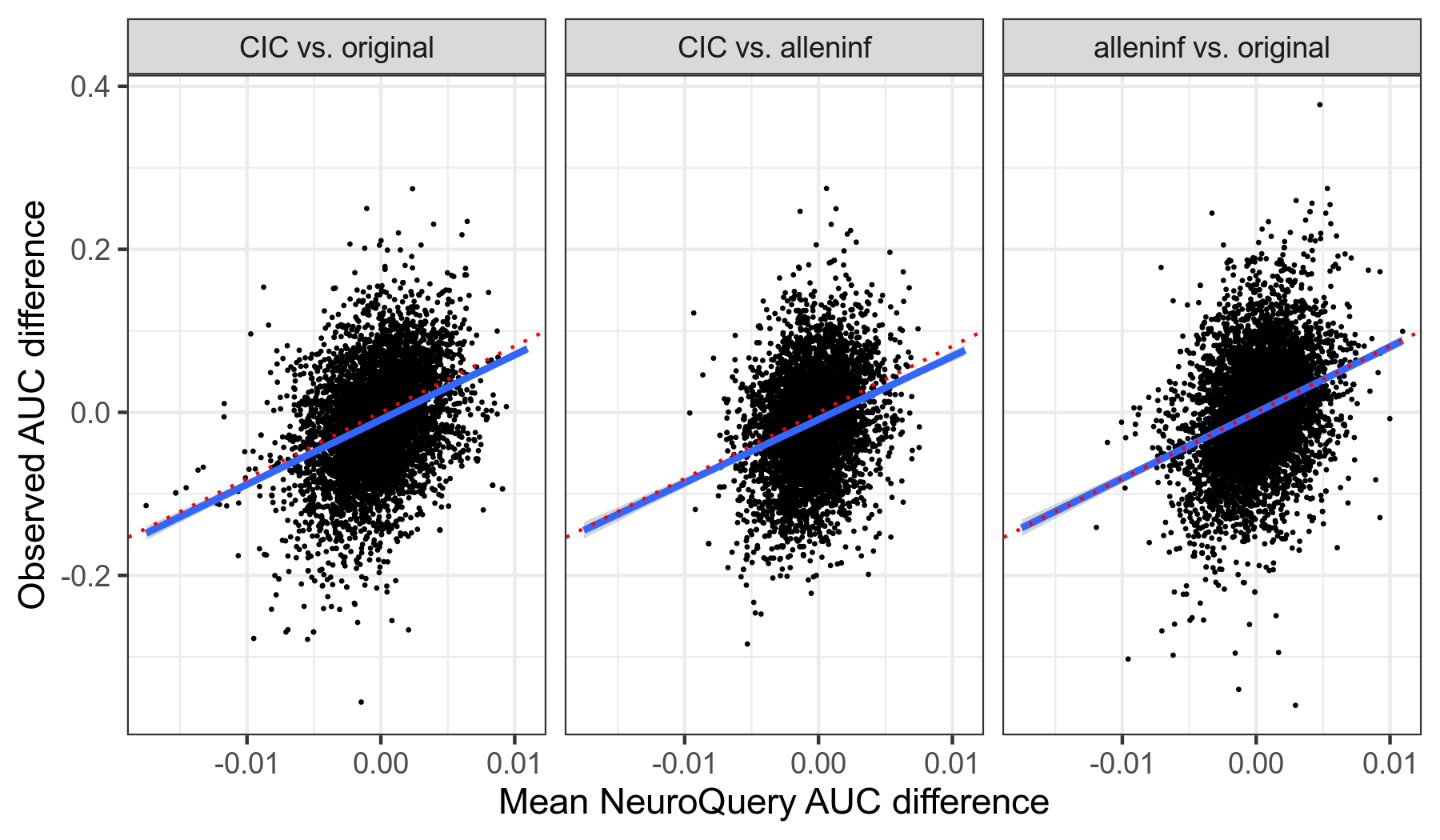

### comparison_distance_shifts.png

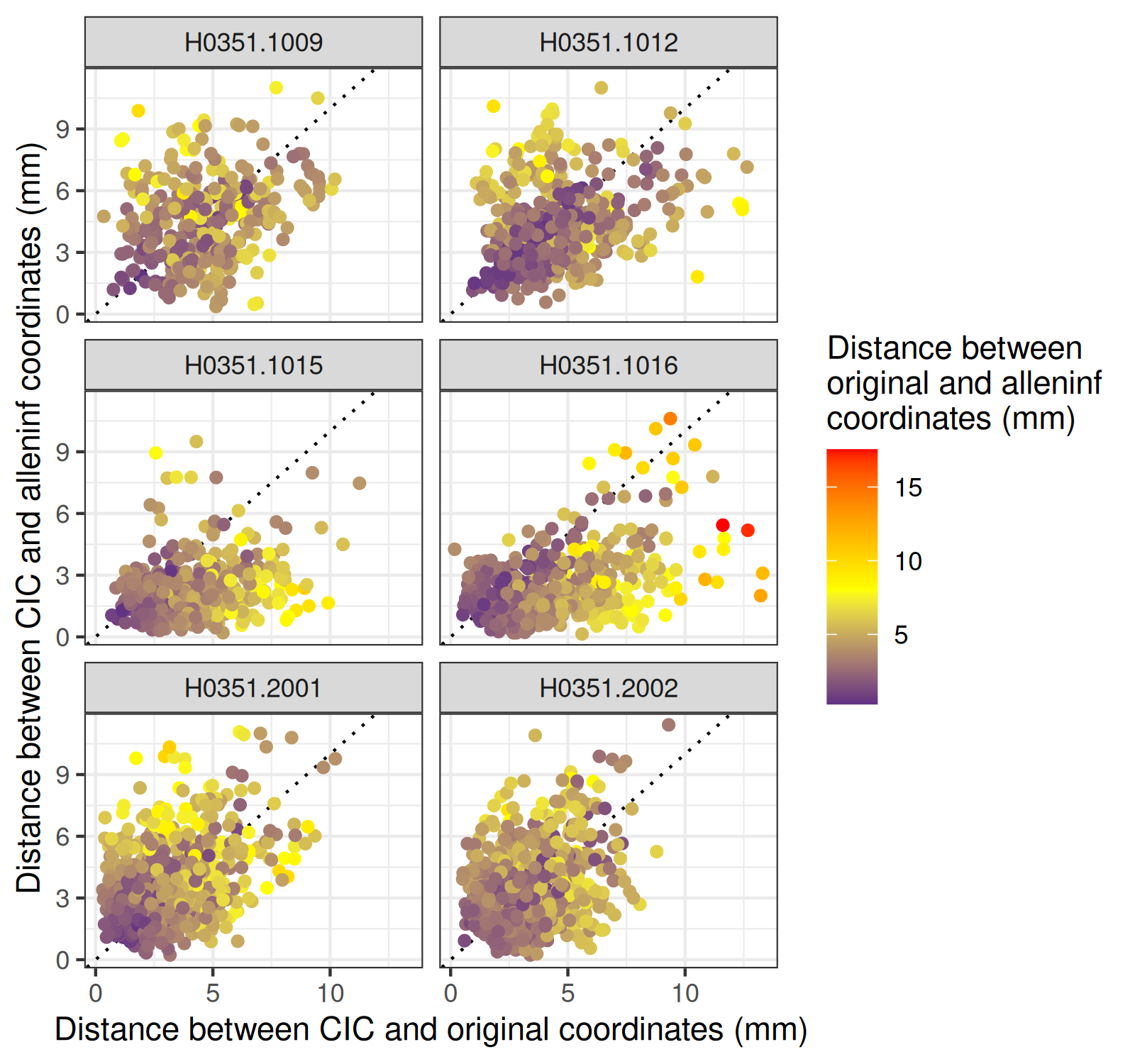

### expression_interpolation_auc_over_k.png

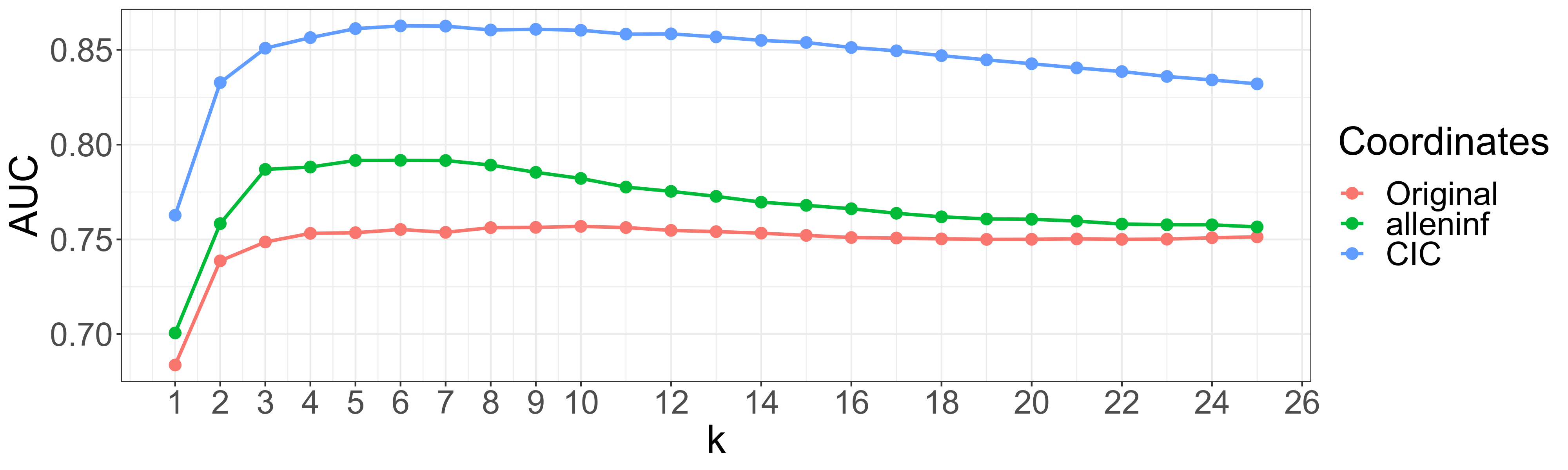

### expression_interpolation_auc_over_k_remapped.png

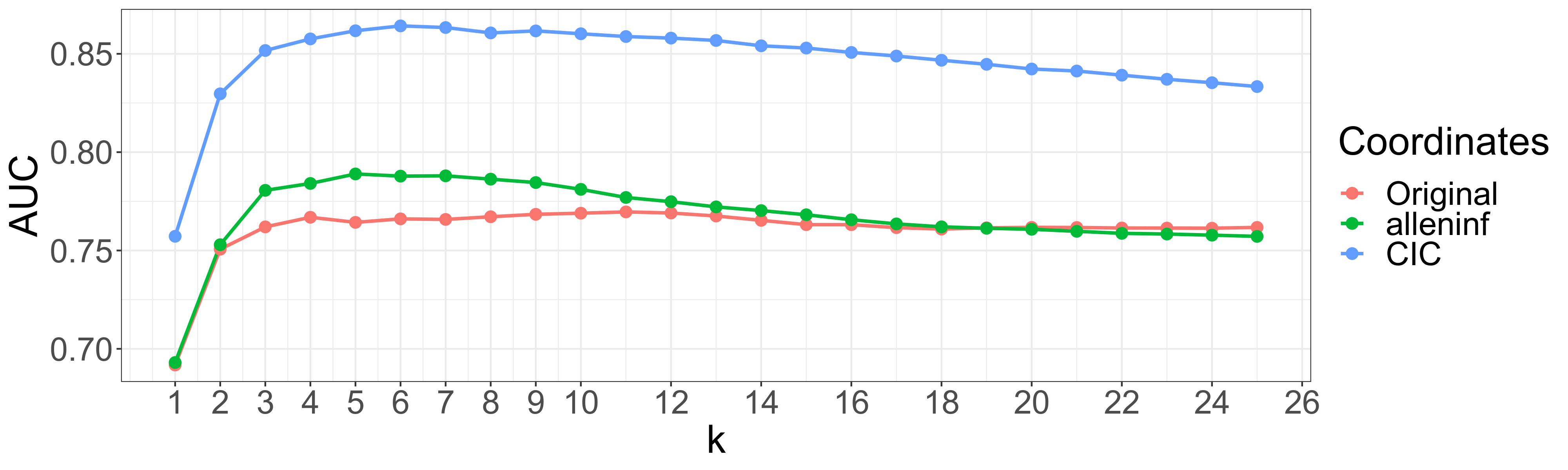

### expression_interpolation_panel_remapped.png

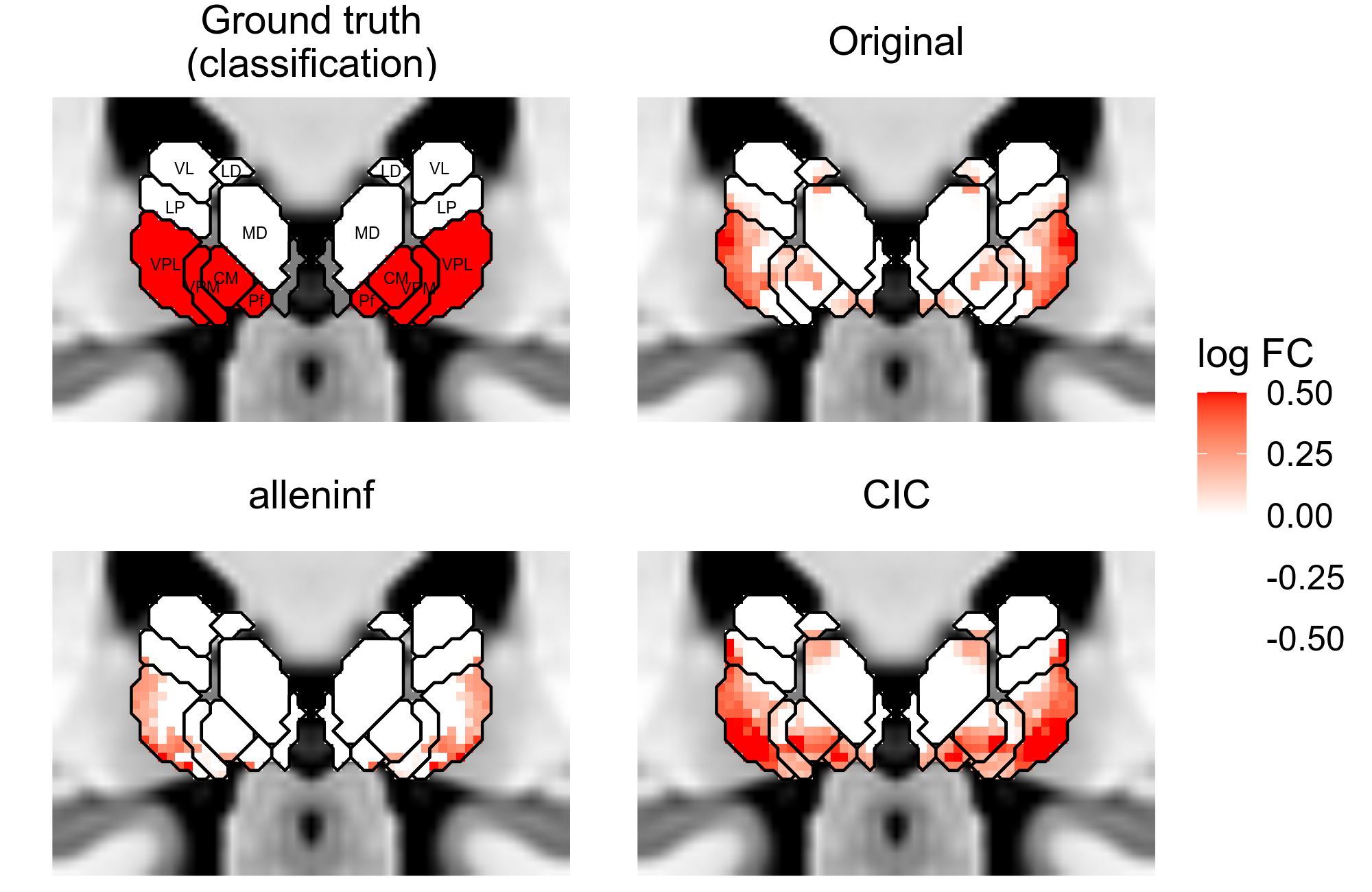

### expression_separation_lasso_training.png

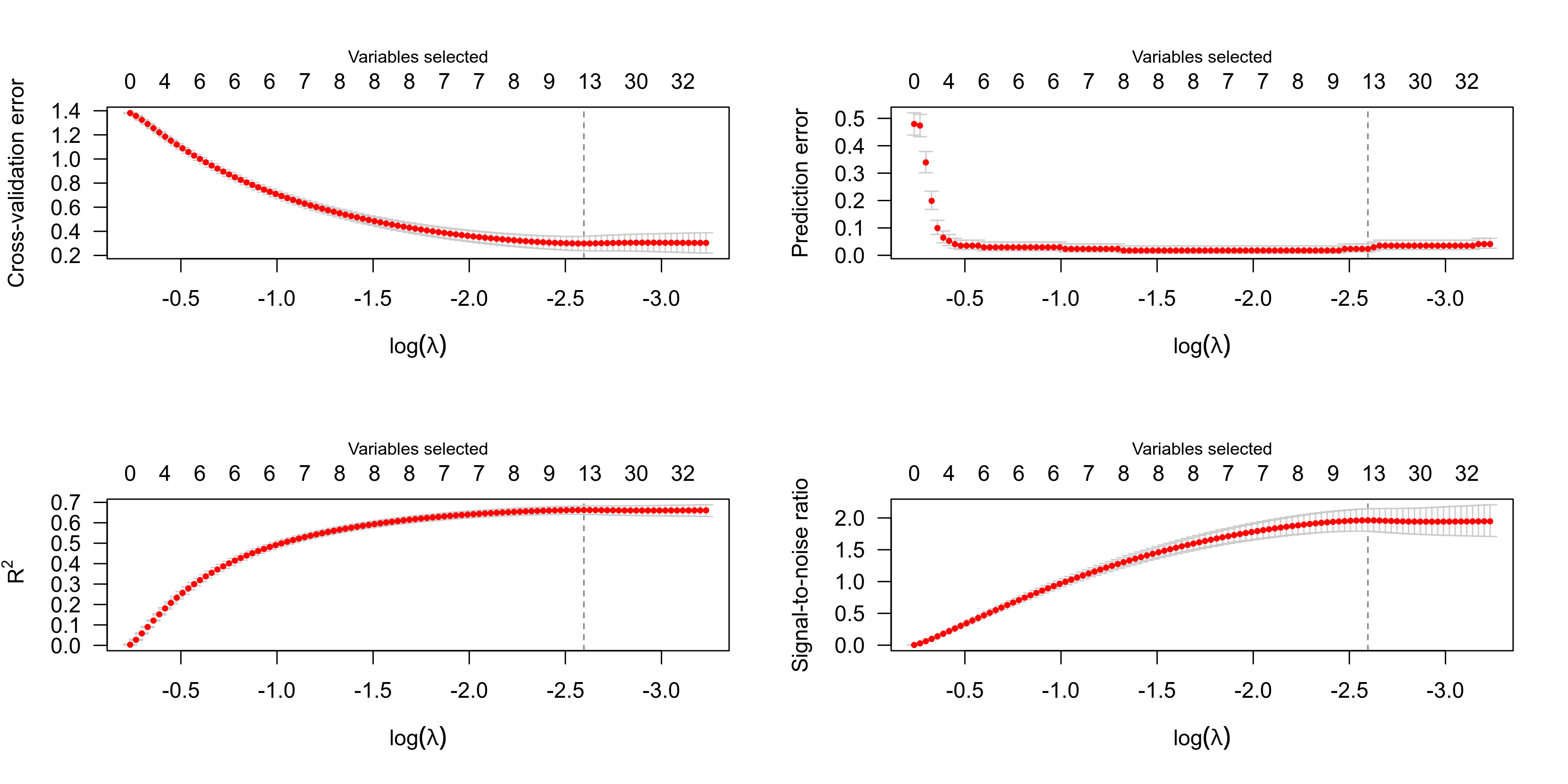

### expression_separation_medial_lateral_orbital_gyrus.png

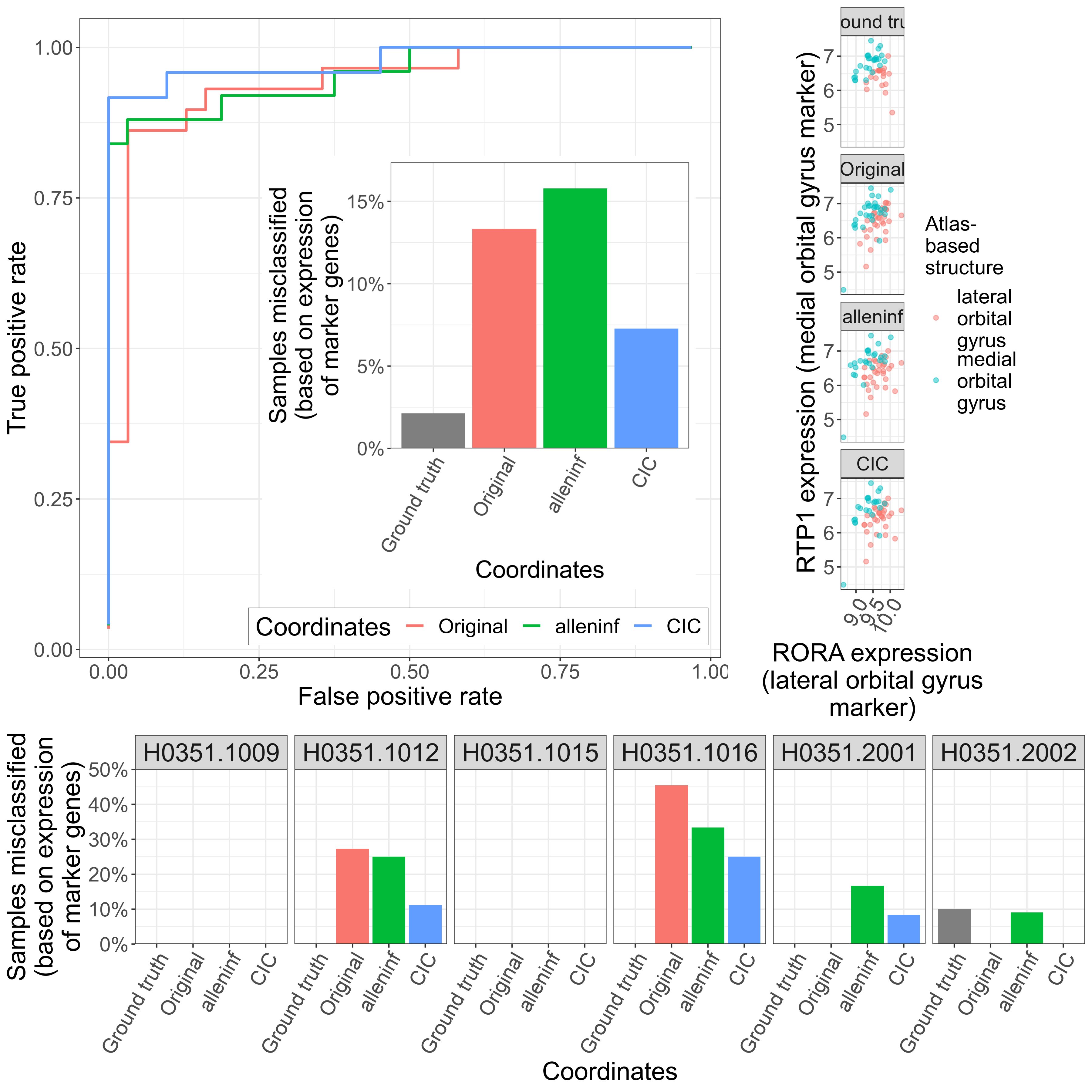

### expression_separation_middle_inferior_temporal_gyrus.png

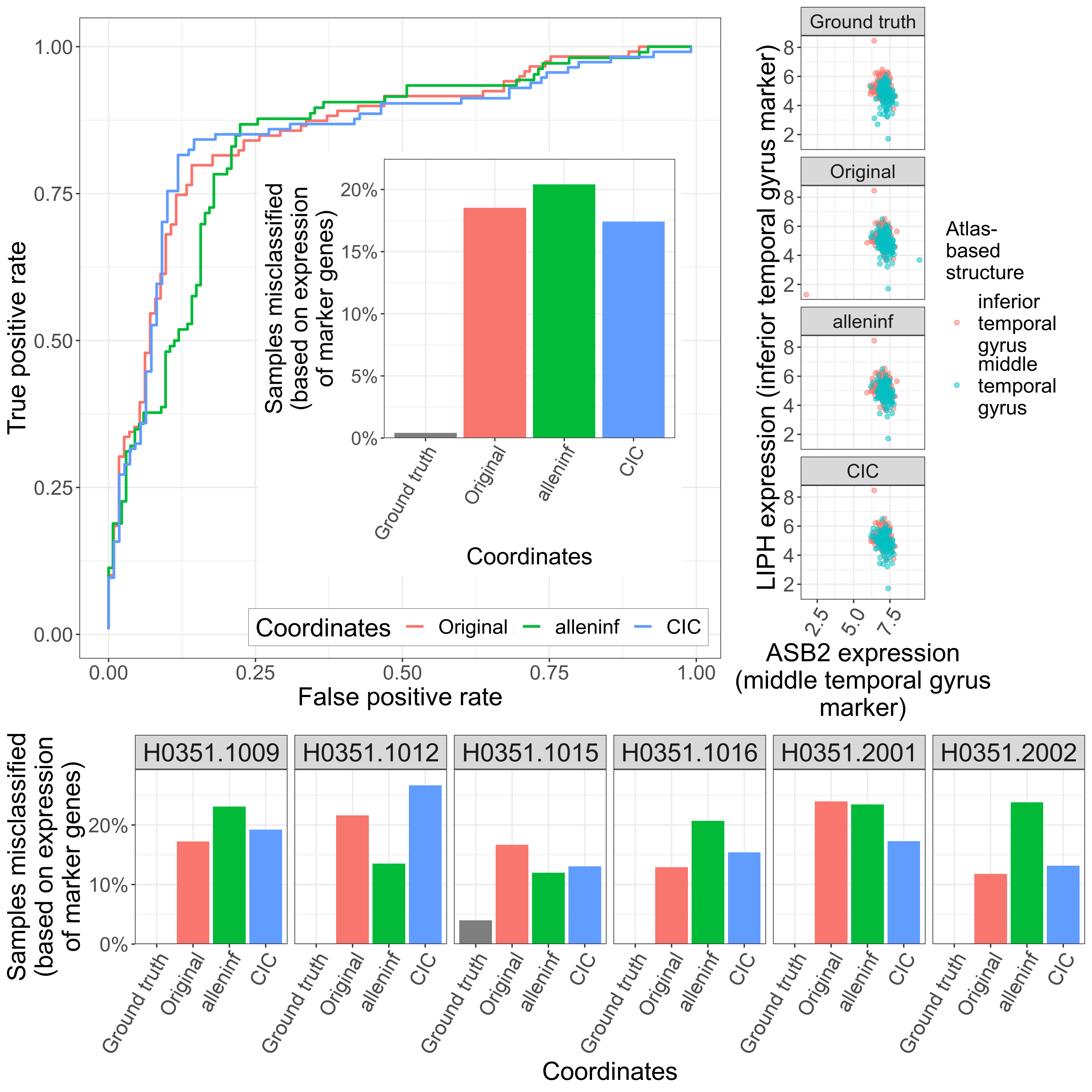

### expression_separation_precentral_postcentral_gyrus.png

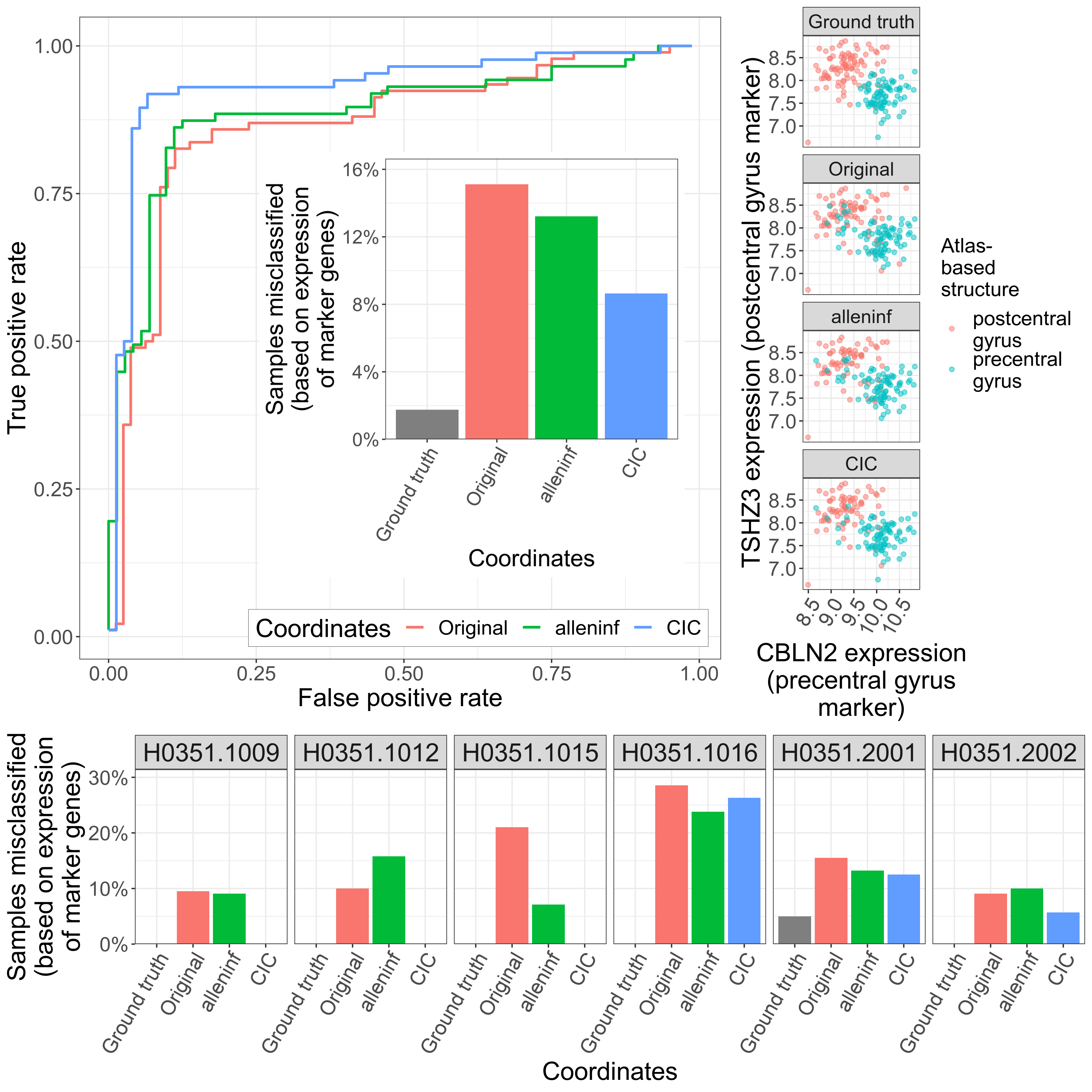

### expression_separation_precentral_postcentral_gyrus_remapped.png

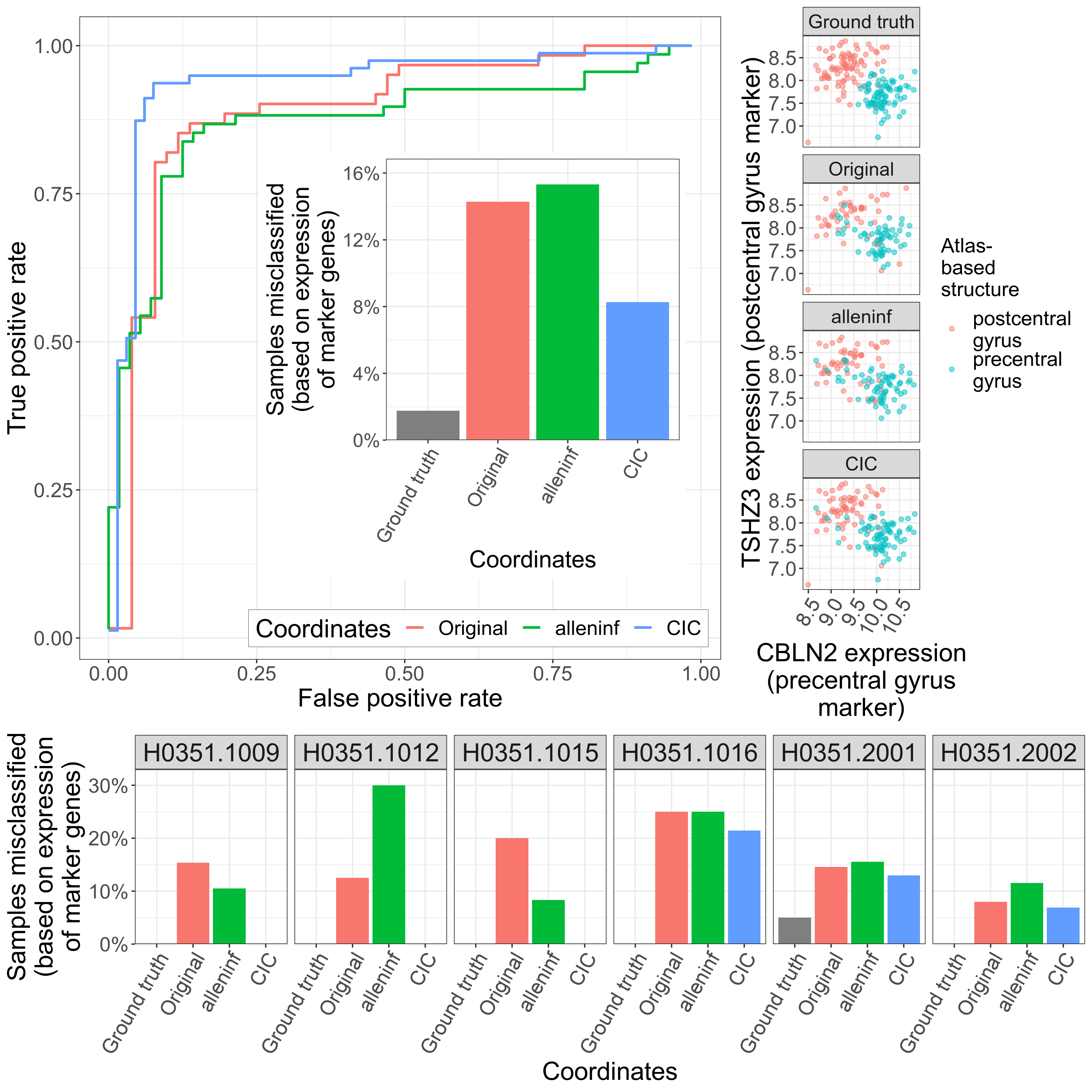

### expression_separation_superior_inferior_temporal_gyrus.png

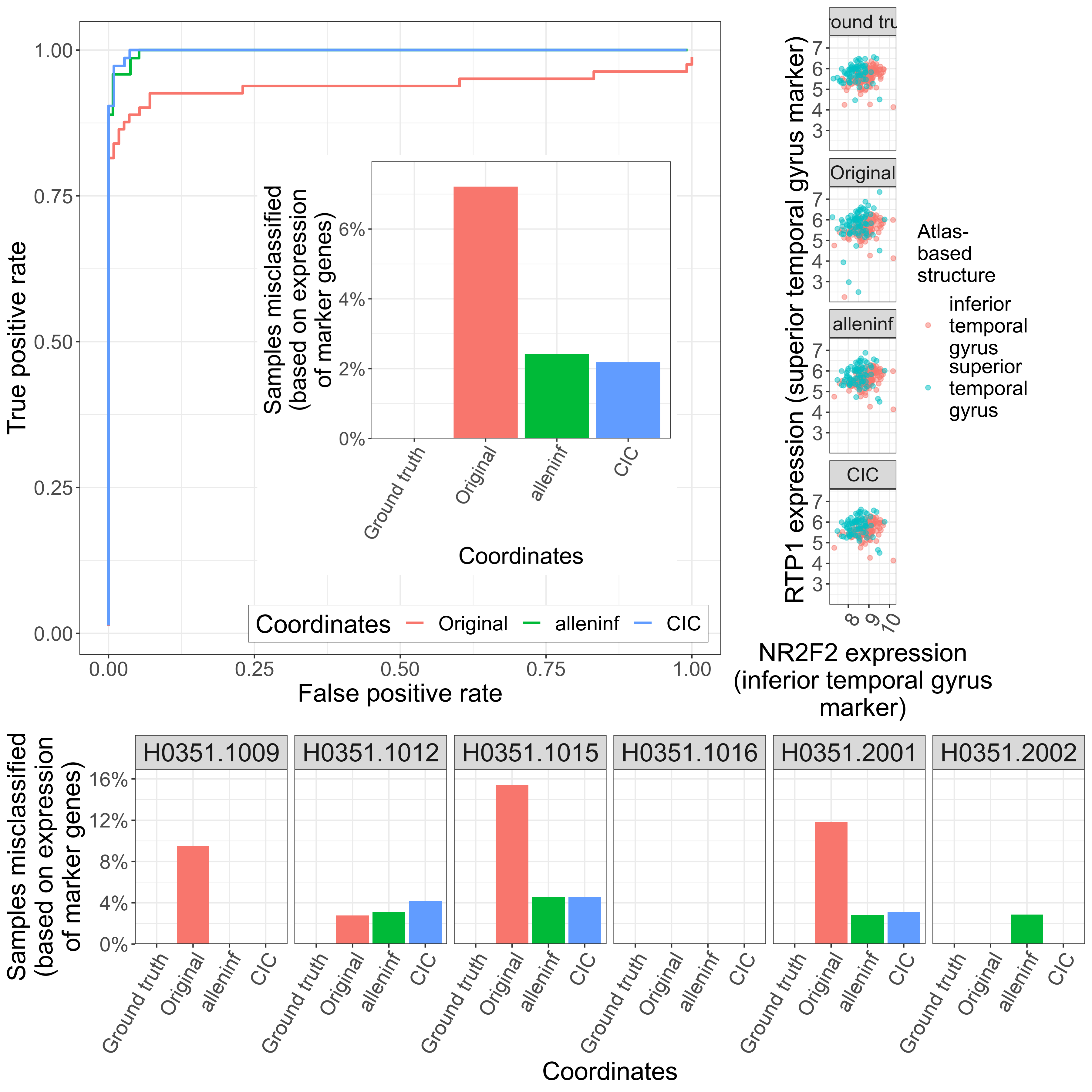

### expression_separation_superior_middle_frontal_gyrus.png

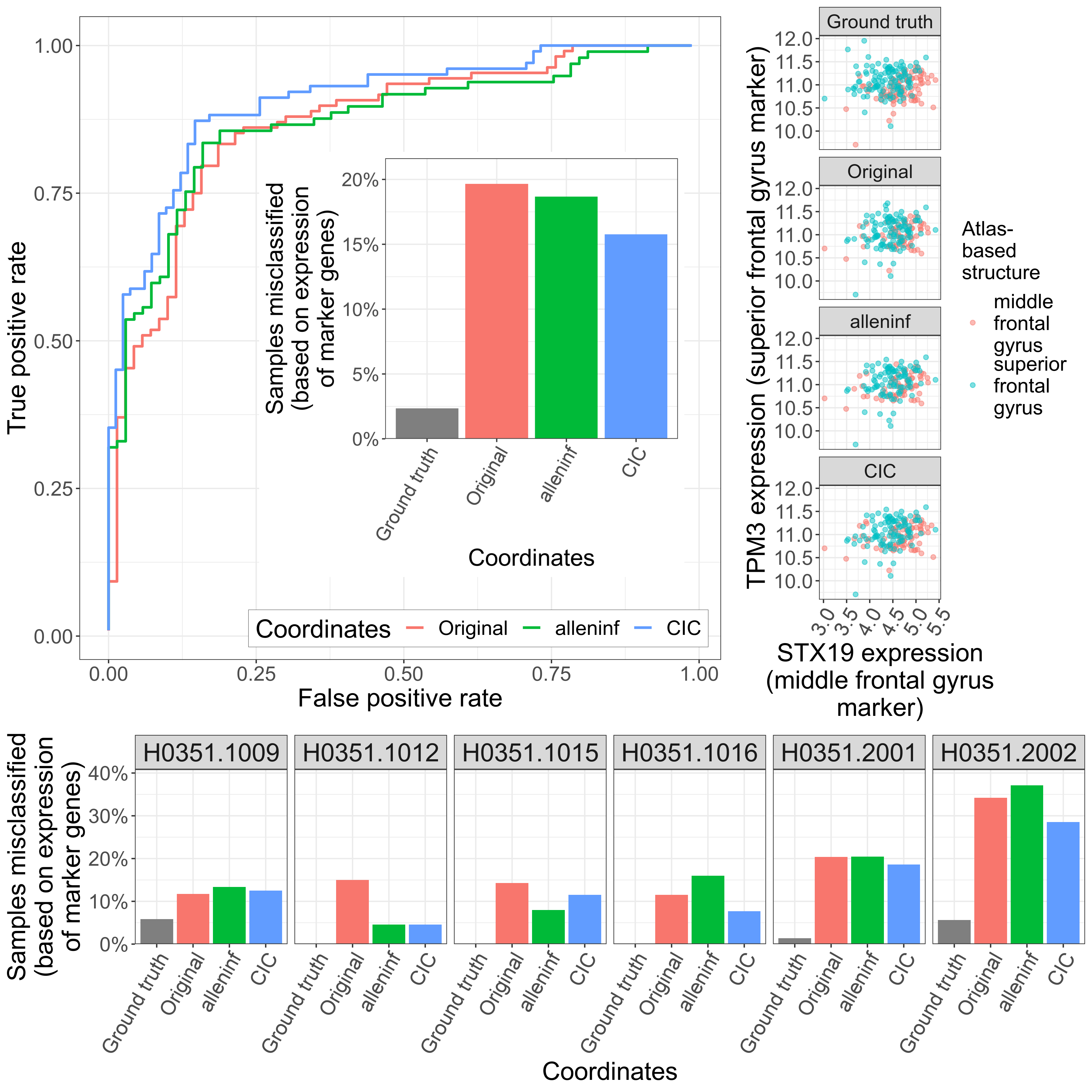

### expression_separation_superior_middle_temporal_gyrus.png

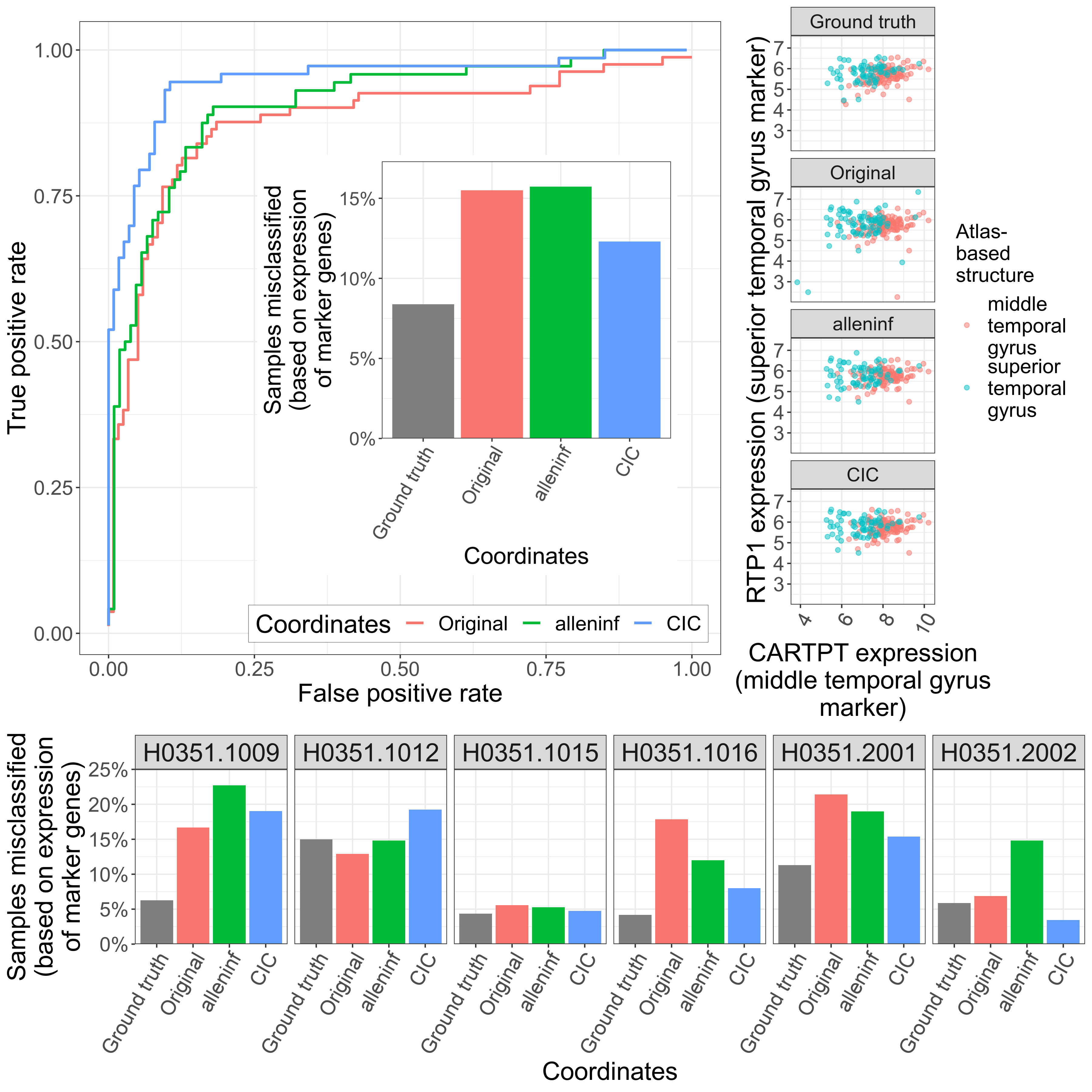

### expression_similarity_subjectwise.png

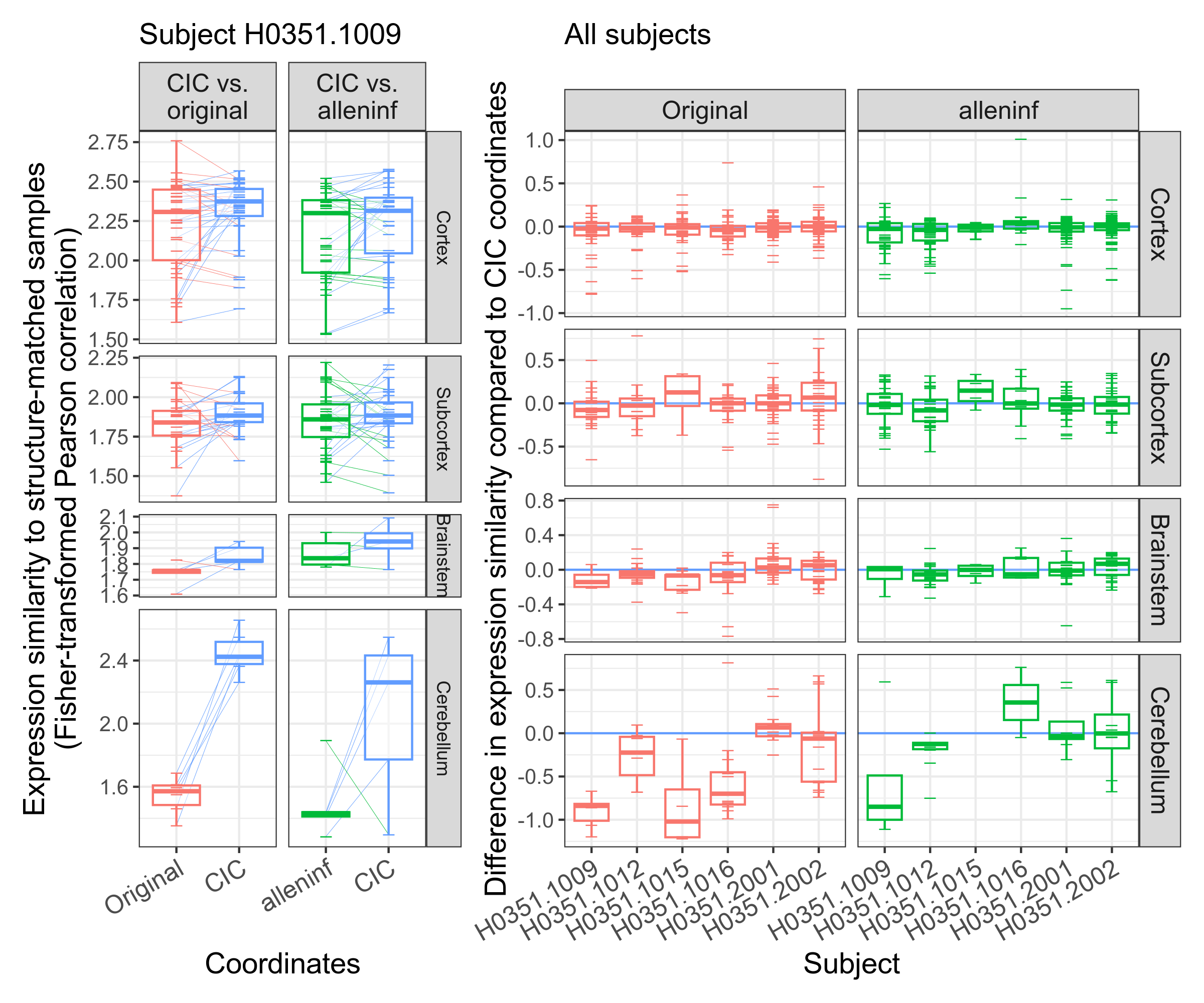

### expression_similarity_subjectwise_remapped.png

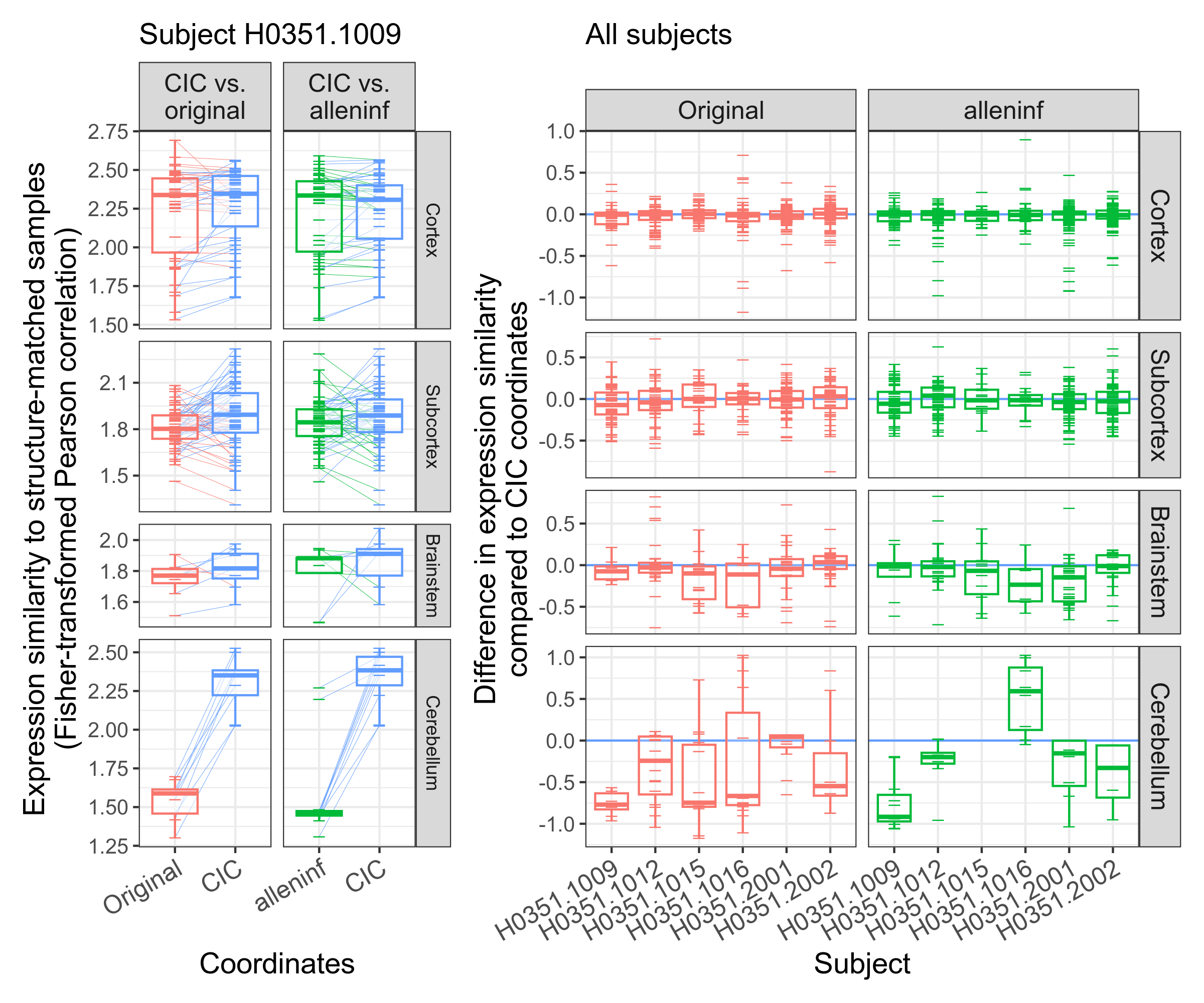

### fig1.png

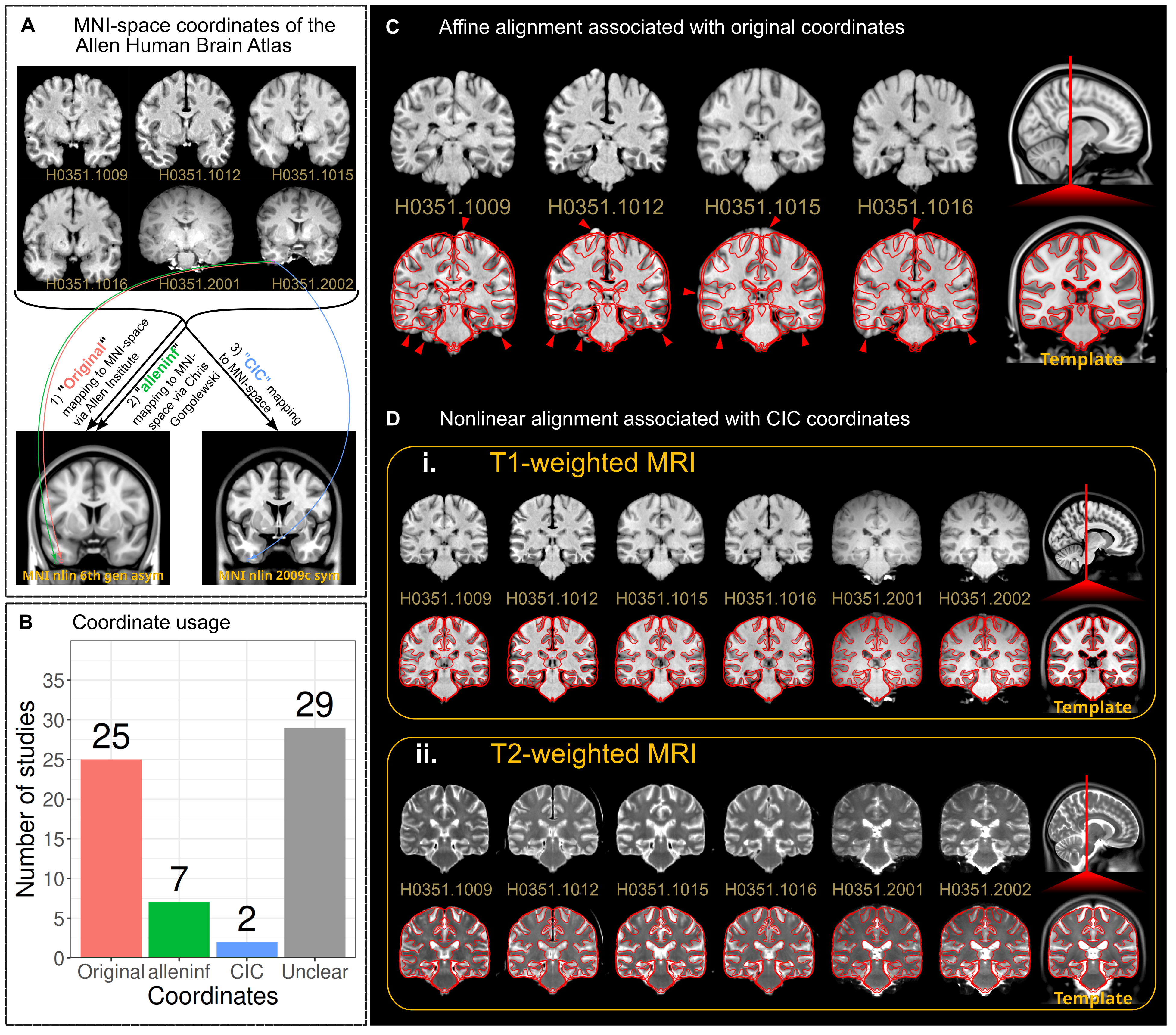

### fig2.png

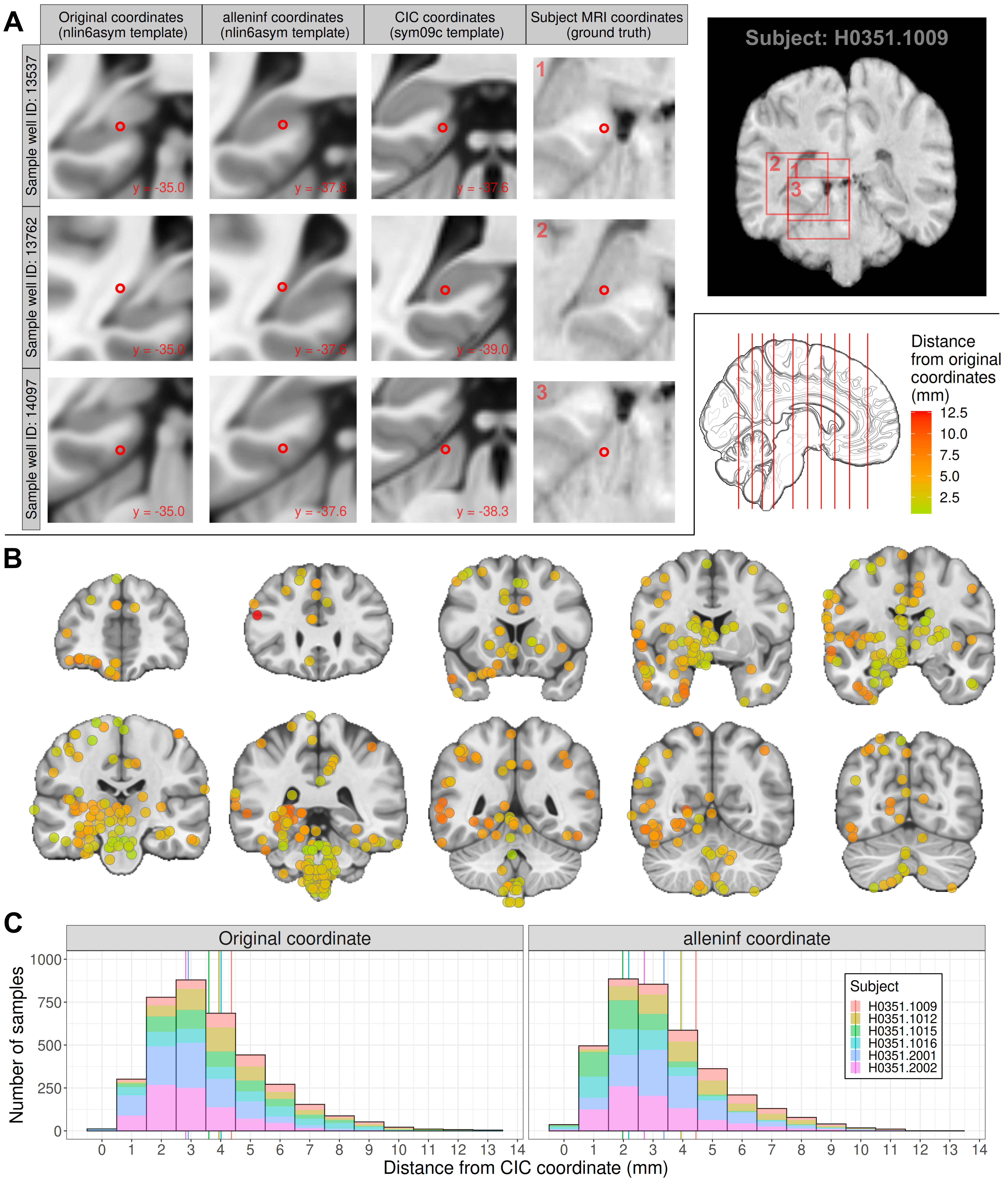

### fig3.png

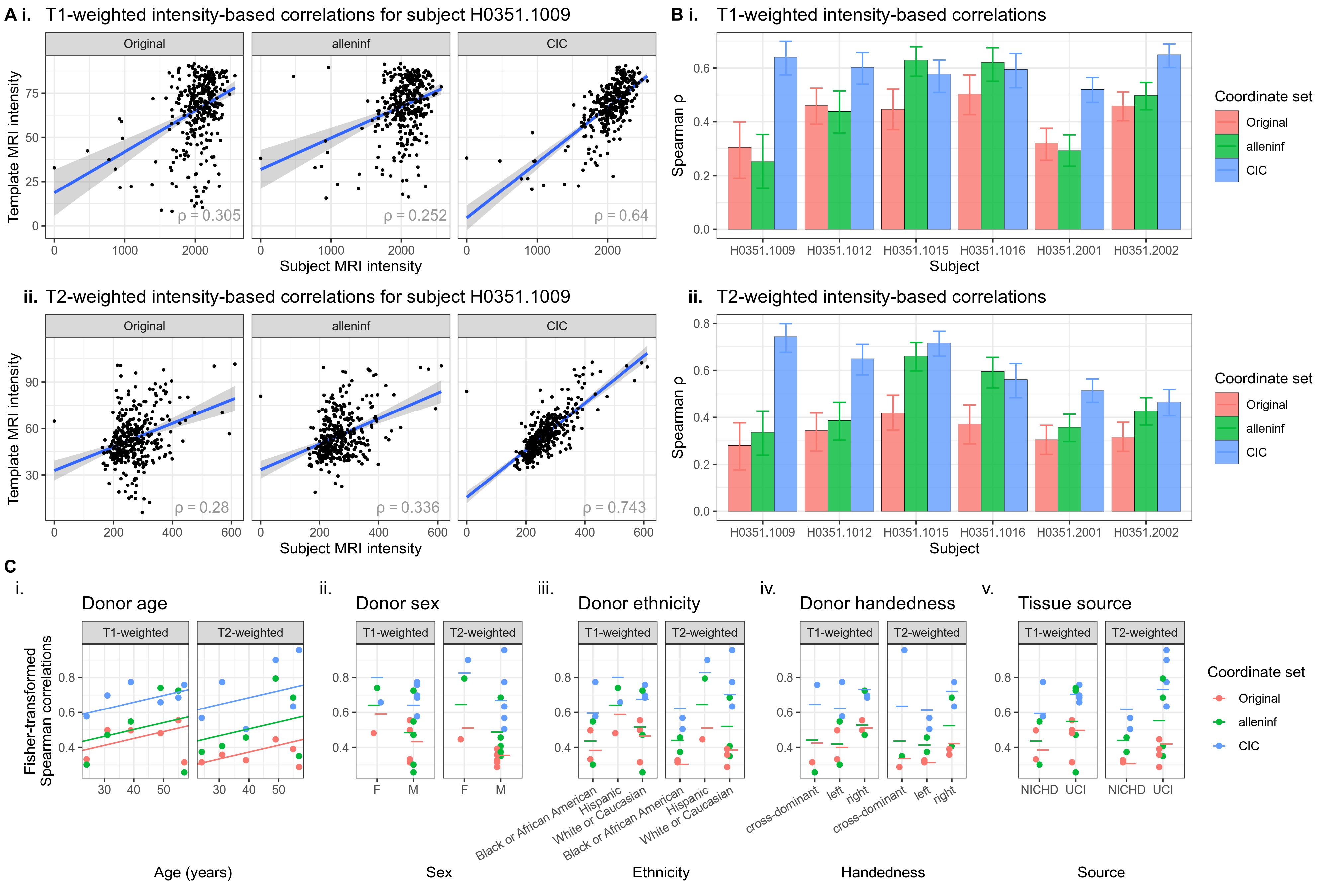

### fig4.png

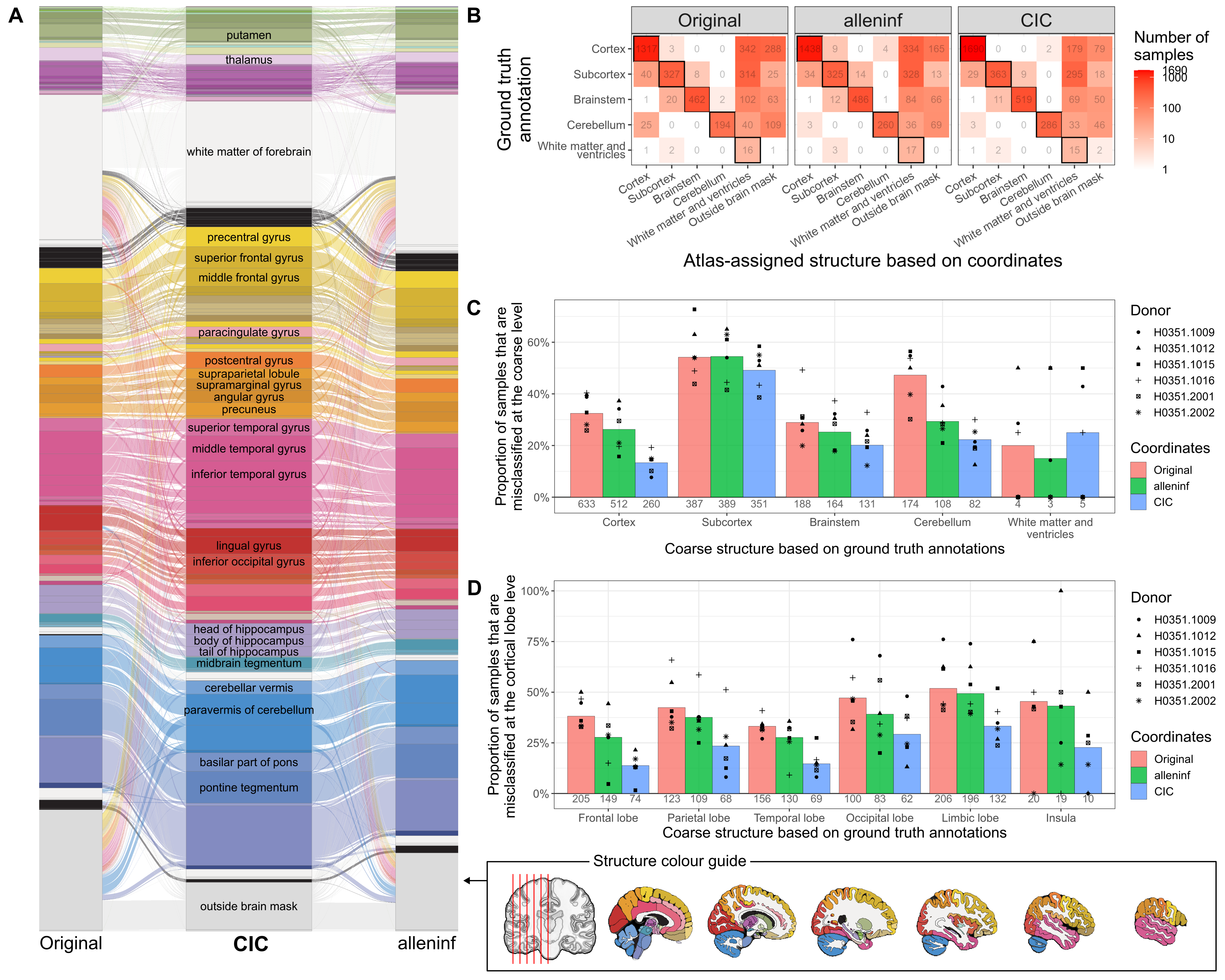

### fig5.png

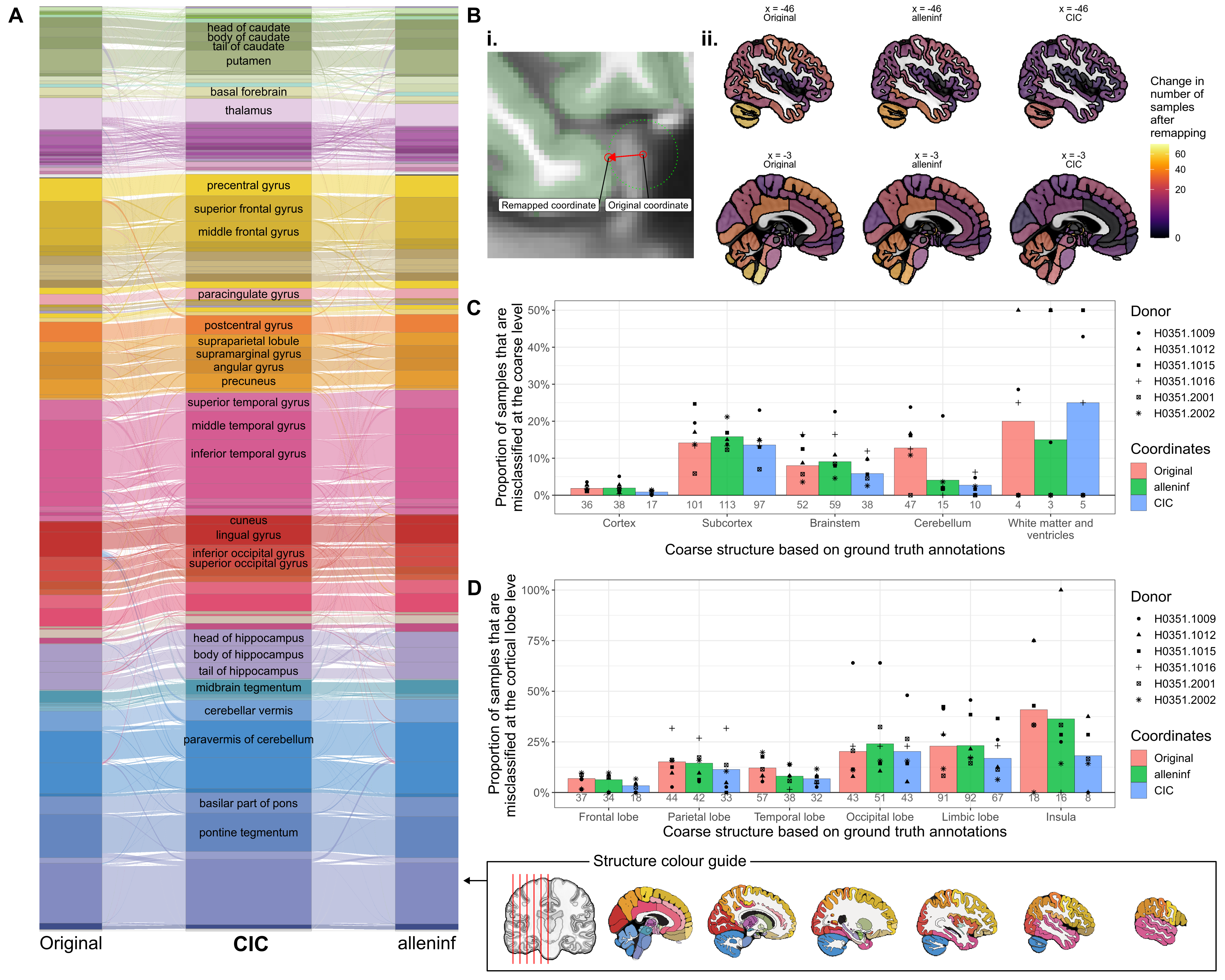

### fig6.png

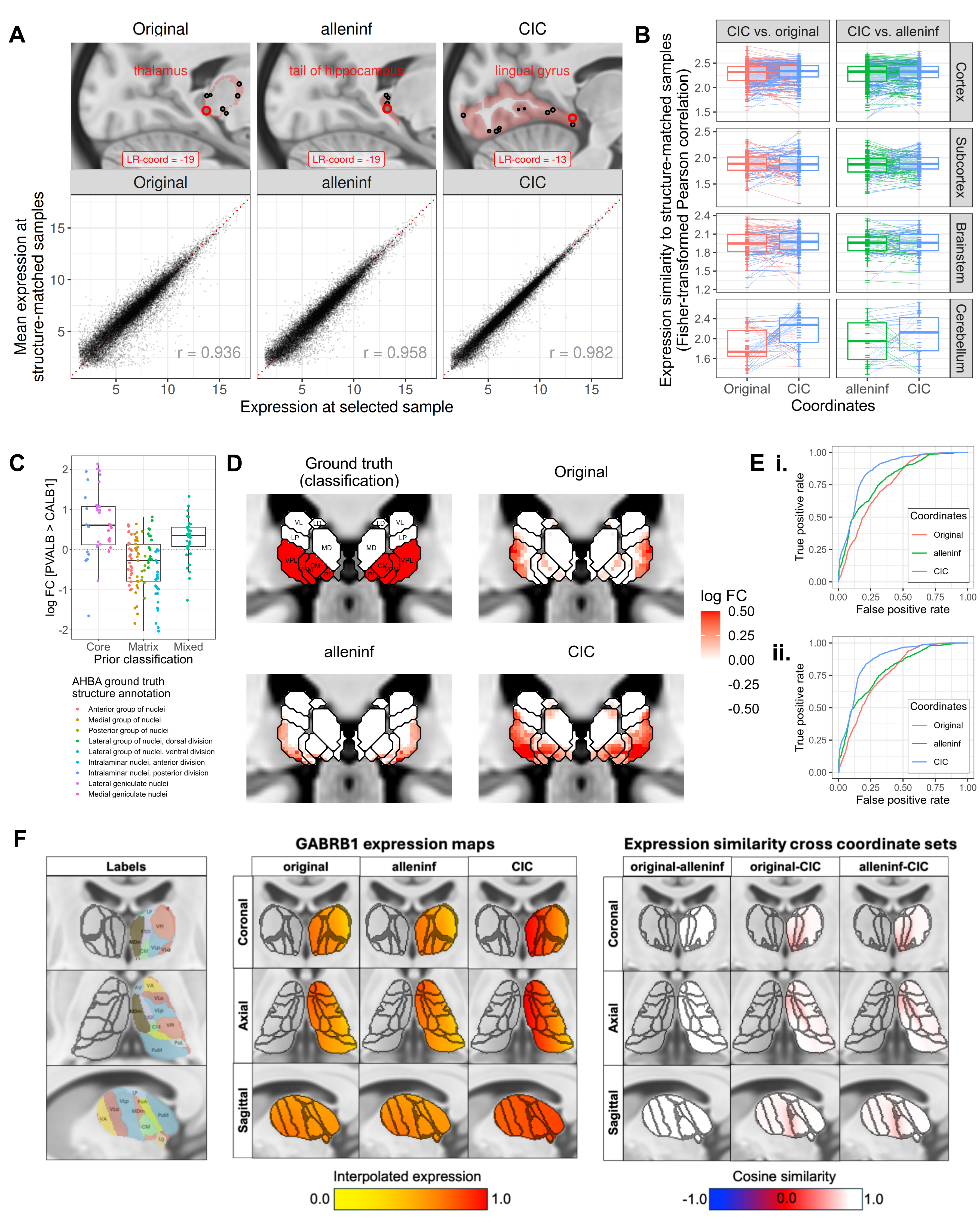

### fig7.png

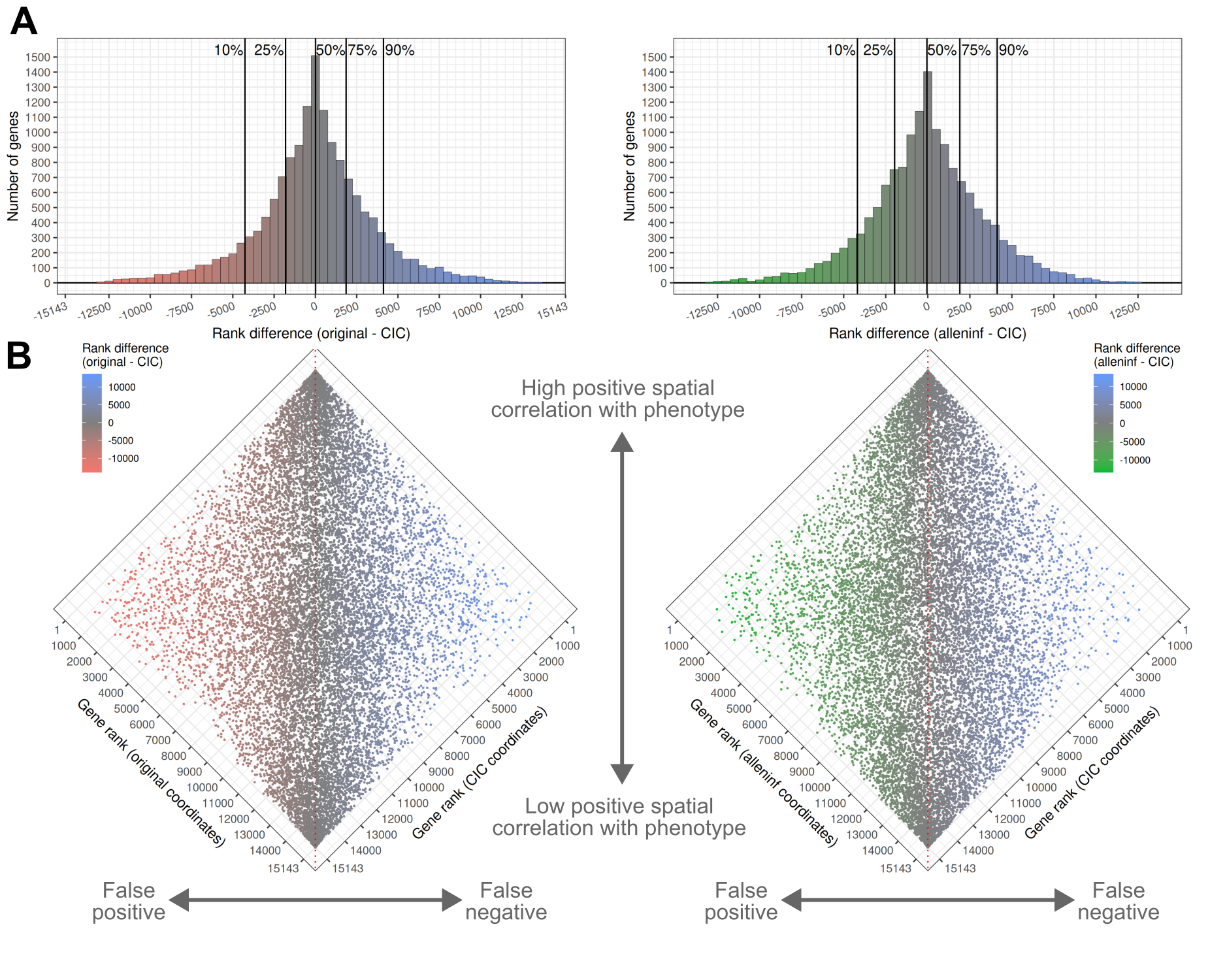

### fig8.png

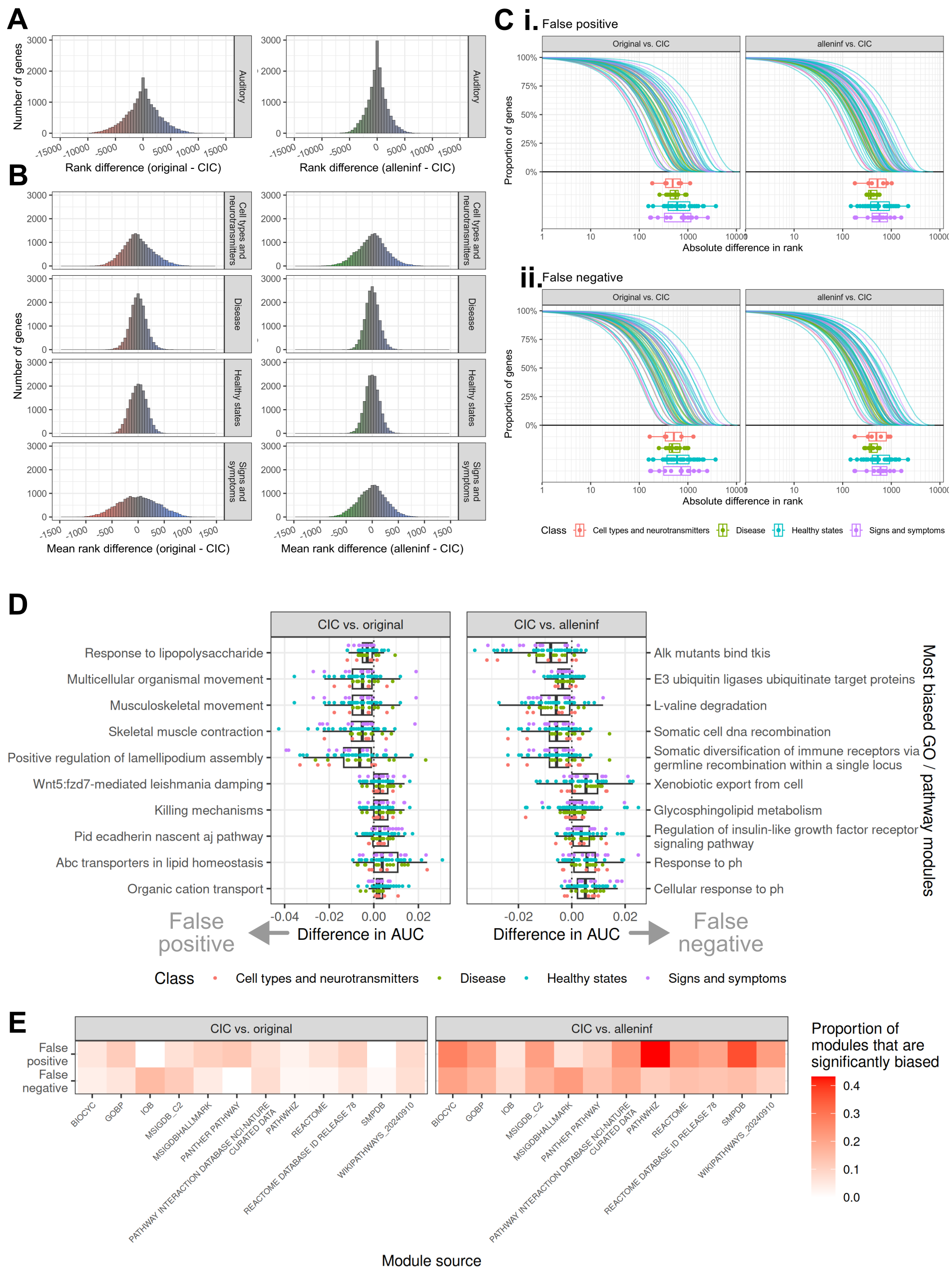

### intensity_nlin6asym.png

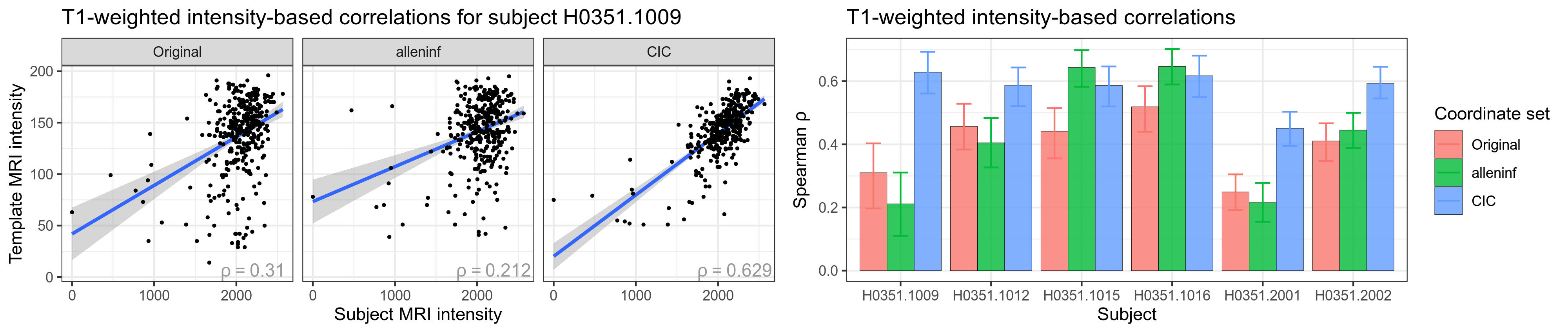

### intensity_remapped.png

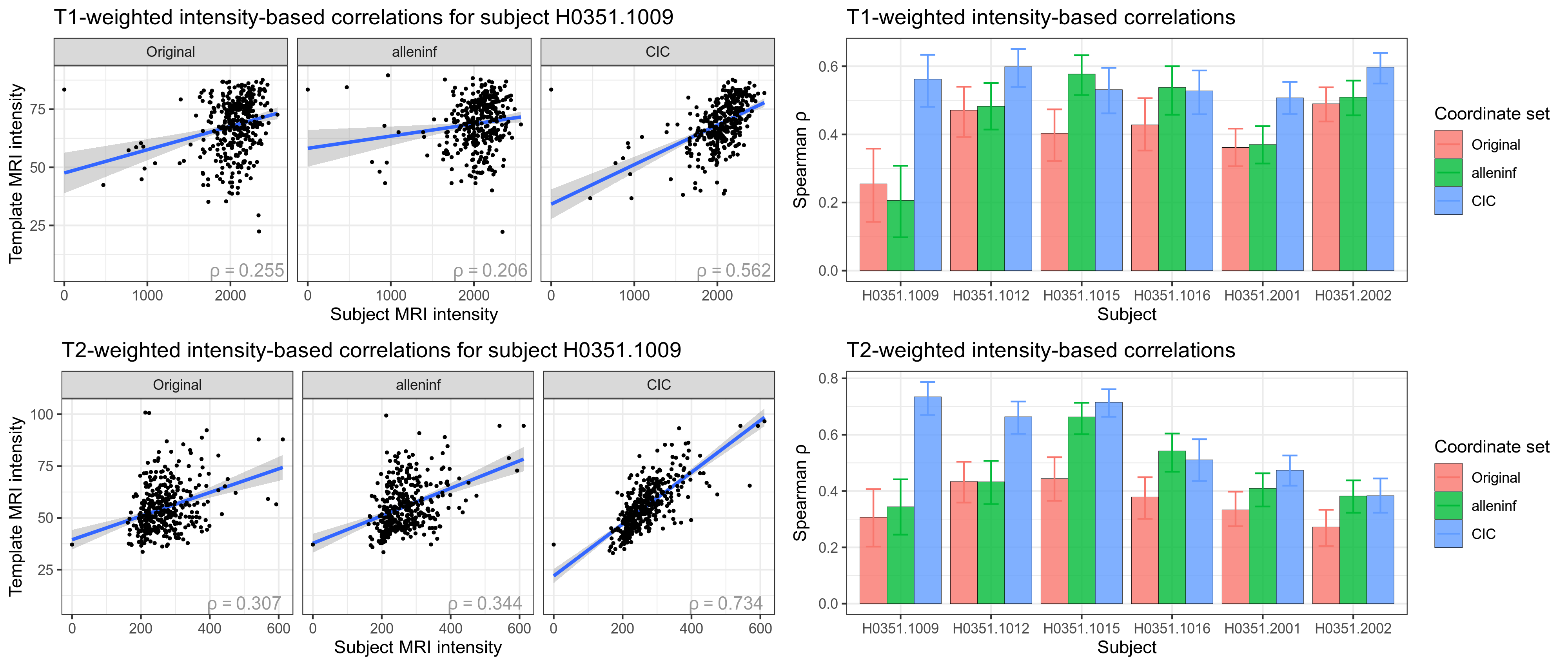

### intensity_substructures.png

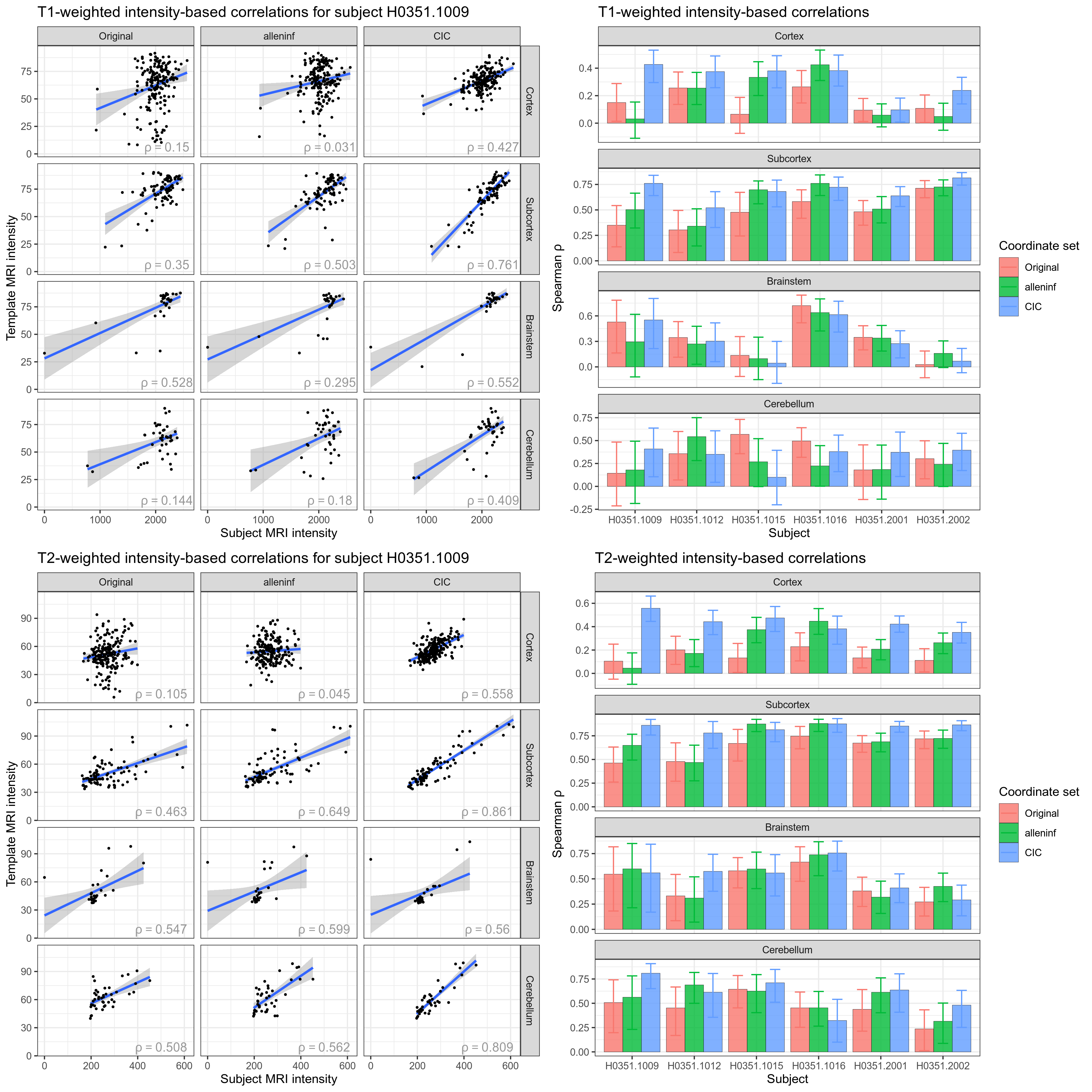

### overview_affineonly.png

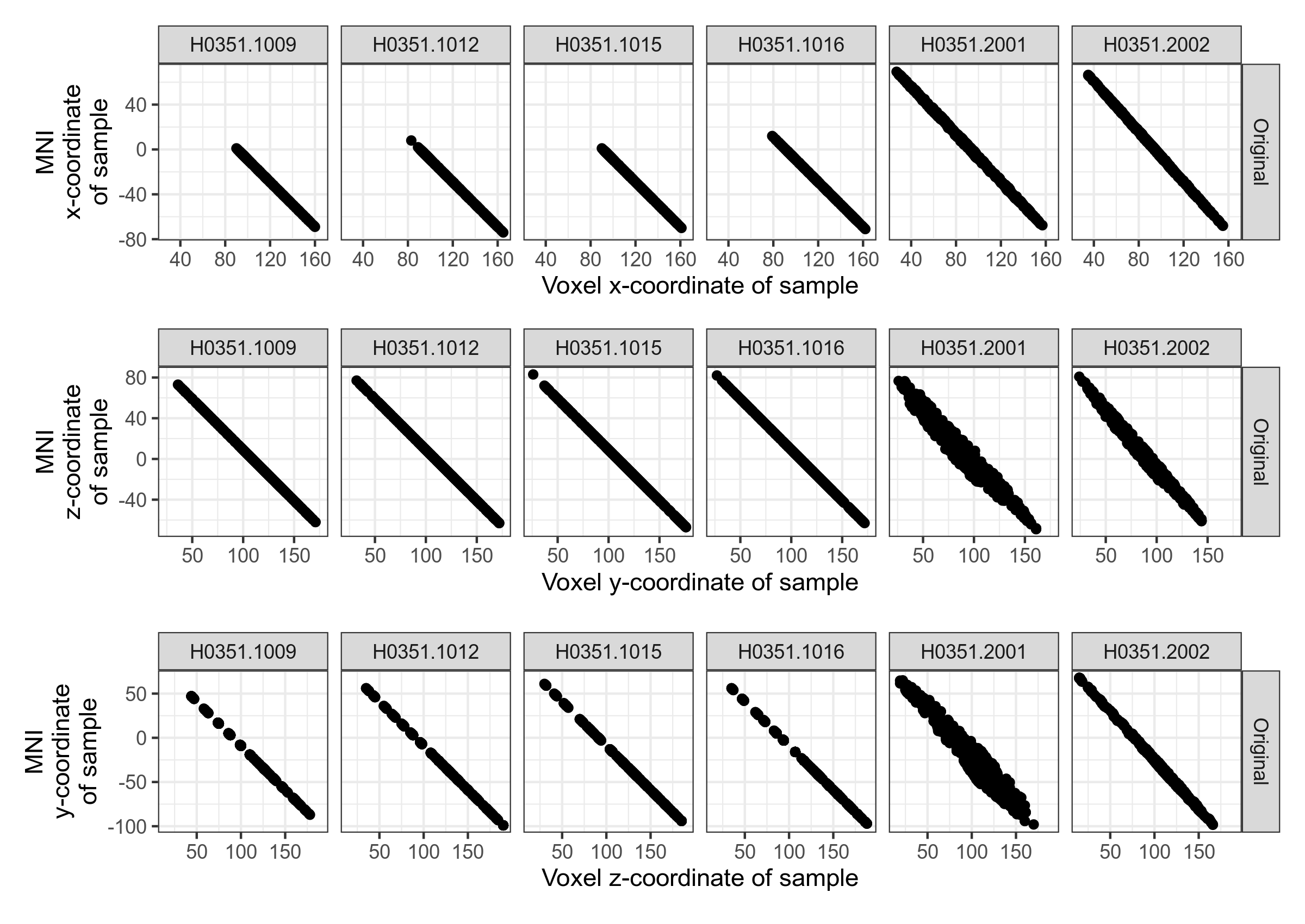

### overview_cerebellum.png

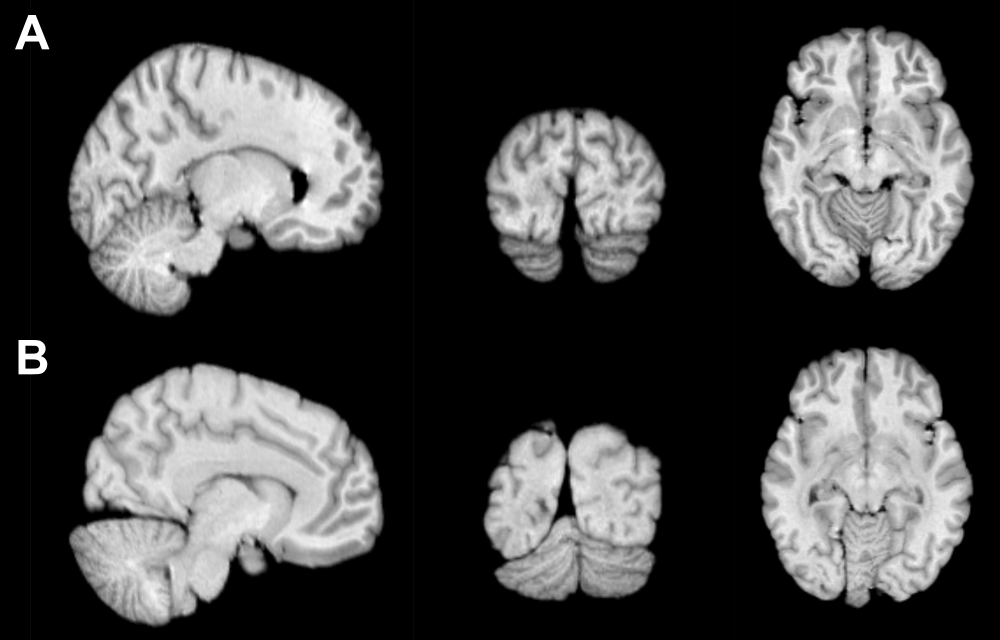

### overview_nonlinearity_index.png

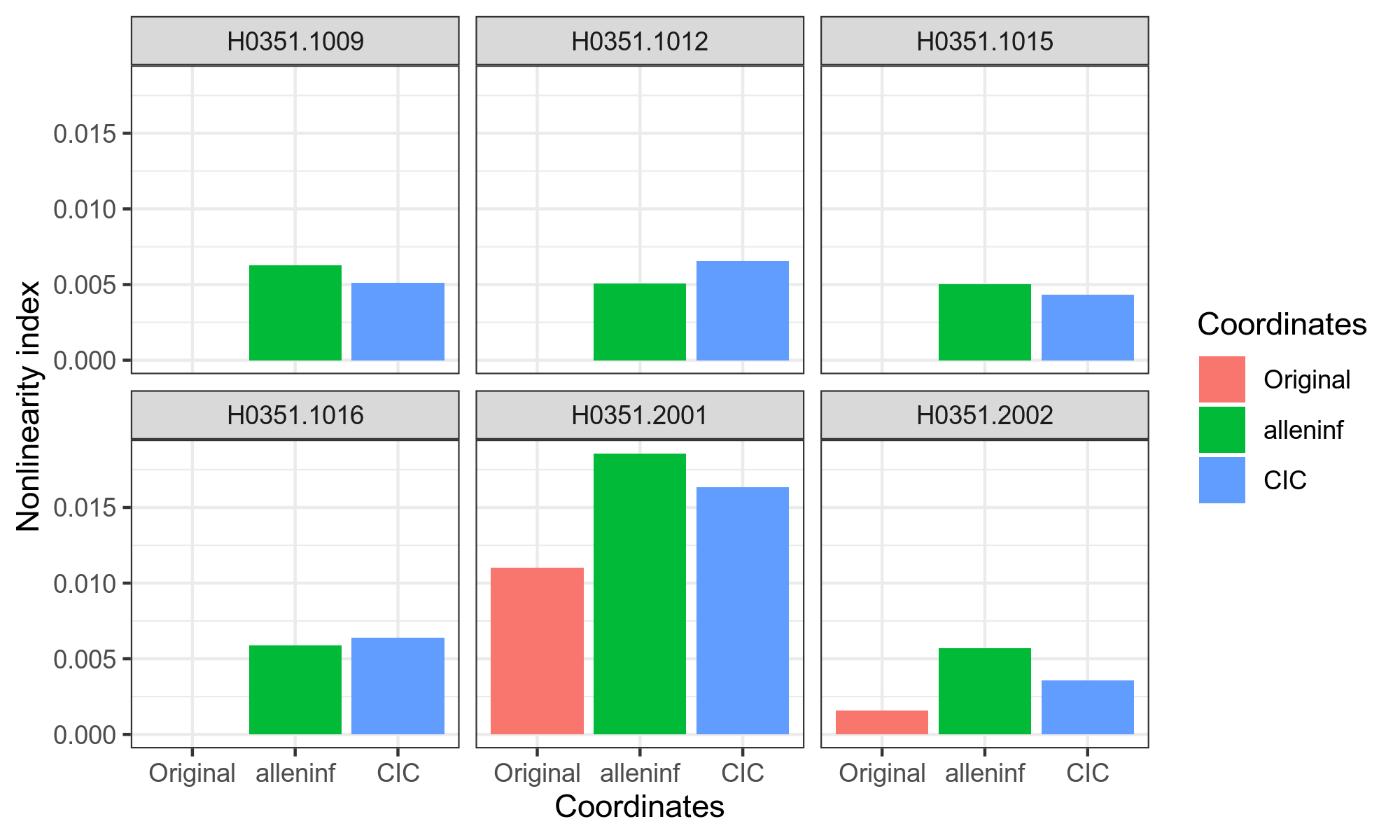

### remapping_distance_distributions.png

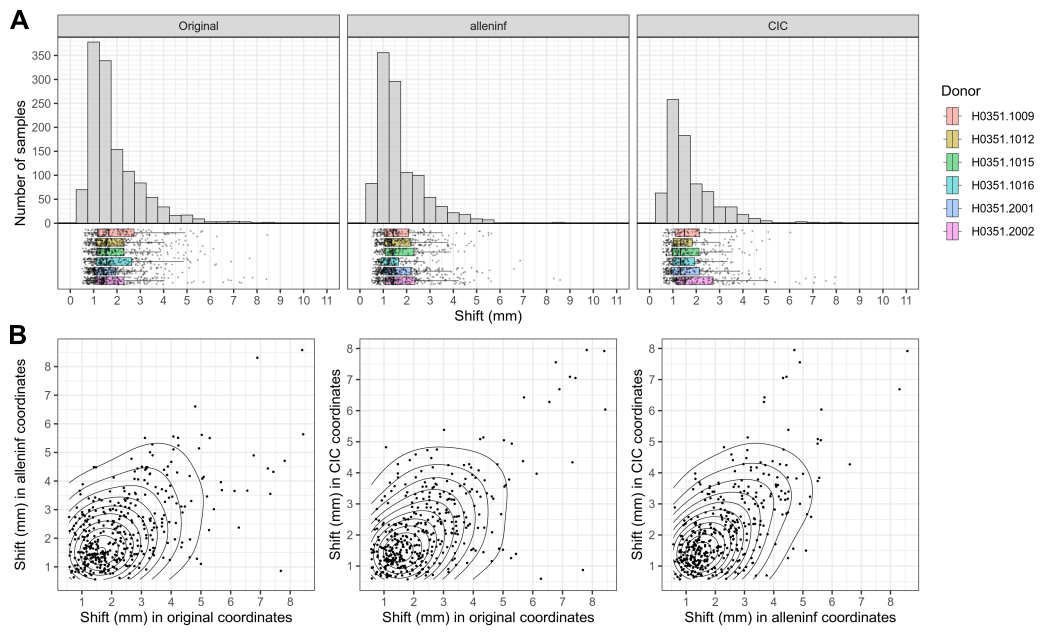

### structure_shifts_coarse_overlap_fdr_nlin6asym.png

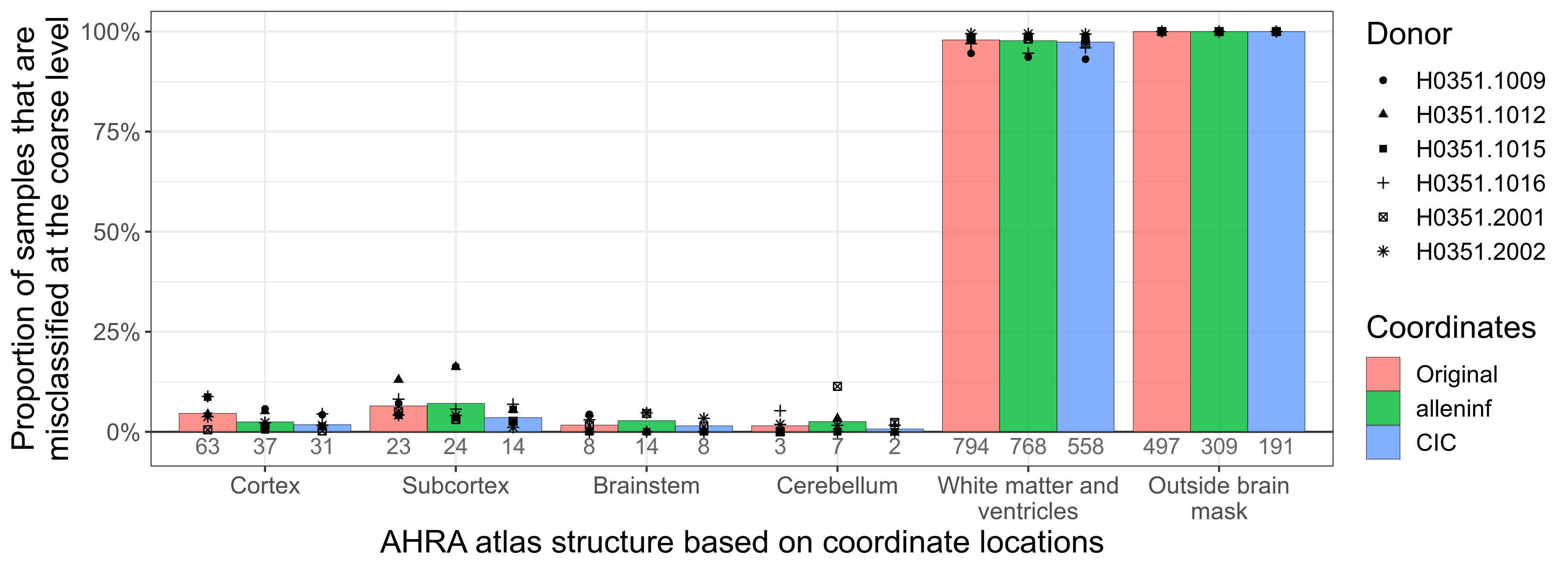

### structure_shifts_coarse_overlap_fdr_sym09c.png

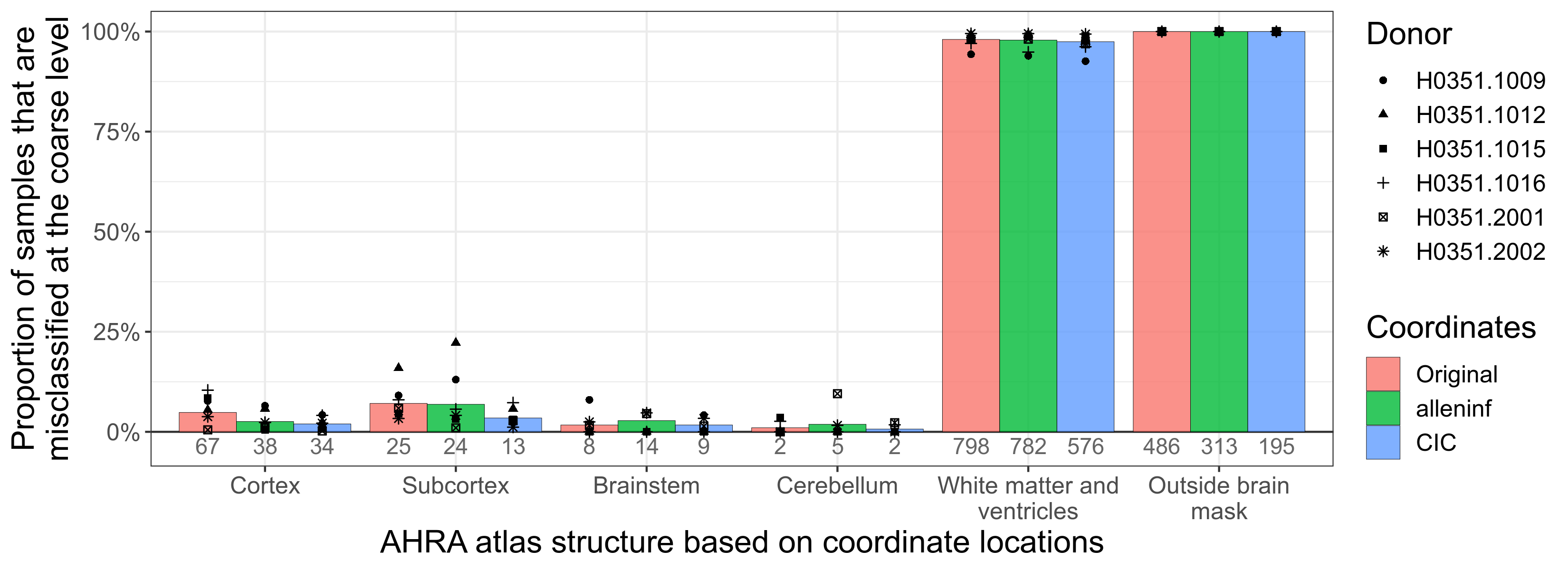

### structure_shifts_coarse_overlap_fdr_sym09c_remapped.png

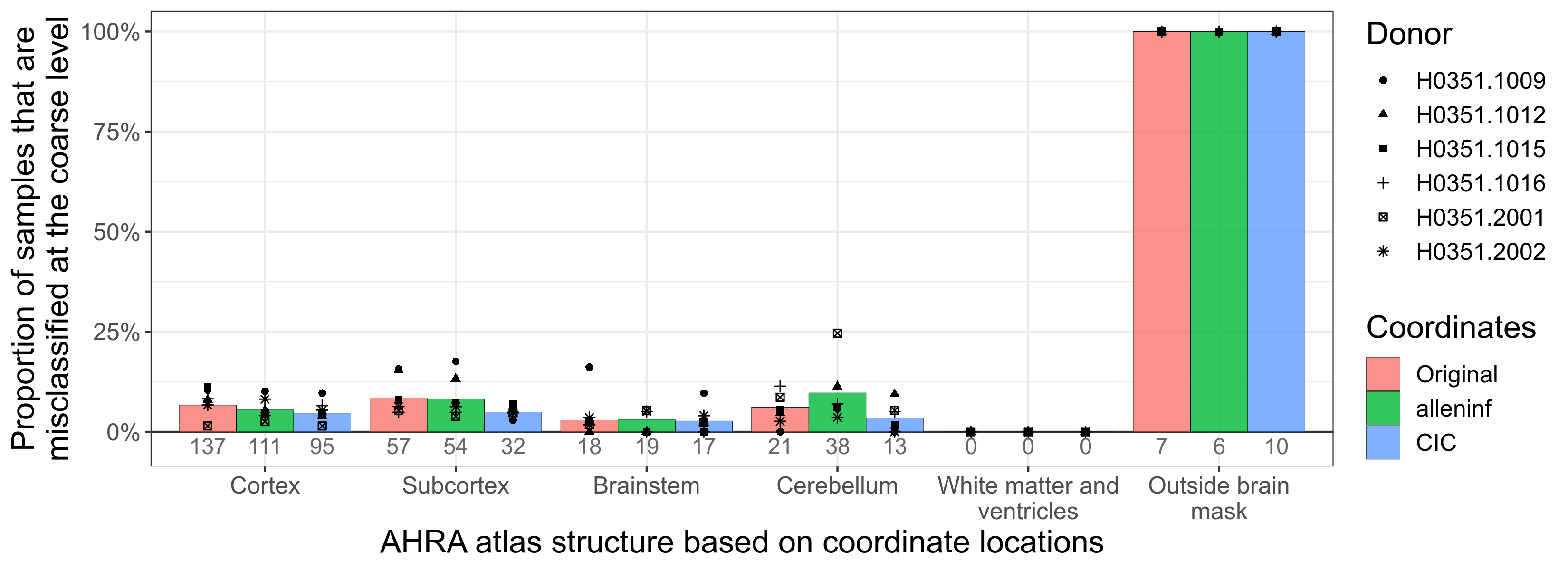

### structure_shifts_coarse_shared_structures_nlin6asym.png

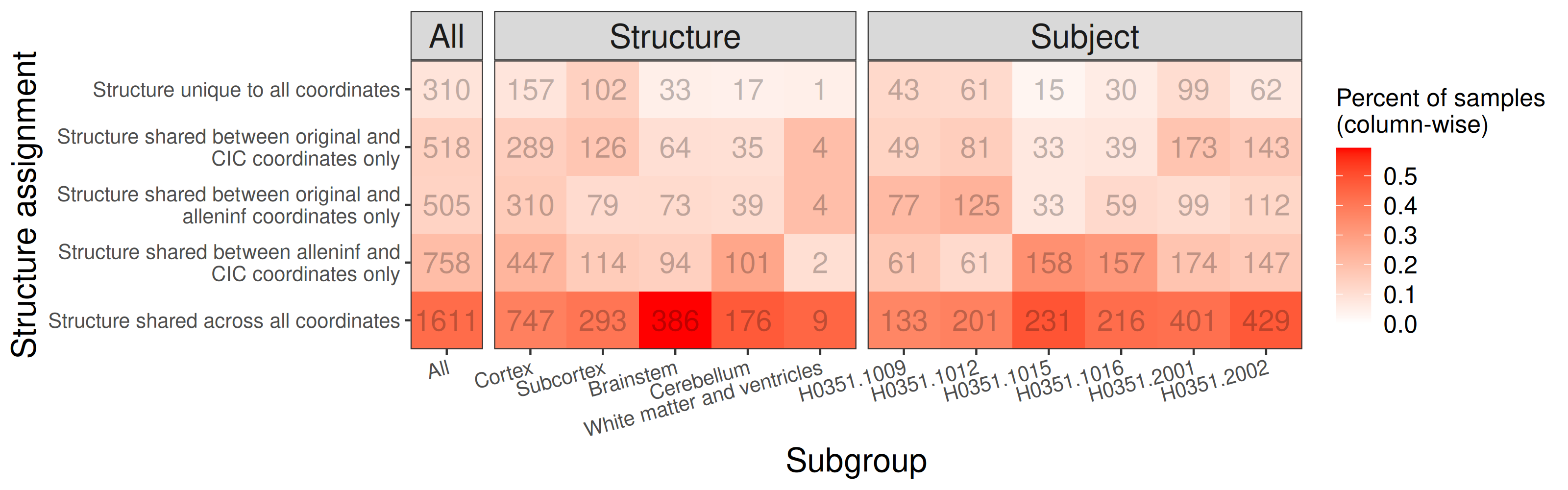

### structure_shifts_coarse_shared_structures_sym09c.png

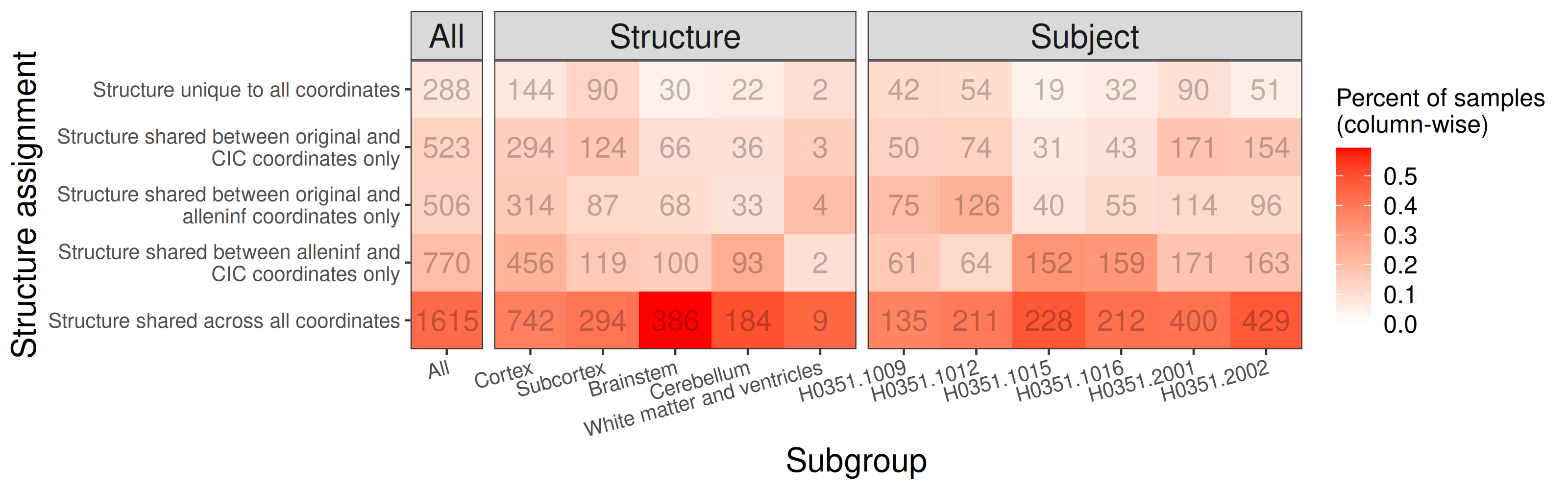

### structure_shifts_coarse_shared_structures_sym09c_remapped.png

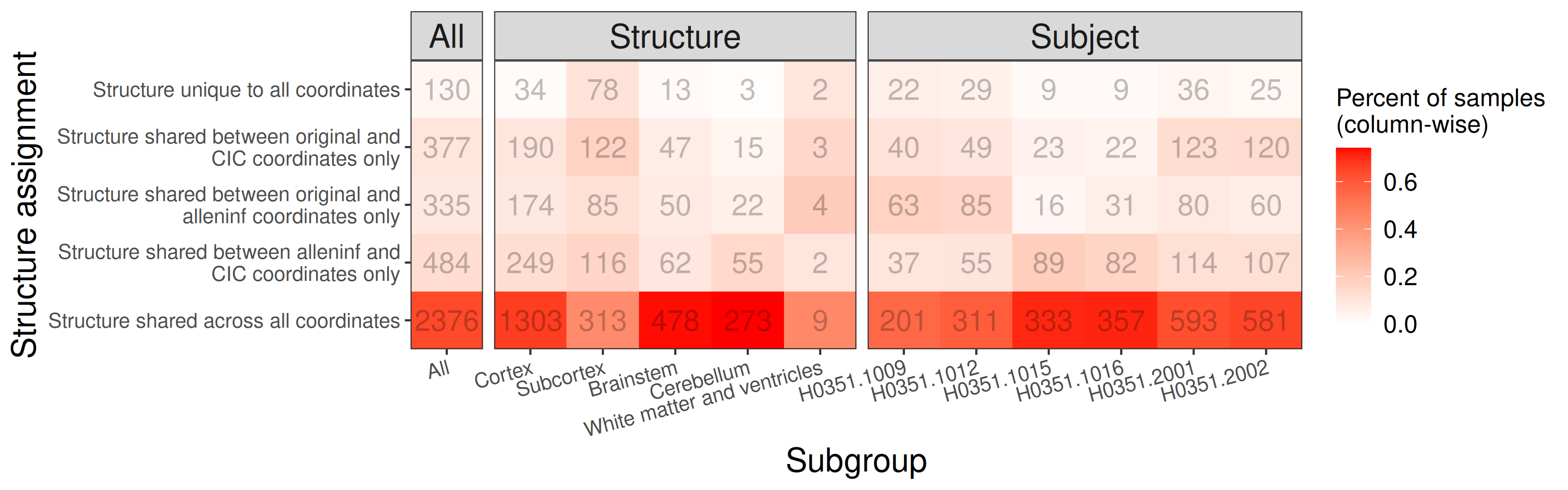

### structure_shifts_lobe_overlap_fdr_nlin6asym.png

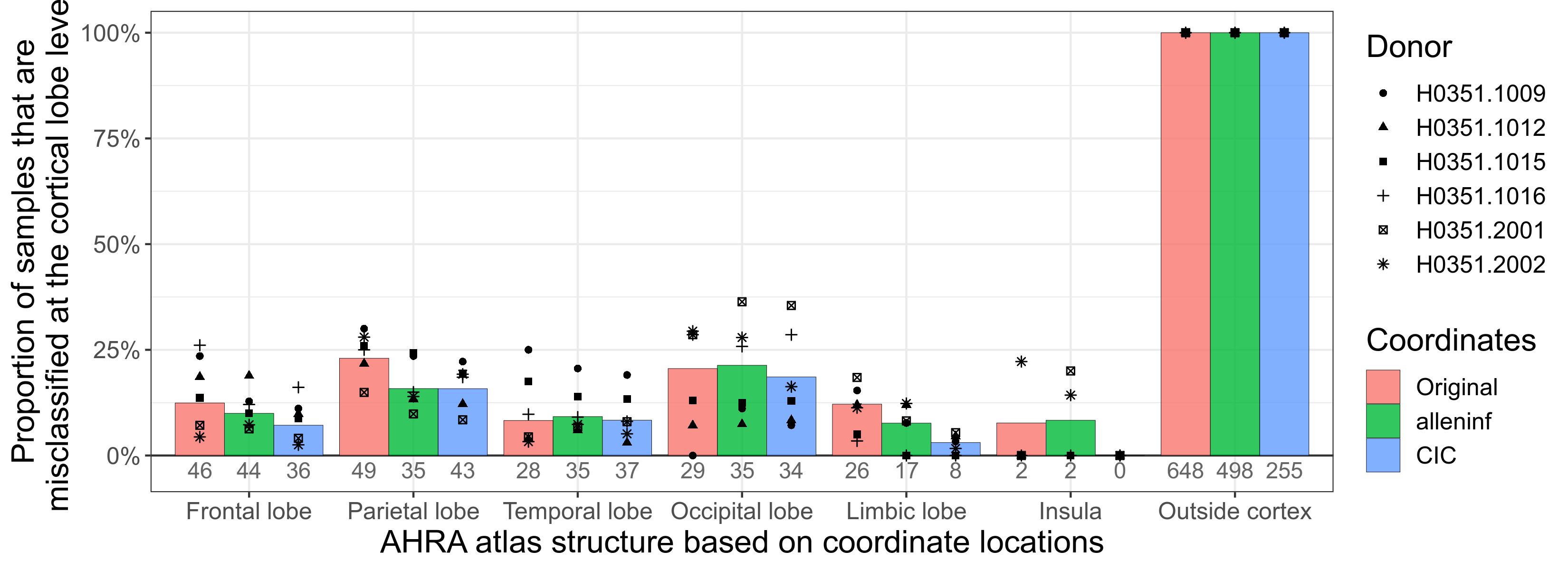

### structure_shifts_lobe_overlap_fdr_sym09c.png

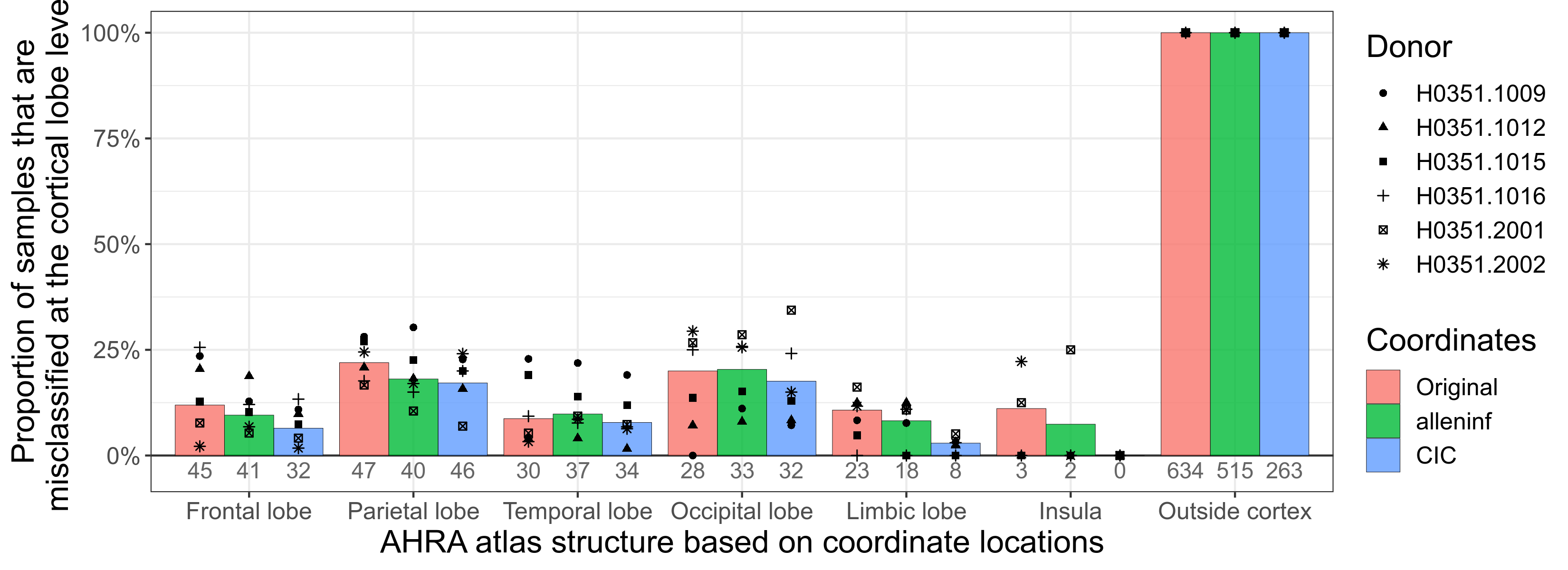

### structure_shifts_lobe_overlap_fdr_sym09c_remapped.png

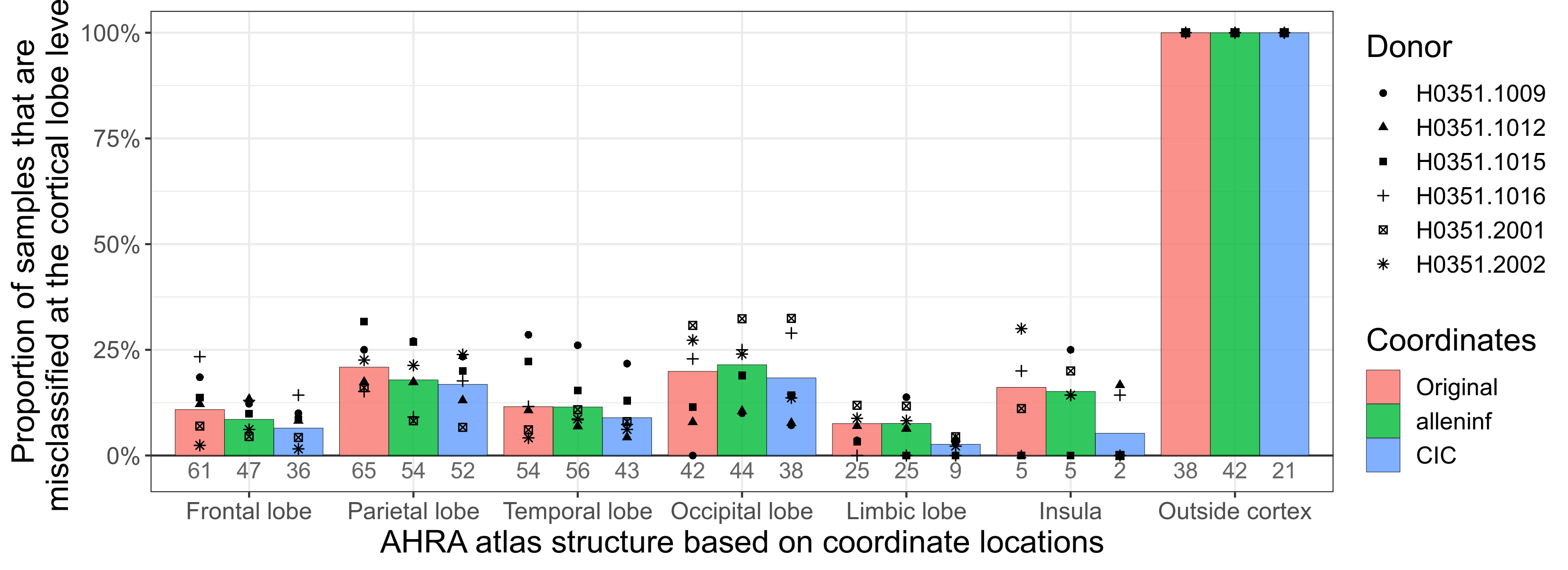

### structure_shifts_panel_nlin6asym.png

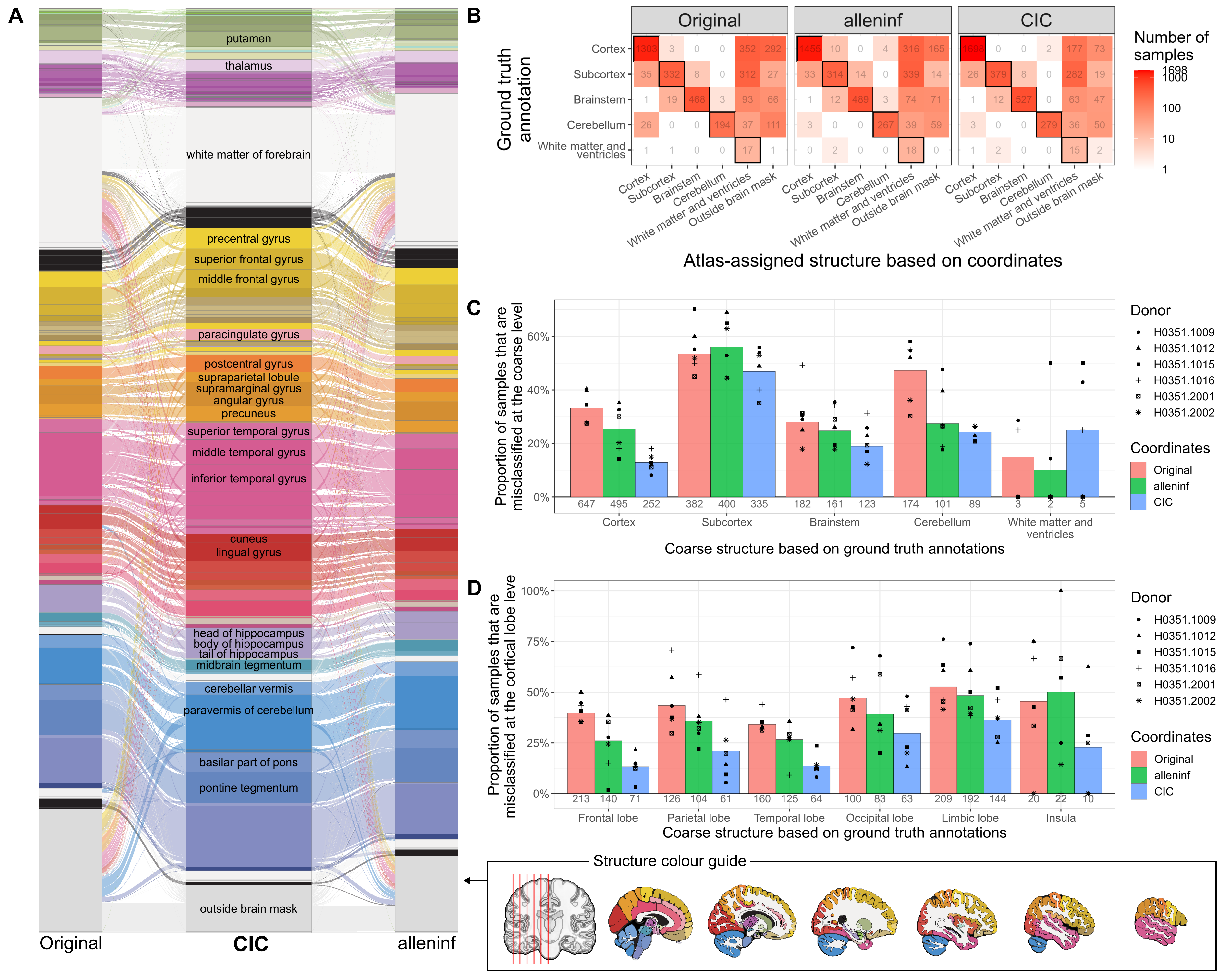

### study_selection.png

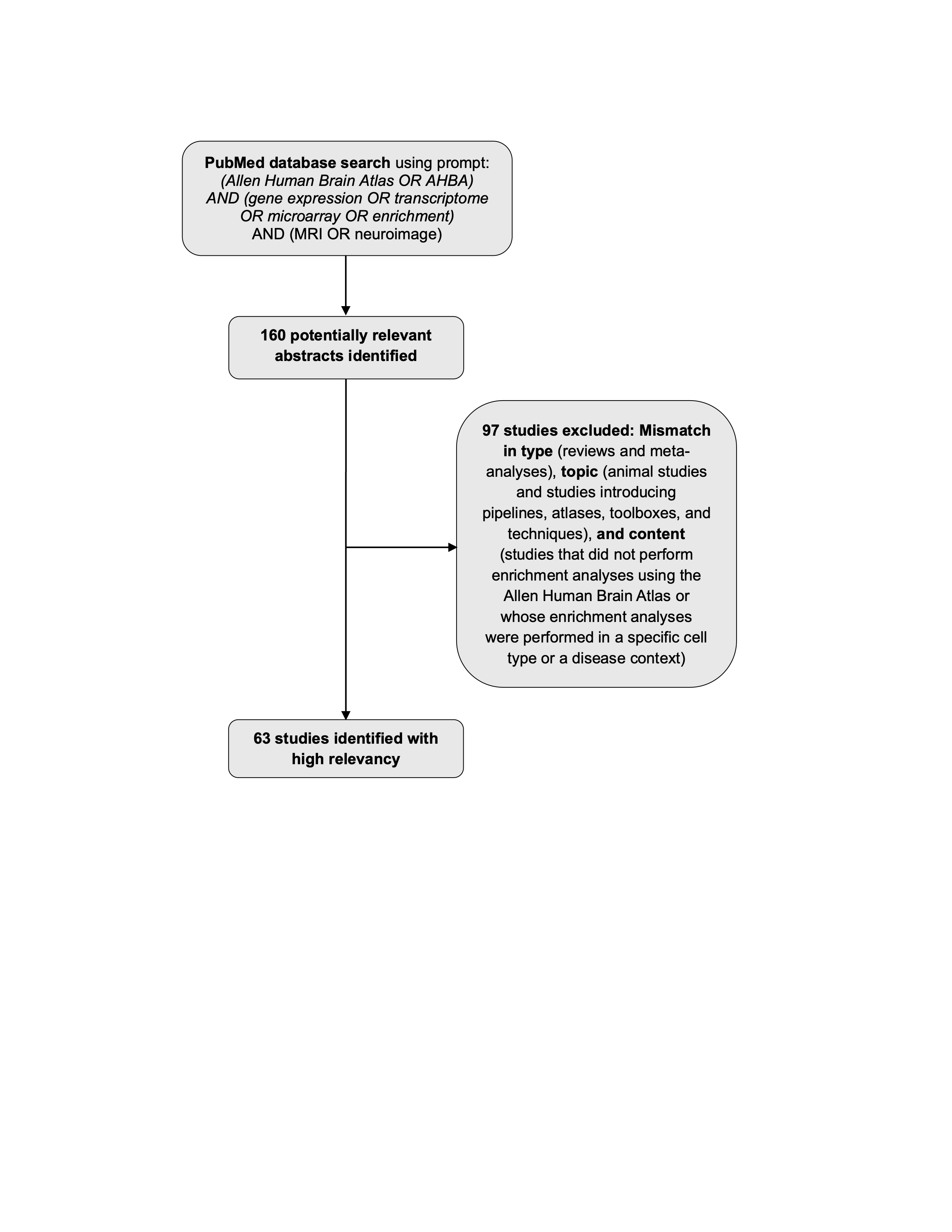
